# Characterizing lipoprotein profiles in coronary atherosclerosis development through quantitative lipidomics and proteomics approaches

**DOI:** 10.1101/2025.03.17.642310

**Authors:** Hiroaki Takeda, Yoshihiro Izumi, Yui Koike, Tomonari Koike, Kohta Nakatani, Kosuke Hata, Masaki Matsumoto, Masashi Shiomi, Takeshi Bamba

**Affiliations:** Division of Metabolomics, Medical Research Center for High Depth Omics, Medical Institute of Bioregulation, Kyushu University, 3-1-1 Maidashi, Higashi-ku, Fukuoka-shi, Fukuoka 812-8582, Japan; Department of Biotechnology and Life Science, Tokyo University of Agriculture and Technology, 2-24-16 Naka-cho, Koganei-shi, Tokyo 184-8588, Japan; Institute for Experimental Animals, Kobe University Graduate School of Medicine, 7-5-1 Kusunoki-cho, Chuo-ku, Kobe 650-0017, Japan; Department of Omics and Systems Biology, Graduate School of Medical and Dental Sciences, Niigata University, 757 Ichibancho, Asahimachi-dori, Chuo-ku, Niigata 951-8510, Japan

## Abstract

Lipoproteins, which are synthesized in hepatic cells and secreted into the blood, play an important role in lipid transport. The quality of lipoproteins, including small dense low-density lipoproteins and the cholesterol efflux capacity of high-density lipoproteins, influences cardiovascular outcomes, although the relationship between lipid changes and lipoprotein quality remains unclear. This study characterized lipid molecules and apolipoproteins in lipoprotein particles during the development of coronary atherosclerosis in myocardial infarction-prone rabbits. Using quantitative lipidomics and proteomics techniques, the lipid composition in lipoprotein particles was evaluated by normalizing lipidomics data with lipoprotein particle numbers estimated by apolipoprotein B-100 concentration. While apolipoproteins and common risk factors for abnormal lipid metabolism, including cholesterol and triacylglycerol, did not change across stages of coronary atherosclerosis, ceramide molecules in lipoprotein particles were significantly associated with early stage progression. This study provides a perspective on the role of lipid composition in the progression of coronary atherosclerosis.

## Introduction

Lipids such as cholesteryl esters (CEs) and triacylglycerols (TGs) are transported in the blood to various tissues as lipoproteins, which are covered with phospholipids and apolipoproteins^1^. An imbalance in lipoprotein levels is commonly assumed to lead to the onset of atherosclerotic cardiovascular disease. Low-density lipoprotein cholesterol (LDL-C) and high-density lipoprotein cholesterol (HDL-C) levels are commonly used diagnostic markers of abnormal lipid metabolism. Although major cardiovascular events are reduced by decreasing the LDL-C concentration with statins, they are not completely suppressed^2,3^. These cohort studies suggest that factors other than cholesterol concentration in lipoproteins can more precisely predict the severity of atherosclerotic cardiovascular disease. Recently, the quality of lipoproteins (e.g., small dense LDL and cholesterol efflux capacity of HDL) has attracted attention as a factor involved in cardiovascular events^4,5^.

Lipid composition in lipoproteins is related to their quality because lipoprotein particles consist of lipid molecules and apolipoproteins. Lipidomics, which is a comprehensive analysis of biological lipids using mass spectrometry (MS), has focused on biomarker discovery and the elucidation of lipid metabolism in serious diseases such as atherosclerosis^6^, diabetes^7^, and Alzheimer’s disease^8^. Although the lipid composition in the plasma and lipoprotein fractions has been the focus^9–12^, the alteration of lipid composition in lipoprotein particles associated with atherosclerotic cardiovascular disease is poorly understood. We have previously developed a novel lipoprotein profiling method to obtain quantitative values of lipid molecules and apolipoproteins in lipoprotein particles using lipidomics and proteomics techniques^13,14^. Single apolipoprotein B-100 (apoB-100) is present in individual very low-density lipoprotein (VLDL) and LDL particles^15,16^, and thus, the number of VLDL and LDL particles can be calculated from the apoB-100 concentration in each lipoprotein fraction determined using a targeted proteomics approach. By normalizing the lipidomics data of the lipoprotein fractions to particle numbers, the number of lipid molecules in the VLDL and LDL particles can be estimated. In addition, our methodology targets apolipoprotein C-II (apoC-II), apolipoprotein C-III (apoC-III), and apolipoprotein E (apoE), which act as activators or inhibitors of lipoprotein lipase and LDL receptor ligands. The application of our lipoprotein profiling methodology can characterize the lipid composition of lipoprotein particles associated with atherosclerotic cardiovascular disease, while monitoring apolipoproteins involved in the regulation of lipoprotein metabolism.

The myocardial infarction-prone Watanabe heritable hyperlipidemic (WHHLMI) rabbit is used as an animal model of myocardial infarction^17,18^. Lipoprotein metabolism in rabbits is similar to that in humans, but differs from that in mice and rats^19–21^. For example, mice and rats have plasma VLDL and LDL particles with apoB-48 due to the expression of the apoB mRNA editing enzyme in the liver. Their major plasma lipoprotein is HDL because of low CE transfer protein activity. In addition to lipoprotein metabolism, atherosclerosis (e.g., expression of VLDL receptors in lesions) and cardiac function (e.g., myosin type) also differ between human/rabbits and mice/rats^22^. Based on this advantage, we characterized various metabolic changes in response to pitavastatin and D-47, a novel lipid-lowering drug that reduces blood lipid concentrations by a mechanism different from that of statins^23,24^. Shiomi et al. reported that coronary atherosclerosis develops with age, and coronary lesions grow rapidly at approximately eight months^18,25,26^. Using metabolomics and lipidomics techniques, we identified candidate serum markers for the rapid progression of coronary lesions at an early stage^26^. Because the serum contains a wide variety of lipoproteins, it is difficult to explain the cause of these alterations. Focusing on individual lipoproteins may provide additional insight.

In this study, we aimed to investigate the association between lipoprotein composition, including lipids and apolipoproteins, and the severity of coronary atherosclerosis. To meet this demand, MS-based targeted quantitative lipidomics and proteomics techniques were applied to lipoprotein fractions and erythrocytes from WHHLMI rabbits. Thirteen lipid subclasses, including fatty acids (FAs), lysophosphatidylcholines (LPCs), lysophosphatidylethanolamines (LPEs), phosphatidylcholines (PCs), phosphatidylethanolamines (PEs), phosphatidylserines (PSs), phosphatidylinositols (PIs), sphingomyelins (SMs), ceramides (Cers), CEs, monoacylglycerols (MGs), diacylglycerols (DGs), and TGs, were quantified at the molecular species level, allowing the determination of individual FA compositions except for TGs, using normal-phase supercritical fluid chromatography (SFC) coupled to triple-quadrupole MS (QqQMS). In addition, apoB-100, apoC-II, apoC-III, and apoE were absolutely quantified using stable isotope-labeled peptides with nanoflow liquid chromatography (nano-LC) coupled to QqQMS. Using these advanced multi-quantitative omics techniques, we characterized the unique lipidomes of lipoprotein particles in the early stages of coronary atherosclerosis progression.

## Results

### Assessment of blood markers and pathological indicators

To characterize the compounds involved in the severity of coronary atherosclerosis, the lipid molecules and apolipoproteins of erythrocytes and/or lipoprotein fractions were analyzed in WHHLMI rabbits. The strategy used in this study is shown in Fig. 1a. Ten rabbits were divided into a severe group (*n* = 6) and a mild group (*n* = 4) based on the coronary severity score developed in our previous study (Fig. 1b and Supplementary Table 1)^26^. No significant differences in body weight were observed between the two groups (Fig. 1c). Although the aortic lesions progressed severely in both the groups, the incidence of coronary stenosis was significantly higher in the severe group (Fig. 1b). Erythrocytes and plasma were fractionated by centrifugation of blood collected from the marginal ear vein. In erythrocytes, hematocrit, an indicator of polycythemia, did not increase in the severe group (Fig. 1c). After further fractionation of lipoproteins by ultracentrifugation, total cholesterol, TG, and protein were measured using biochemical analyses (Supplementary Table 2). Cholesterol and TG concentrations in lipoproteins, which are diagnostic factors for abnormal lipid metabolism in humans, did not change significantly, except for total cholesterol at 12 months (Supplementary Fig. 1).

**Fig. 1:**
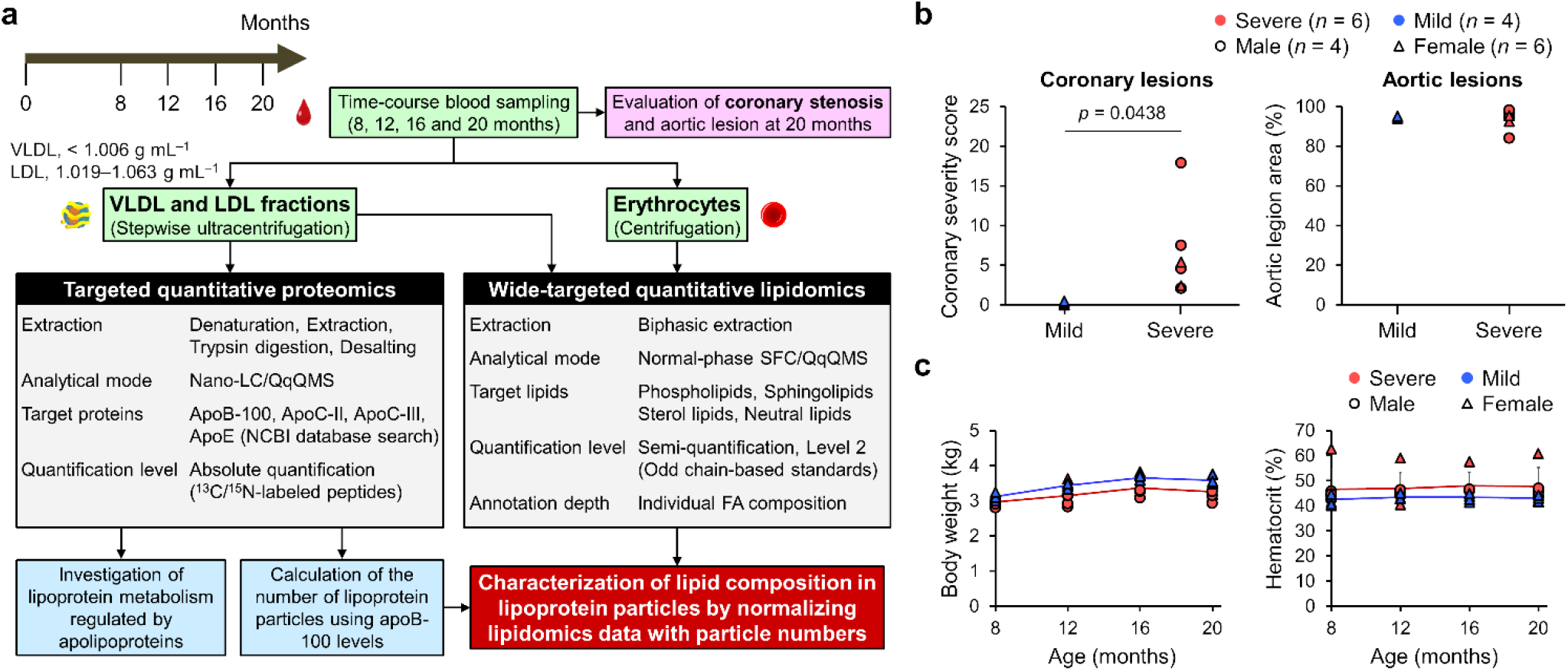
Multi-quantitative omics approach for characterizing coronary atherosclerosis development. (**a**) Strategy for characterizing lipid composition in lipoprotein particles using quantitative lipidomics and proteomics techniques. (**b**) Severity of coronary and aortic lesions in WHHLMI rabbits at 20 months. Rabbits were divided into severe and mild groups based on the coronary severity score. Circles and triangles represent males (*n* = 4) and females (*n* = 6), respectively. (**c**) Body weight and hematocrit values of WHHLMI rabbits. Red and blue indicate severe and mild groups, respectively, while circles and triangles represent males and females, respectively. Error bars represent standard deviations of biological replicates. Statistical significance was assessed using Student’s or Welch’s *t*-test following an *F*-test for variance.

### Relationship between apolipoproteins and coronary atherosclerosis

To assess the differences in lipoprotein metabolism, apolipoproteins in the VLDL and LDL fractions were quantified using nano-LC/MS-based targeted proteomics (Fig. 2a, b and Supplementary Fig. 2a). Peptide sequences of rabbit apolipoproteins were searched using the National Center for Biotechnology Information (NCBI) protein database, and multiple reaction monitoring (MRM) transitions for trypsin-digested peptides were determined using Skyline software (Supplementary Table 3)^27^. Based on the screening analyses of peptide fragments derived from apoB-100, apoC-II, apoC-III, and apoE, the targeted peptides used for quantification were selected^14^ (Fig. 2a), and their ^13^C- and ^15^N-labeled peptide internal standards were spiked into the peptide extracts (Fig. 2b). The selection criteria for the targeted peptides were as follows: i) the peptide sequences did not overlap with other sequences (not less than 10 amino acids); ii) they had three or four different product ions; iii) they represented a good peak shape and high sensitivity; and iv) they did not co-elute with other sequences. The time-course of apolipoprotein levels is shown in Fig. 2c and Supplementary Table 4. Except for the apoB-100 concentration in the LDL fraction at 20 months, the apolipoprotein concentrations (apoB-100, apoC-II, apoC-III, and apoE) were not significantly different between the severe and mild groups. Apolipoproteins were not associated with the severity of coronary atherosclerosis in WHHLMI rabbits in this study.

**Fig. 2:**
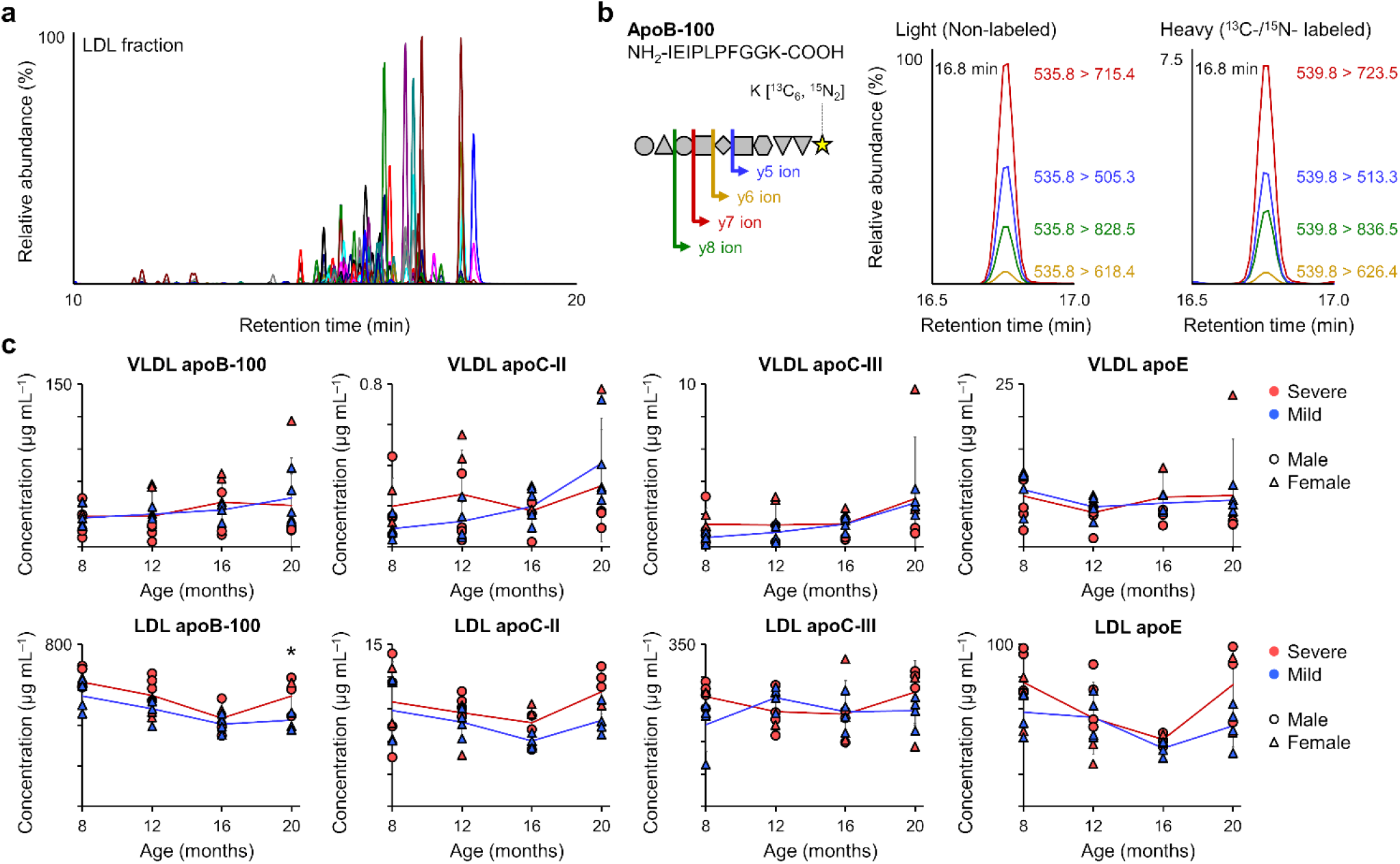
Targeted proteomics of apolipoproteins using nano-LC/MS. (**a**) Total ion chromatogram of LDL fractions using nano-LC/MS. (**b**) Quantification of apolipoproteins using stable-isotope-labeled synthetic standards. (**c**) Time-course changes in apolipoproteins within VLDL and LDL fractions. Red and blue indicate severe and mild groups, respectively, while circles and triangles represent males and females, respectively. Error bars indicate standard deviations of biological replicates. Statistical significance was assessed using Student’s or Welch’s *t*-test following an *F*-test for variance (^*^*p* < 0.05).

### Correlation between apoB-100 and lipid subclasses

Since apolipoproteins did not change significantly, other components, such as lipid molecules, seemed to be involved in lipoprotein quality. We further examined the differences in lipoprotein quality between the mild and severe groups. When the serum lipidomics data from the previous study^26^ were reanalyzed, the sex difference was not shown by principal component analysis (PCA) (Fig. 3a). The lipid molecules in each sample were annotated using in-house lipid-screening methods developed in our previous study^28^. Our in-house lipid screening methods contain approximately 2500 lipid molecules, including 22 lipid subclasses with a variety of 23 FA side chains. Based on the lipid screening of erythrocyte, serum, and lipoprotein fractions, 196, 232, and 237 molecules were detected in each reference sample containing equal amounts of lipid extracts, respectively. Lipid subclasses were highly separated within 20 minutes using the normal phase mode (Fig. 3b). Therefore, these constituent lipid molecules were quantified using a dodecanoyl- or heptadecanoyl-based synthetic internal standard mixture (Supplementary Fig. 2b and Supplementary Tables 5–8), and the concentrations of the lipid subclasses were calculated by summing the levels of the lipid molecules. Since a single apoB-100, a non-exchangeable apolipoprotein, is contained in single VLDL and LDL particles, it reflects the number of lipoprotein particles^15,16^. The correlation between apoB-100 and each lipid subclass in the lipoprotein fractions was evaluated each month using Pearson’s correlation coefficient to characterize lipoprotein quality, regardless of the severity of coronary atherosclerosis (Fig. 3c, d). Although the concentrations of apoB-100 and most lipid subclasses were significantly correlated each month in the VLDL fractions, the number of lipid subclasses that correlated with apoB-100 was reduced in the LDL fractions.

**Fig. 3:**
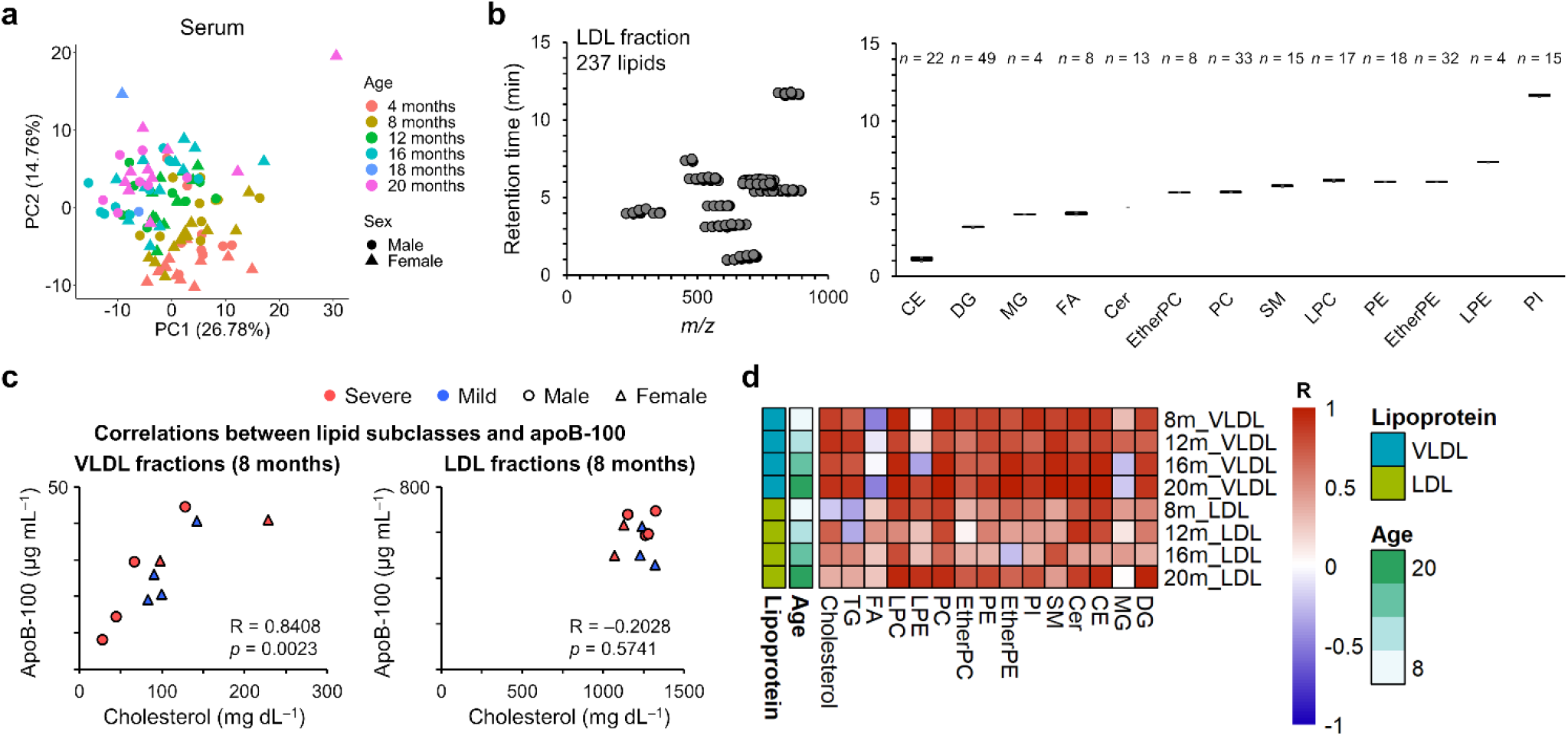
Targeted lipidomics of lipoprotein lipids using SFC/MS. (**a**) Serum lipid profiles visualized to assess age, sex, and disease severity. PCA was performed on the autoscaled data (mean-centered and variance-scaled) to assess the variance explained by each principal component. Lipidomics data were reanalyzed from a previous study^26^. (**b**) Separation of lipid subclasses in LDL fractions using normal-phase SFC/MS. The number of lipids detected is shown. (**c**) Correlation between apoB-100 and cholesterol levels in VLDL and LDL fractions at 8 months. Red and blue represent severe and mild groups, respectively, while circles and triangles represent males and females, respectively. Correlation analysis was conducted using Pearson’s method. (**d**) Correlation between apoB-100 and lipid subclasses across at 8, 12, 16, and 20 months in VLDL and LDL fractions.

### Time-course alteration of lipid subclasses

We characterized the lipid subclasses in erythrocytes and lipoprotein particles involved in coronary atherosclerosis. The number of VLDL and LDL particles was estimated from the apoB-100 concentration analyzed by nano-LC/MS. The lipidomics data of the erythrocytes and lipoprotein fractions were normalized to the number of red blood cells and lipoproteins, respectively (Fig. 4a). A comparison of the lipid subclasses between the two groups is shown in Supplementary Fig. 3. A decrease in serum ether PC and an increase in serum Cer levels were observed in the severe group at eight months (Fig. 4b, c). Focusing on lipoprotein particles, Cer in the VLDL and LDL particles increased significantly in the severe group at eight months (Fig. 4b, c). The change in serum Cer at eight months was the result of increased Cer concentrations in VLDL and LDL particles. In erythrocytes, Cer did not change significantly from month to month, but PI decreased significantly at 16 months (Fig. 4b, c).

**Fig. 4:**
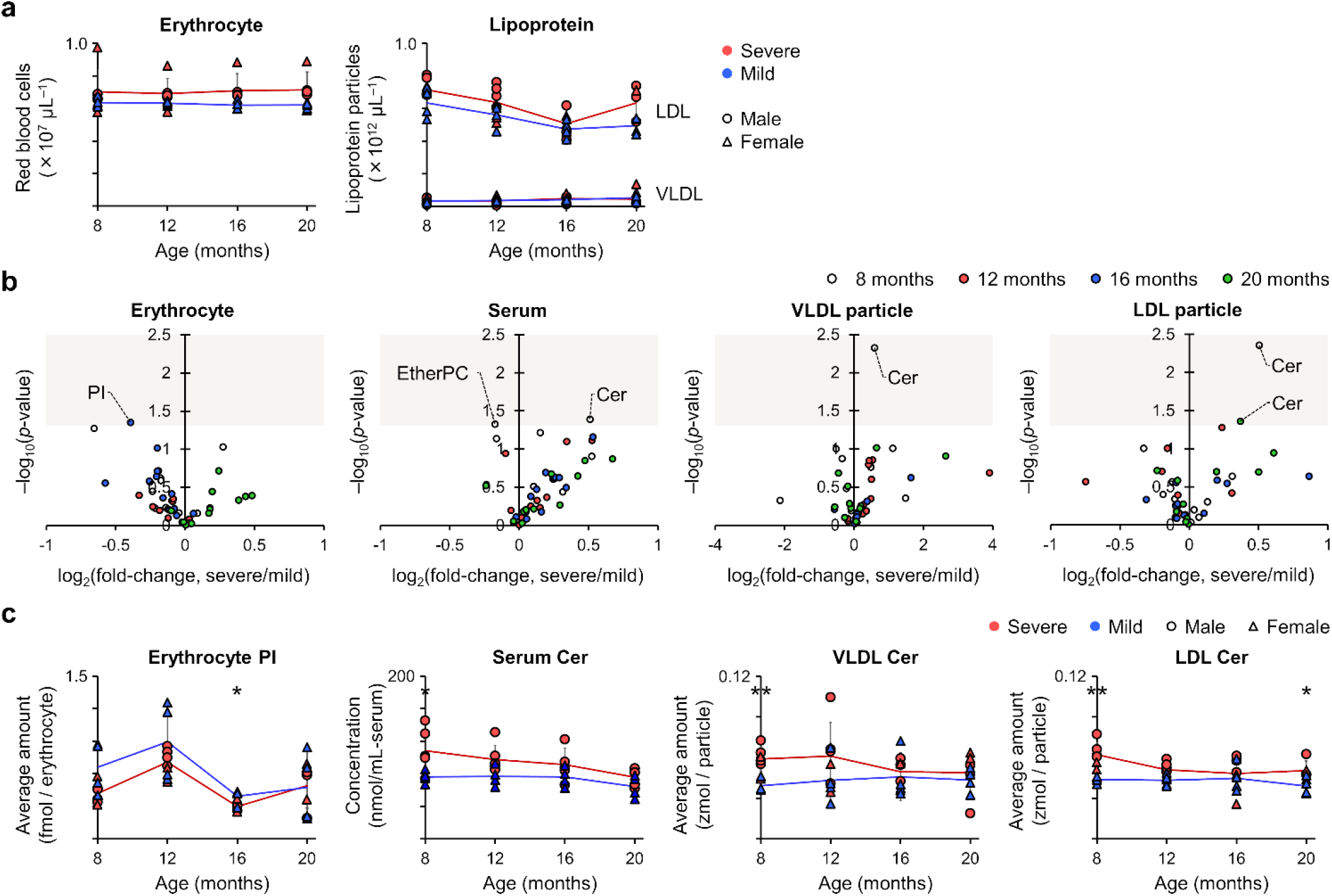
Time-course of lipid subclasses in erythrocytes and lipoprotein particles. (**a**) Number of erythrocytes and lipoprotein particles in blood. ApoB-100 molecules in 20 μL of VLDL and LDL fractions were calculated using Avogadro’s constant. Red and blue represent severe and mild groups, respectively, while circles and triangles represent males and females, respectively. Error bars indicate standard deviations of biological replicates. Statistical significance was assessed using Student’s or Welch’s *t*-test following an *F*-test for variance. These values were used for normalization of the lipidomics data. (**b**) Comparison of lipid subclasses between severe and mild groups using a volcano plot. Beige areas highlight significantly different lipid subclasses (*p* < 0.05). (**c**) Changes in representative lipid subclasses between severe and mild groups across ages. Red and blue indicate severe and mild groups, respectively, while circles and triangles represent males and females, respectively. Error bars represent standard deviations of biological replicates. Statistical significance was determined using Student’s or Welch’s *t*-test followed by *F*-test for variance (^*^*p* < 0.05 and ^**^*p* < 0.01). A complete time-course analysis of lipid classes is shown in Supplementary Fig. 3.

### Unraveling lipid dynamics by profiling lipoproteins and erythrocytes in coronary atherosclerosis

We further examined the constituent lipid molecules in the erythrocytes and lipoprotein particles on a monthly basis to gain specific insights (Supplementary Fig. 4). Similar to the behavior of the lipid subclasses, serum lipid profiling at eight months characterized 18 molecules that differed significantly between the severe and mild groups (*p* < 0.05) (Fig. 5a, b). Notably, nine Cer molecules, representing more than half of the detected Cer molecules, were significantly increased in the severe group (Fig. 5b). In lipoproteins, 8 and 14 lipid molecules in the VLDL and LDL particles, respectively, also exhibited significant changes (Fig. 5a, b). Among these, five Cer molecules in the LDL particle were increased in the severe group compared to the mild group, similar to the result of serum lipid profiling (Fig. 5b, c). In addition, two Cer molecules and total Cer increased in the VLDL particle (Fig. 5b, 5c). The number of Cer molecules that significantly changed ranked first or second among all lipid subclasses, with Cers in the serum and LDL particles showing the highest ratio of changes relative to the total number of lipids detected (Fig. 5b). In particular, saturated Cer molecules such as Cer 18:1;O2/18:0 and Cer 18:1;O2/22:0 have been characterized in both serum and lipoprotein particles during the early stages of atherosclerosis (Fig. 5c).

**Fig. 5:**
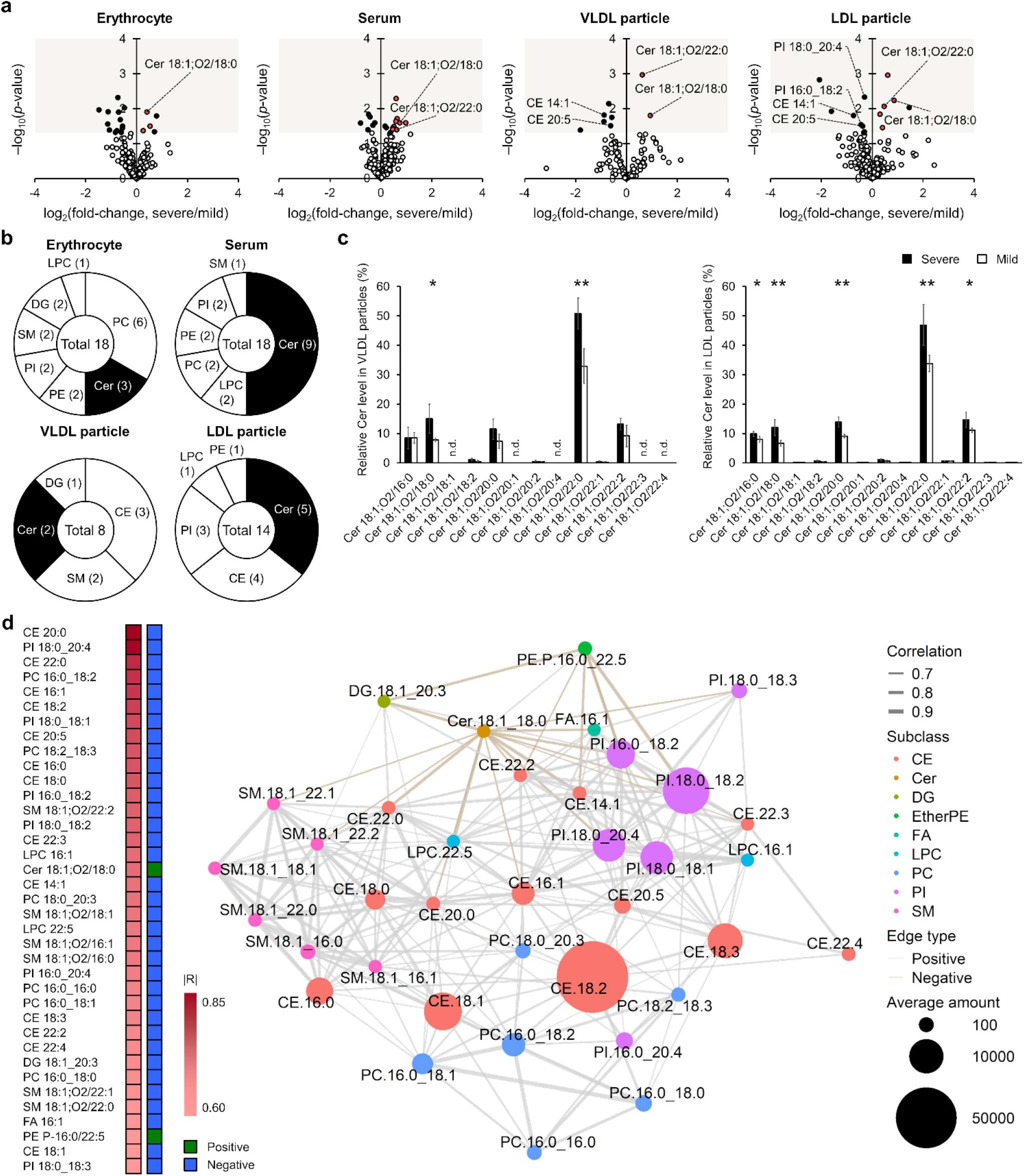
Increased Cers in lipoprotein particles with coronary lesion progression at 8 months. (**a**) Lipid profiling of erythrocytes, serum, and lipoprotein particles at 8 months using a volcano plot. Black circles represent all lipids that showed significant changes (*p* < 0.05), while red circles indicate significantly altered Cer molecules. A complete profile of lipid molecules is shown in Supplementary Fig. 4. (**b**) Number of significantly altered lipid molecules in erythrocytes, serum, and lipoprotein particles. Black areas represent Cer molecules and the numbers in brackets indicate the number of altered lipids. (**c**) Comparison of Cer molecules in VLDL and LDL particles between severe (*n* = 6) and mild (*n* = 4) groups. Black and white bars represent the severe and mild groups, respectively. Error bars indicate standard deviations of biological replicates. Statistical significance was assessed using Student’s or Welch’s *t*-test after *F*-test for variance (^*^ *p* < 0.05, ^**^ *p* < 0.01). (**d**) Correlation between coronary severity scores and individual lipid molecules in LDL particles at 8 months. Pearson’s correlation analysis was performed, focusing only on lipids detected in all samples. Lipids with significant correlations (|R| > 0.6) are displayed. Positively and negatively correlated lipids are shown in green and blue, respectively. The network of lipid molecules correlated with the coronary severity score (|R| > 0.6) was constructed based on pairwise correlation coefficients (|R| > 0.6). Nodes represented individual lipid molecules, with node size corresponding to the average lipid abundance. The color of the edges was assigned based on the direction of the correlation, with positive correlations in gray and negative correlations in beige.

To explore lipid dynamics from another perspective, we performed Pearson correlation analysis between lipid levels and the absolute coronary severity score (Fig. 5d). This revealed that while phospholipids and CEs showed significant negative correlations with the severity score, Cer 18:1;O2/18:0 was uniquely and significantly positively correlated. In addition, the network analysis highlighted the proximity of lipid subclasses, suggesting that lipids within the same subclass tend to behave similarly in the progression of coronary atherosclerosis (Fig. 5d). The clustering of lipids further highlighted the distinct and potentially subclass-specific roles in atherosclerosis, providing new insights into lipid dynamics in coronary disease.

While the significance of Cers in serum and lipoproteins was most apparent at eight months, other lipid subclasses and time points also showed notable changes. For example, CE molecules changed significantly in VLDL and LDL particles at eight months (Fig. 5b), while PC molecules in erythrocytes showed the highest number of alterations among all subclasses (Fig. 5b). At 16 months, erythrocyte lipids appeared to be affected by the progression of coronary atherosclerosis, with PE molecules emerging as prominent markers, as suggested by seven significantly altered PE and ether PE molecules (Supplementary Fig. 4). Early changes in Cer molecules were observed in VLDL and erythrocytes, but these changes were less consistent across time points (Supplementary Fig. 4). In serum and LDL particles, some Cer molecules showed significant changes throughout the study period, although the number of altered lipids was significantly reduced compared to eight months (Supplementary Fig. 4). These findings highlight the unique lipid dynamics across compartments and time points, and further emphasize the critical change in Cers at eight months in the context of atherosclerosis progression.

## Discussion

The WHHLMI rabbit is an animal model of coronary atherosclerosis^17^. The severity of atherosclerotic lesions in the coronary arteries varied between rabbits, although atherosclerotic lesions in the aorta developed severely in each rabbit (Supplementary Table 1). Since conventional factors (e.g., glucose, insulin, oxidized LDL, superoxide dismutase activity, and NADPH oxidase) did not change significantly between the severe and mild groups^26^, it is suggested that other factors are involved in the severity of coronary atherosclerosis. The concentration of lipid molecules in Japanese white rabbits is extremely low compared to that in WHHLMI rabbits^13,26^. In fact, many lipid molecules were below the limit of detection in the Japanese white rabbits. Therefore, we investigated the correlation between blood lipid molecules and the progression of coronary atherosclerosis in WHHLMI rabbits.

We have previously developed a methodology for lipoprotein profiling using targeted lipidomics and proteomics approaches (Fig. 1a)^14^. The targeted lipidomics system has been described in a previous study^28^. Briefly, each lipid subclass was separated using SFC with a diethylamine-bonded silica column, because its stationary phase recognizes the head groups rather than the FA side chains of lipid molecules. Lipid molecules were measured using the MRM mode of QqQMS. By applying fatty acyl-based MRM transitions in negative ion mode, lipid molecules, including structural isomers [e.g., PC 16:0_20:4 (840.6 > 255.3 and 840.6 > 303.2) and PC 18:2_18:2 (840.6 > 279.3)] can be distinguished even when co-eluted by SFC. Although the target molecules must be determined in advance when using the MRM mode, qualitative information can be obtained by screening lipid molecules in a reference sample using in-house lipid libraries (approximately 2500 compounds). Lipid molecules can be analyzed within a reasonable quantification range of 64.9% to 103.5% by adding an internal standard of each lipid subclass to normalize the corresponding ionization efficiency^28^. In addition, our previous study indicated that each lipid subclass could be quantified within a plausible value by summing the constituent lipids of the same subclass^29^. A targeted proteomics system was also described in a previous study^14^. Based on the complete amino acid sequences of apolipoproteins obtained from the NCBI protein database, trypsin-digested peptides of apolipoproteins were determined using Skyline software^27^. To quantify proteins using the nano-LC-MS system, it is necessary to prepare isotopomers of the target peptides to normalize for matrix effects caused by co-eluted components. The number of lipoprotein particles can be calculated using the Avogadro constant because a single apoB-100 molecule is present in individual VLDL and LDL particles. The average number of moles in lipoprotein particles was estimated by normalizing the lipid concentration in the lipoprotein fractions to the number of lipoprotein particles.

Lipid molecules in blood originate from blood cells or plasma. PCs and SMs are distributed outside the membrane in erythrocyte lipid bilayers. In contrast, PEs and PSs are distributed within the membrane^30^. To transport oxygen, erythrocytes circulate in the blood approximately 200–300 thousand times over a lifetime of 120 days. Since they come into contact with different cells in the course of their circulation, their properties reflect the status *in vivo*. Polycythemia is caused by an increase in red blood cells, which leads to thrombus and atherosclerosis^31^. In the present study, some lipid molecules in erythrocytes in the severe group were significantly changed compared with those in the mild group (Supplementary Fig. 4). In particular, PC and PE changed significantly at 8 and 16 months, respectively. These results indicate that membrane lipids are affected by the progression of coronary atherosclerosis in WHHLMI rabbits.

Lipoproteins also reflect atherosclerotic disease status. After VLDLs are formed from lipid molecules and apolipoproteins in hepatocytes, lipoproteins transport lipids from the liver to peripheral tissues. In lipoprotein metabolism, TGs and phospholipids in VLDLs are hydroxylated by lipoprotein- and hepatic lipases^32^, and the VLDLs are then converted to LDLs. In Watanabe heritable hyperlipidemic (WHHL) rabbits, LDL receptors are genetically deficient. Delayed clearance of VLDL and its metabolites increases LDL production and LDL accumulates in the blood^33^. LDL modification is known to trigger the development of atherosclerosis. For example, oxidized LDL and its constituent lipid molecules contribute to the formation of atherosclerotic lesions^34,35^. In the present study, the concentrations of most lipid subclasses correlated with the number of VLDL particles at all ages, whereas this trend was reduced for the LDL fractions (Fig. 3c, d). This association was observed regardless of the severity of the coronary atherosclerosis. A possible reason for this is that the lipid components of lipoproteins are modified after LDL particles are formed from VLDL particles by hepatic and lipoprotein lipases, regardless of the severity of coronary atherosclerosis. In addition, it is possible that the reduction in the ratio of lipid molecules by hepatic and/or lipoprotein lipases varies from individual to individual.

Although the apolipoproteins that regulate lipoprotein lipase and LDL receptor ligands did not change (Fig. 2c), the lipid composition of VLDL and LDL particles differed between the severe and mild groups. Among all lipid molecules, Cer molecules in serum and lipoprotein particles show a unique behavior. The number of Cer molecules that significantly changed in LDL particles ranked first at 8 months (Fig. 5b), and the ratio of Cer molecules that changed to those that were detected was highest among all lipid subclasses. Cer, which functions in intracellular signaling, is composed of a sphingosine backbone and FAs. In hepatic cells, Cers are not only metabolized to complex sphingolipids (e.g., SMs, gangliosides, and glucosylceramides) but are also incorporated into VLDLs^36^. To date, several functions of Cers in the pathogenesis of atherosclerosis have been reported. They induce apoptosis and suppress proliferation of cells involved in vascular endothelial cell damage^37,38^. They are also involved in atherogenesis through LDL aggregation in the arterial wall^39^. The increase in Cer molecules in VLDL and LDL particles requires further investigation of liver metabolism and coronary atherosclerosis phenotype at eight months. Except for Cer molecules, other lipid molecules, such as CE and PI molecules, were significantly decreased in LDL particles (Fig. 5a, b). PI promotes reverse cholesterol transport by increasing cholesterol efflux from peripheral tissues to HDL and enhancing its transport to the liver for excretion in bile and feces^40–42^. Recently, oxidized PI molecules derived from PI molecules have been identified in human oxidized LDL and atherosclerotic plaques^43^. These relationships will be elucidated in future studies using WHHLMI rabbits.

Limitations of the present study include the small number of WHHLMI rabbits. The current sample set did not show a significant sex difference in the PCA (Fig. 3a). These findings highlight universal alterations that occur independent of sex differences. Nevertheless, the study of sex-specific variations remains crucial for understanding the underlying biological mechanisms. We focused on the time-dependent profiles of apolipoproteins and lipid molecules that correlate with the severity of coronary atherosclerosis at 20 months. In future studies, it will be important to evaluate the correlation between lipid molecules and phenotypes at eight months. In addition, lipid metabolism in organs that interact with erythrocytes and/or lipoproteins should be characterized to understand the cause of blood alterations.

In conclusion, we characterized the lipids involved in the progression of coronary atherosclerosis in WHHLMI rabbits. In particular, Cer molecules in VLDL and LDL particles were associated with the severity of coronary atherosclerosis at an early stage, although apolipoproteins and common risk factors for abnormal lipid metabolism did not change. These findings may contribute to the elucidation of pathological mechanisms and the development of novel therapeutic drugs, as well as to further exploration of the components that are directly responsible for pathogenesis.

## Methods

### Chemicals and reagents

JIS special grade sodium chloride and sodium bromide, sodium hydroxide solution (1 mol L^−1^) and ethylenediaminetetraacetic acid disodium salt dihydrate were purchased from Wako Pure Chemical Industries, Ltd. (Osaka, Japan). Ammonium acetate was purchased from Sigma-Aldrich (St. Louis, MO). MS grade methanol was purchased from Kanto Chemical Co., Ltd. (Tokyo, Japan), HPLC-grade chloroform was purchased from Kishida Chemical (Osaka, Japan), and HPLC-grade distilled water was purchased from Wako Pure Chemical Industries. All synthetic lipid standards were purchased from Avanti Polar Lipids, Inc. (Alabaster, AL, USA). Carbon dioxide (99.5% grade, Yoshida Sanso Co., Ltd. (Fukuoka, Japan) was used as the mobile phase for SFC. Stable isotope-labeled peptides (^13^C- and ^15^N-labeled arginine and lysine: IEIPLPFGGK [^13^C_6_, ^15^N_2_] for apoB-100, VQESLSSYWDSAK [^13^C_6_, ^15^N_2_] for apoC-II, and AGQPWELALGR [^13^C_6_, ^15^N_4_] for apoE) were acquired from Eurofins Genomics (Tokyo, Japan). Stable isotope-labeled peptides (^13^C- and ^15^N-labeled arginine and lysine; GWVDAGISSLK [^13^C_6_, ^15^N_2_] for apoC-III) were acquired from Genenet Co. Ltd. (Fukuoka, Japan).

### Animals

All animal procedures were approved by the Kobe University Animal Care and Use Committee and were conducted in accordance with the Regulations for Animal Experimentation of Kobe University, the Act on Welfare and Management of Animals (Law No. 105, 1973, revised in 2006), Standards Relating to the Care and Management of Laboratory Animals and Relief of Pain (Notification No. 88, 2006), and Fundamental Guidelines for the Proper Conduct of Animal Experiments and Related Activities in Academic Research Institutions under the Jurisdiction of the Ministry of Education, Culture, Sports, Science and Technology (Notice No.71, 2006). Ten WHHLMI rabbits (males, *n* = 4) were housed individually in metal cages and fed standard rabbit chow (120 g day^−1^; LRC4, Oriental Yeast Co., Ltd, Tokyo, Japan) and water *ad libitum*. Room was kept with constant temperature (22 ± 2 °C) and lighting cycle (12 h light/dark). Ten WHHLMI rabbits were sacrificed at 20 months of age after blood sampling.

### Pathological analysis of WHHLMI rabbits

The degree of lesions in ten WHHLMI rabbits, including cross-sectional narrowing (CSN) and aortic surface lesion area, was previously evaluated^26^. Based on these results, the rabbits were divided into two groups (severe and mild)^26^. The severity of the coronary lesions was assessed using the coronary severity score, calculated as follows: WHHLMI rabbits were classified based on this score, with the severe group defined as > 2.0 and the mild group as < 0.5.

[Coronary severity score] = [average CSN] × (1 + [frequency of sections with 75%−89% CSN] × 10 + [frequency of sections with >90% CSN] × 20)

### Lipoprotein fractionation

Blood was collected from the marginal ear vein of WHHLMI rabbits after fasting for > 12 h. Blood samples were collected at 8, 12, 16, and 20 months to monitor lipid alterations in erythrocytes and lipoproteins related to the severity of coronary atherosclerosis. Plasma lipoproteins were fractionated by ultracentrifugation using a stepwise method (VLDL, d < 1.006 g mL^−1^ and LDL, 1.019 g mL^−1^ < d < 1.063 g mL^−1^), according to a previously described method^44,45^. Prior to fractionation, a sodium chloride solution (d = 1.006 g mL^−1^) and a sodium chloride/bromide solution (d = 1.182 g mL^−1^) were prepared. Additionally, intermediate-density solutions (d = 1.019 g mL^−1^ and 1.063 g mL^−1^) were prepared by mixing these two solutions. For the isolation of VLDL (d < 1.006 g mL^−1^), 3.2 mL of the sodium chloride solution was slowly added to 1.5 mL of plasma in ultracentrifugation tubes using a Pasteur pipette. After ultracentrifugation at 375,000 g for 5 h at 4 °C, the upper and lower layers were carefully separated with a microslicer and collected in clean tubes (upper VLDL fraction, 1.5 mL; lower fraction, 3.2 mL). The fractions were then subjected to a second ultracentrifugation under the same conditions for further purification. As with the VLDL isolation, LDL was purified by repeating the fractionation steps: first, the density was changed to 1.019 g mL^−1^ to remove fractions with d < 1.019 g mL^−1^ (with LDL and HDL collected from the lower layer), and then adjusted to 1.063 g mL^−1^ to eliminate fractions with d > 1.063 g mL^−1^ (with LDL collected from the upper layer). To achieve the target density of 1.019 g mL^−1^ or 1.063 g mL^−1^, a denser solution (d = 1.182 g mL^−1^) was initially added, followed by the appropriate 1.019 g mL^−1^ or 1.063 g mL^−1^ solution to fill the ultracentrifuge tube to 4.7 mL.

### Biochemical analysis

Cholesterol and TG concentrations were measured using a Cholesterol E-Test Assay Kit and Triglyceride E-Test Assay Kit (Wako Pure Chemical Industries, Ltd.), respectively, following the manufacturer’s instructions. To determine the sample volume (containing 50 μg of proteins) for apolipoproteins extraction, protein levels were measured using a Pierce™ BCA Protein Assay Kit and Micro BCA™ Protein Assay Kit (Thermo Fisher Scientific Inc., Waltham, MA) following the manufacturer’s instructions.

### Lipids extraction

Lipoprotein fractions were separated for lipid and apolipoprotein analyses. Lipid molecules in erythrocytes and lipoprotein fractions were extracted using Bligh and Dyer’s method, with minor modifications as described in our previous study^28,46,47^. Before lipid extraction, a dodecanoyl- or heptadecanoyl-based synthetic internal standard mixture was added to the samples for lipid quantification. The concentrations of the internal standards spiked into the samples were as follows: 1.6 μM (SM 18:1;O2/17:0 and Cer 18:1;O2/17:0); 8 μM (LPC 17:0); 16 μM (MG 17:0, DG 12:0_12:0, and TG 17:0_17:0_17:0); 130 μM (PE 17:0_17:0); 160 μM (LPE 17:1, PC 17:0_17:0, CE 17:0, and FA 17:0); and 1,300 μM (PS 17:0_17:0). Samples were mixed with 30 μL of internal standard mixture and 1000 μL of methanol/chloroform/water (10:5:3, v/v/v) and extracted using a vortex mixer and ultrasonication for 1 min and 5 min, respectively. The solution was then centrifuged at 16,000 *g* for 5 min at 4 °C, and 700 μL of the supernatant was transferred to a clean tube. The supernatant was mixed with 235 μL of chloroform and 155 μL of water using a vortex mixer for 1 min. After centrifugation at 16,000 *g* for 5 min at 4 °C, the lower layer was collected.

### Apolipoproteins extraction

Apolipoproteins were extracted in four steps: denaturation (reduction and alkylation) of apolipoproteins, methanol and chloroform precipitation, trypsin digestion, and desalting. In short, 50 μg of protein samples was diluted to 220 μL with lysis buffer (100 mM tris-HCl, pH 8.8, 7 M urea, 2% sodium dodecyl sulfate). Samples were mixed with 1 μL of tris (2-carboxyethyl) phosphine hydrochloride solution (final concentration, 2.5 mM) and incubated for 30 min at 37 °C for reduction of apolipoprotein. In addition, they were mixed with 2.5 μL of iodoacetamide solution (final concentration, 12.5 mM) and reacted for 30 min at room temperature (approximately 25 °C) for alkylation of apolipoprotein. After the reduction and alkylation steps, 1200 μL of cold methanol/chloroform/water (600:150:450, v/v/v) was added to samples, and they were centrifuged at 2,000 *g* for 15 min at 4 °C using a swing rotor. Aliquots of the organic layer were removed, and 450 μL of chloroform was added to them. After they were vortexed at the maximum setting for 1 min, they were centrifuged at 2,000 *g* for 15 min at 4 °C using the swing rotor. The supernatant was removed completely and 100 μL of ammonium bicarbonate (100 mM) was added to them. For digestion of proteins, 1 μL of trypsin (500 ng μL^−1^) were added to samples and incubated for 3 h at 37 °C. After this trypsin digestion, 1 μL of trypsin (500 ng μL^−1^) were added to samples again and incubated for 12 h at 37 °C to prevent autodigestion. Styrenedivinylbenzene was used for desalting of samples and the peptide extract was diluted to a final volume of 10 μL with a mixed solvent (0.1% trifluoroacetic acid, 1% acetonitrile and 98.9% water) for the nano-LC/MS analyses.

### Targeted quantitative lipidomics using SFC/MS

Lipid analysis was performed using SFC/MS. SFC-MS was performed using an ACQUITY Ultra Performance Convergence Chromatography (UPC^2^) system (Waters, Milford, MA) with a Xevo TQ-S micro tandem mass spectrometer (Waters) controlled by MassLynx software version 4.1 (Waters). An HPLC 515 pump (Waters) was used as the make-up pump and controlled manually. This was introduced to enhance ionization efficiency under the initial gradient conditions. The SFC and MS analytical conditions were optimized as described in a previous study^28^. The SFC conditions were as follows: mobile phase, liquefied carbon dioxide (A) and methanol/water (95/5, v/v) with 0.1% (w/v) ammonium acetate (B); make-up solvent, methanol/water (95/5, v/v) with 0.1% (w/v) ammonium acetate; injection volume, 1 μL; flow rate, 1.0 mL min^−1^ (mobile phase) and 0.2 mL min^−1^ (make-up pump); modifier gradient; 1% (B) (1 min), 1–65% (B) (11 min), 65% (B) (6 min), 65–1% (B) (0.1 min), 1% (B) (1.9 min); temperature of column manager, 50 °C; temperature of sample manager, 4 °C; analytical time, 20 min; and column, Torus DEA (3.0 × 100 mm, particle size: sub-1.7 μm, Waters). The MS analysis conditions were as follows: capillary voltage, 3.0 kV; desolvation temperature, 500 °C; cone gas flow rate, 50 L h^−1^; and desolvation gas flow rate, 1,000 L h^−1^. The MRM parameters were as follows: limit on number of MRM transitions, 150; dwell time, 1 ms; MS inter-scan and inter-channel delay, 2 ms; and polarity switch inter-scan, 15 ms.

### Targeted quantitative proteomics using nano-LC/MS

Apolipoproteins were quantified using nano-LC/MS. Nano-LC-MS was performed using an Ultimate 3000 system (Thermo Fisher Scientific, Waltham, MA, USA) with an LCMS-8060 tandem mass spectrometer (Shimadzu) and controlled using Labsolution software version 5.6 (Shimadzu). Nano-LC and MS analytical conditions were optimized based on our previous study^14^. The nano-LC conditions were as follows: mobile phase, water with 0.1% formic acid (A) and acetonitrile with 0.1% formic acid (B); injection volume, 1 μL; flow rate, 250 nL min^−1^; modifier gradient; 0% (B) (7 min), 0–65% (B) (38 min), 65–100% (B) (5 min), 100% (B) (15 min), 100–75% (B) (2 min), 75– 0% (B) (8 min), 0% (B) (15 min); temperature of column manager, room temperature (approximately 25 °C); temperature of sample manager, 4 °C; analytical time, 90 min; and column, L-column 2 (0.1 × 150 mm, particle size: 3 μm, CERI, Tokyo, Japan). The MS analysis conditions were as follows: electrospray voltage, 2.0 kV; desolvation line, 250 °C; heat block temperature, 400 °C; drying gas flow, 0 L h^−1^; and gas pressure of collision-induced dissociation, 0.27 MPa. Each peptide (IEIPLPFGGK for apoB-100, VQESLSSYWDSAK for apoC-II, GWVDAGISSLK for apoC-III, and AGQPWELALGR for apoE) was measured to quantify the apolipoproteins. The MRM transitions of each peptide are summarized in Supplementary Table 2.

### Data analysis

Serum lipidomics data were obtained from our previous study^26^. In the present study, we analyzed lipid molecules in erythrocytes and lipid molecules and apolipoproteins in lipoprotein fractions. The annotation and peak picking of the lipid molecules was carried out using the MassLynx software version 4.1. Lipidomic data were normalized to the number of erythrocytes and lipoprotein particles. In this study, the quantification type of the SFC/MS system corresponds to Level 2 of the Lipidomics Standards Initiative International Guideline, indicating that the lipid subclass of the analyte and internal standard are identical, with co-ionization of both analyte and internal standard^48,49^. Additionally, the nano-LC/MS system enables absolute quantification, as fully prepared ^13^C- and ^15^N-labeled internal standards were utilized in this analysis.

### Visualization and statistical analysis

The individual lipid profiles of the mild and severe groups were visualized using a volcano plot. Statistical significance was determined using either the Student’s or Welch’s *t*-test after *F*-test for variance. Correlations between apoB-100 and lipid subclasses, and between coronary severity scores and lipid molecules were evaluated using Pearson’s correlation coefficient. Statistical significance was set at ^*^*p* < 0.05 and ^**^*p* < 0.01.

PCA was performed using the prcomp function in the R programming environment on autoscaled data (mean-centered and variance-scaled). The explained variance was calculated as the ratio of the squared singular values, and the results were visualized using ggplot2. PCA score plots were generated for the first two components, with data points color-coded by age (4, 8, 12, 16, and 20 months) and shaped by sex (male and female). The heatmap was generated using the pheatmap, tidyverse, and openxlsx packages. Data matrices were prepared by excluding metadata columns (age and lipoprotein) and then annotated with metadata for visualization. It was generated without clustering for rows or columns using a custom divergent color scale ranging from blue to red, with white as the midpoint. Lipoprotein types were color-coded (VLDL: cyan; LDL: yellow-green). Network analysis of lipid molecules was performed using the R packages igraph, tidygraph, and ggraph. First, Pearson correlation coefficients were calculated between each lipid molecule and the response variable (coronary severity score), and lipids with |R| > 0.6 were selected. Next, pairwise correlation coefficients were calculated between the selected lipids, and lipid pairs with |R| > 0.6 were connected as edges to construct the network. Nodes represented individual lipid molecules, with node size corresponding to the average lipid abundance and node color indicating lipid subclasses. Edge thickness reflects the correlation strength (|R|) between lipid pairs. The color of the edges was assigned based on the direction of the correlation, with positive correlations in gray and negative correlations in beige. The Kamada-Kawai layout was used to highlight the relationships between lipid molecules.

## Data availability

The numerical source data behind the graphs can be found in the Supplementary Datafile.

## Acknowledgments

This study was partly supported by the AMED-BINDS (JP22ama121055 for T.B.) from the Japan Agency for Medical Research and Development (AMED); a grant from the JST-CREST Program (JPMJCR22N5 for Y.I.) of the Japan Science and Technology Agency (JST); JST A-STEP (JPMJTR204J for T.B.); JST-NBDC (JPMJND2305 for Y.I.); JST Moonshot (JPMJMS2011-62 for Y.I.); and KAKENHI (JP22H05185 for Y.I.; JP21K14472 for K.N.; JP23K13611 for K.H.) from the Japan Society for the Promotion of Science (JSPS). This work was also supported in part by the MEXT Cooperative Research Project Program, Medical Research Center Initiative for High Depth Omics, and CURE:JPMXP1323015486 for Medical Institute of Bioregulation, Kyushu University.

## Author contributions

H.T., Y.I., T.K., M.M., M.S., and T.B. designed the study, including the sampling points for WHHLMI rabbits and analytical strategies for lipids and apolipoproteins. Y.K. performed time-course sampling of WHHLMI rabbits. H.T. and Y.K. fractionated lipoproteins and erythrocytes. H.T. extracted and analyzed the lipid molecules. H.T., K.N., and K.H. extracted and analyzed the apolipoproteins. H.T. and Y.I. wrote the manuscript. The other authors revised and approved the manuscript.

