## Supporting Information for "Characterizing lipoprotein profiles in coronary atherosclerosis development through quantitative lipidomics and proteomics approaches"

#### Contents

Supplementary Figures 1–4

Supplementary Tables 1–8

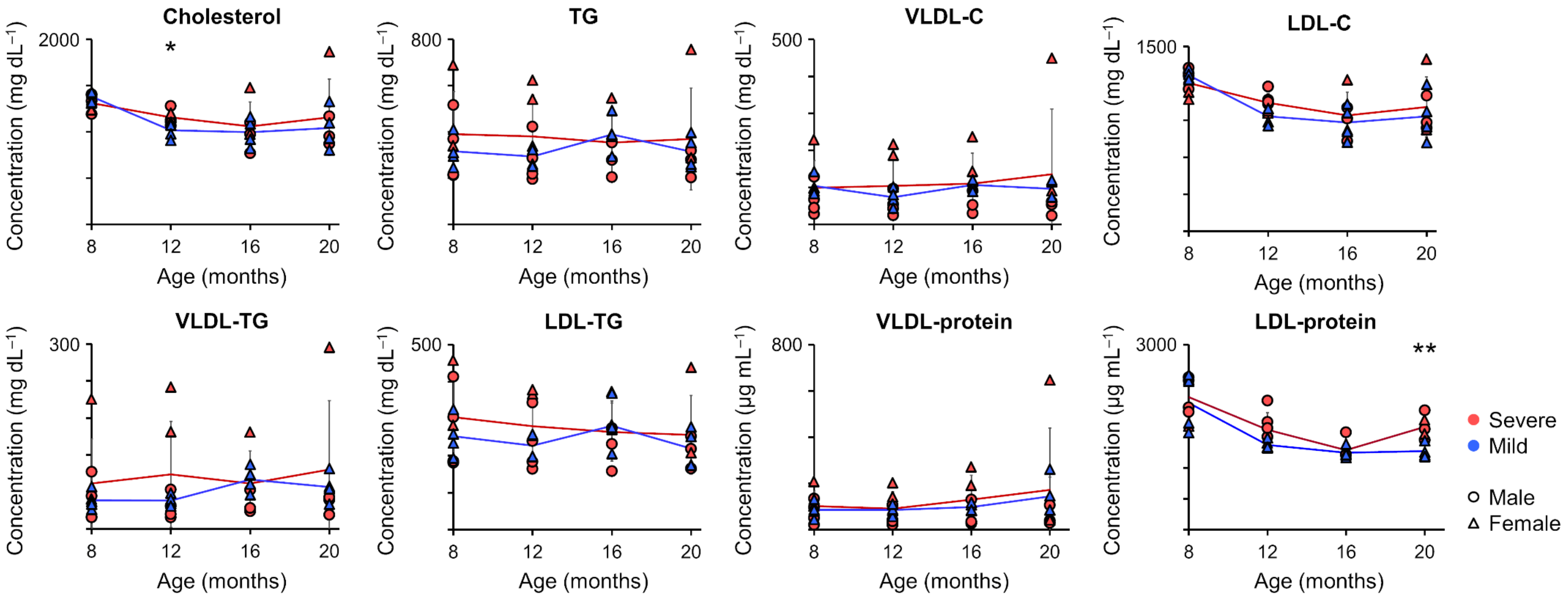


**Supplementary Fig. 1 Time-course alterations in total cholesterol, TG, and proteins.** Red and blue indicate severe and mild groups, respectively, while circles and triangles represent males and females, respectively. Error bars represent standard deviations of biological replicates. Statistical significance was determined using Student's or Welch's *t*-test followed by *F*-test for variance ($^{*}p < 0.05$ and $^{**}p < 0.01$).

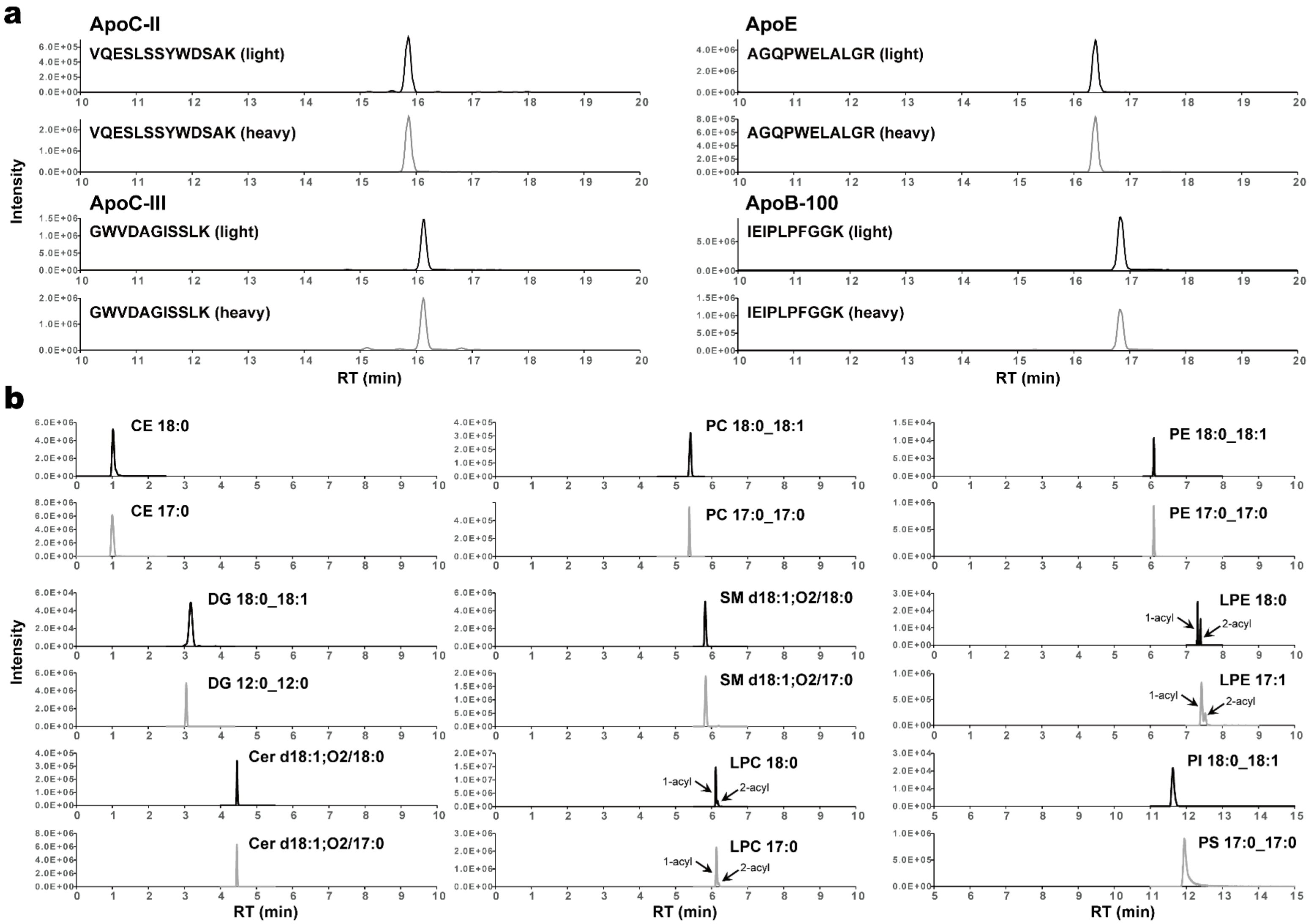


**Supplementary Fig. 2 Multi-quantitative omics of plasma lipoprotein fractions.** Chromatograms of apolipoproteins (**a**) and representative lipid molecules (**b**) in the LDL fraction are shown.

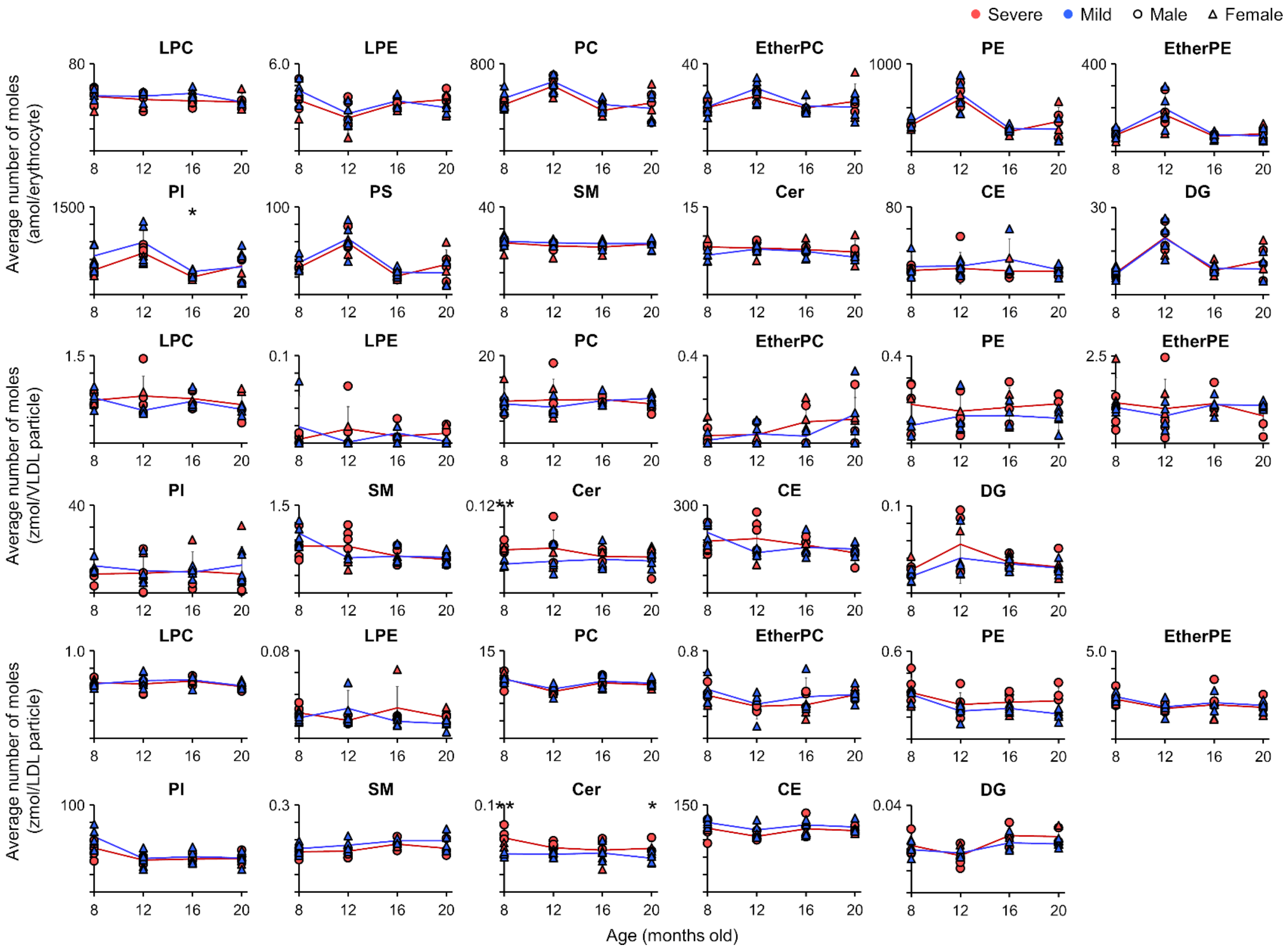


**Supplementary Fig. 3 Time-course alterations of lipid subclasses in erythrocytes and lipoprotein particles.** Red and blue indicate severe and mild groups, respectively, while circles and triangles represent males and females, respectively. Error bars represent standard deviations of biological replicates. Statistical significance was determined using Student's or Welch's *t*-test following an *F*-test for variance ($^{*}p < 0.05$ and $^{**}p < 0.01$).

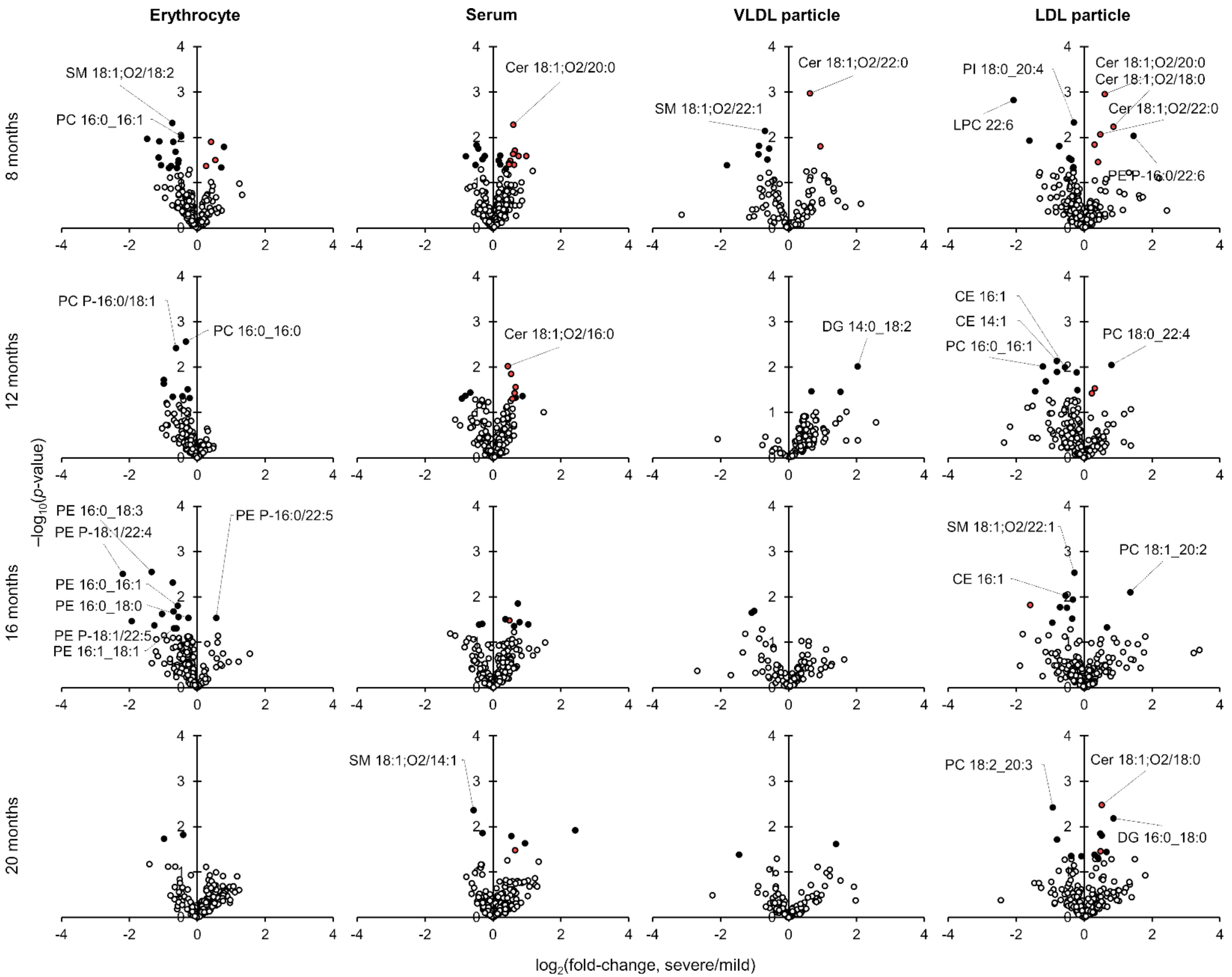


**Supplementary Fig. 4 Lipid profiling of erythrocytes and lipoprotein particles.** Characteristics of lipid molecules at each age are shown using volcano plots. Black circles represent all lipids that showed significant changes ($p < 0.05$), while red circles indicate significantly altered Cer molecules.

### Supplementary Tables

#### Supplementary Table 1 Sample information for WHHLMI rabbits.

| Group | Sample No. | Rabbit ID | Sex | Age of death (months) | Date of death | Examined sections | Sections with lesions | Sections with >75% CSN | Sections with >90% CSN | Average CSN across all sections (%) | Coronary severity score | Aortic lesion (%) | Comment |
|---|---|---|---|---|---|---|---|---|---|---|---|---|---|
| Severe | C1 | 26-135-7f | female | 20 | 2015/2/17 | 21 | 20 | 12 | 3 | 66.1 ± 29.6 | 5.38 | 95.33 | |
| | C2 | 26-135-11f | female | 20 | 2015/2/17 | 22 | 19 | 6 | 4 | 42.4 ± 35.1 | 2.35 | 92.59 | |
| | C5 | 26-135-3m | male | 20 | 2015/2/17 | 18 | 18 | 3 | 2 | 55.2 ± 20.6 | 2.08 | 96.96 | |
| | C6 | 26-135-17m | male | 20 | 2015/2/17 | 22 | 21 | 13 | 8 | 71.0 ± 31.1 | 7.48 | 84.07 | |
| | D5 | 25-136-17m | male | 20 | 2015/3/24 | 22 | 10 | 10 | 7 | 52.2 ± 41.1 | 4.55 | 94.60 | |
| | D6 | 26-136-19m | male | 15 | 2014/9/25 | 15 | 15 | 14 | 13 | 94.2 ± 7.2 | 17.89 | 98.28 | |
| Mild | C3 | 24-135-18f | female | 20 | 2015/2/17 | 19 | 3 | 0 | 0 | 4.2 ± 11.8 | 0.04 | 94.35 | |
| | C4 | 24-135-19f | female | 20 | 2015/2/17 | 23 | 16 | 0 | 0 | 32.2 ± 29.4 | 0.32 | 94.00 | |
| | D1 | 26-136-2f | female | 20 | 2015/3/24 | 21 | 1 | 0 | 0 | 0.0 ± 0.2 | 0.00 | 94.88 | |
| | D4 | 25-136-14f | female | 20 | 2015/3/24 | 19 | 17 | 0 | 0 | 44.7 ± 23.6 | 0.45 | 95.22 | |
| | D2 | 26-136-11m | male | 16 | 2014/11/6 | | | | | | | | Exclude further analysis due to renal tumor |
| | D3 | 26-137-12f | female | 18 | 2015/1/22 | | | | | | | | Exclude further analysis due to lymphoma |

### Supplementary Table 2 Biochemical assay for WHHLMI rabbits.

| | 8 months | | | | 12 months | | | | 16 months | | | | 20 months | | | |
|---|---|---|---|---|---|---|---|---|---|---|---|---|---|---|---|---|
| | Severe | | Mild | | Severe | | Mild | | Severe | | Mild | | Severe | | Mild | |
| | Mean | SD | Mean | SD | Mean | SD | Mean | SD | Mean | SD | Mean | SD | Mean | SD | Mean | SD |
| Body weight (kg) | 2.97 | 0.14 | 3.12 | 0.11 | 3.15 | 0.28 | 3.44 | 0.07 | 3.37 | 0.28 | 3.65 | 0.08 | 3.26 | 0.31 | 3.58 | 0.12 |
| Cholesterol (mg $dL^{-1}$) | 1312.60 | 85.14 | 1380.28 | 57.27 | 1157.14 | 75.45 | 1017.66 | 90.25 | 1059.28 | 262.06 | 998.16 | 155.40 | 1157.21 | 412.27 | 1041.99 | 225.44 |
| TG (mg $dL^{-1}$) | 390.23 | 184.68 | 315.76 | 69.42 | 381.18 | 176.34 | 293.31 | 43.67 | 354.55 | 128.82 | 388.76 | 81.70 | 368.98 | 220.52 | 315.03 | 71.96 |
| VLDL-C (mg $dL^{-1}$) | 99.15 | 72.99 | 103.96 | 26.60 | 103.69 | 79.98 | 73.20 | 24.11 | 110.34 | 82.32 | 106.55 | 14.33 | 135.42 | 177.02 | 95.77 | 25.45 |
| VLDL-TG (mg $dL^{-1}$) | 73.96 | 72.12 | 46.39 | 15.76 | 88.48 | 85.99 | 46.20 | 9.06 | 74.13 | 52.11 | 80.44 | 21.19 | 96.45 | 111.76 | 68.07 | 23.56 |
| LDL-C (mg $dL^{-1}$) | 1203.21 | 100.19 | 1265.94 | 41.26 | 1042.45 | 77.32 | 933.08 | 72.63 | 939.20 | 190.98 | 881.20 | 140.52 | 1007.82 | 243.94 | 932.82 | 201.60 |
| LDL-TG (mg $dL^{-1}$) | 304.03 | 114.32 | 252.95 | 55.25 | 279.51 | 95.42 | 227.12 | 34.18 | 264.03 | 78.46 | 280.61 | 67.38 | 256.63 | 106.18 | 219.94 | 53.33 |
| VLDL-protein ($\mu$g $mL^{-1}$) | 102.10 | 65.64 | 86.07 | 35.46 | 91.75 | 71.66 | 85.03 | 21.99 | 130.93 | 104.03 | 98.07 | 17.20 | 172.78 | 266.75 | 144.61 | 83.27 |
| LDL-protein ($\mu$g $mL^{-1}$) | 2149.67 | 336.71 | 2056.75 | 474.22 | 1622.00 | 278.33 | 1375.50 | 75.25 | 1292.60 | 159.79 | 1249.00 | 97.57 | 1670.20 | 190.55 | 1275.75 | 115.52 |

**Supplementary Table 3 MRM transitions used for apolipoprotein quantification.**

| Apolipoproteins | Peptide sequences | Label | Collision energy (eV) | Q1 (*m/z*) | Q3 (*m/z*) | | | |
|---|---|---|---|---|---|---|---|---|
| | | | | | y8 | y7 | y6 | y5 |
| ApoB-100 | IEIPLPFGGK | Light | −20.9 | 535.8 | 828.5 | 715.4 | 618.4 | 505.3 |
| | | Heavy | −20.9 | 539.8 | 836.5 | 723.4 | 626.4 | 513.3 |
| ApoC-II | VQESLSSYWDSAK | Light | −29.5 | 750.4 | 943.4 | 856.4 | 769.4 | |
| | | Heavy | −29.5 | 754.4 | 951.4 | 864.4 | 777.4 | |
| ApoC-III | GWVDAGISSLK | Light | −22.2 | 566.8 | 790.5 | 675.4 | 604.4 | |
| | | Heavy | −22.2 | 570.8 | 798.5 | 683.4 | 612.4 | |
| ApoE | AGQPWELALGR | Light | −23.5 | 599.3 | 941.5 | 844.5 | 658.4 | |
| | | Heavy | −23.5 | 604.3 | 951.5 | 854.5 | 668.4 | |

Internal standard peptides (heavy) were labeled with [$^{13}C_6$, $^{15}N_2$]-lysine (Lys, K) or [$^{13}C_6$, $^{15}N_4$]-arginine (Arg, R).

**Supplementary Table 4 Quantitative proteomics for lipoprotein fractions.**

| | Quantification (pmol mL$^{-1}$) | | | | | | | | | | | | | | | |
|---|---|---|---|---|---|---|---|---|---|---|---|---|---|---|---|---|
| | 8 months | | | | 12 months | | | | 16 months | | | | 20 months | | | |
| | Severe | | Mild | | Severe | | Mild | | Severe | | Mild | | Severe | | Mild | |
| | Mean | SD | Mean | SD | Mean | SD | Mean | SD | Mean | SD | Mean | SD | Mean | SD | Mean | SD |
| VLDL apoB-100 | 54 | 28 | 51 | 19 | 54 | 44 | 58 | 10 | 79 | 52 | 65 | 17 | 74 | 85 | 87 | 43 |
| VLDL apoC-II | 18 | 13 | 8 | 5 | 23 | 20 | 11 | 8 | 16 | 9 | 18 | 8 | 27 | 25 | 37 | 20 |
| VLDL apoC-III | 122 | 91 | 50 | 28 | 120 | 118 | 78 | 52 | 126 | 77 | 124 | 38 | 265 | 343 | 242 | 60 |
| VLDL apoE | 219 | 107 | 246 | 88 | 148 | 63 | 172 | 50 | 214 | 119 | 189 | 41 | 222 | 244 | 201 | 48 |
| LDL apoB-100 | 1187 | 137 | 1055 | 153 | 1059 | 167 | 932 | 120 | 843 | 138 | 787 | 86 | 1055 | 169 | 822 | 86 |
| LDL apoC-II | 874 | 333 | 802 | 281 | 782 | 187 | 704 | 107 | 701 | 147 | 549 | 51 | 959 | 184 | 718 | 125 |
| LDL apoC-III | 21330 | 2310 | 15871 | 5196 | 18413 | 3773 | 21149 | 2239 | 17938 | 6526 | 18403 | 3473 | 22245 | 6059 | 18568 | 2829 |
| LDL apoE | 2153 | 523 | 1635 | 376 | 1534 | 628 | 1551 | 393 | 1164 | 96 | 1011 | 128 | 2113 | 689 | 1398 | 367 |

Proteomics data were normalized using the ion abundance of internal standards added after apolipoprotein extraction.

### Supplementary Table 5 Quantitative lipidomics for serum.

| Lipids | Quantification (nmol mL$^{-1}$) | | | | | | | | | | | | | | | |
|---|---|---|---|---|---|---|---|---|---|---|---|---|---|---|---|---|
| | 8 months | | | | 12 months | | | | 16 months | | | | 20 months | | | |
| | Severe | | Mild | | Severe | | Mild | | Severe | | Mild | | Severe | | Mild | |
| | Mean | SD | Mean | SD | Mean | SD | Mean | SD | Mean | SD | Mean | SD | Mean | SD | Mean | SD |
| FA 14:0 | 12.6 | 2.2 | 13.9 | 2.6 | 12.8 | 2.0 | 16.0 | 2.6 | 15.6 | 2.9 | 20.8 | 3.3 | 15.1 | 4.3 | 16.7 | 2.7 |
| FA 16:0 | 192.2 | 29.5 | 199.3 | 36.4 | 210.5 | 35.9 | 216.1 | 43.7 | 249.4 | 45.5 | 291.2 | 81.9 | 244.4 | 74.1 | 215.0 | 17.1 |
| FA 16:1 | 27.3 | 19.7 | 37.3 | 6.7 | 31.5 | 12.9 | 55.5 | 18.9 | 44.0 | 20.6 | 105.5 | 59.2 | 47.8 | 27.9 | 64.4 | 13.8 |
| FA 18:0 | 176.8 | 19.4 | 182.2 | 41.3 | 191.1 | 24.8 | 179.5 | 15.3 | 207.4 | 26.5 | 198.6 | 48.2 | 201.6 | 50.1 | 177.0 | 17.2 |
| FA 18:1 | 157.7 | 72.7 | 173.2 | 31.9 | 193.3 | 63.0 | 223.7 | 68.2 | 282.7 | 95.0 | 372.5 | 151.2 | 282.8 | 139.6 | 274.6 | 48.9 |
| FA 18:2 | 130.4 | 62.1 | 130.5 | 27.5 | 155.2 | 60.8 | 178.5 | 45.5 | 220.8 | 66.1 | 288.8 | 118.3 | 220.2 | 103.8 | 216.8 | 39.5 |
| FA 18:3 | 32.5 | 24.1 | 36.5 | 8.2 | 31.8 | 17.8 | 50.0 | 15.0 | 42.7 | 17.1 | 76.3 | 39.7 | 37.7 | 20.4 | 44.8 | 10.7 |
| FA 20:4 | 3.2 | 0.5 | 3.7 | 0.8 | 4.3 | 1.6 | 4.8 | 1.8 | 4.6 | 1.5 | 6.6 | 2.1 | 5.0 | 3.0 | 5.0 | 0.8 |
| FA total | 732.7 | 210.3 | 776.8 | 135.8 | 830.5 | 205.3 | 924.1 | 199.1 | 1067.1 | 255.4 | 1360.2 | 502.0 | 1054.5 | 416.4 | 1014.2 | 121.2 |
| LPC 14:0 | 0.6 | 0.2 | 0.5 | 0.2 | 0.5 | 0.1 | 0.4 | 0.2 | 0.4 | 0.1 | 0.4 | 0.2 | 0.5 | 0.1 | 0.5 | 0.1 |
| LPC 16:0 | 268.4 | 23.9 | 250.0 | 22.0 | 243.4 | 32.3 | 226.4 | 14.1 | 239.0 | 45.1 | 203.2 | 32.3 | 242.1 | 34.5 | 225.1 | 30.9 |
| LPC 16:1 | 4.5 | 0.9 | 5.2 | 0.7 | 4.3 | 1.4 | 4.9 | 0.6 | 4.5 | 1.2 | 5.1 | 0.6 | 4.2 | 1.0 | 5.4 | 0.6 |
| LPC 18:0 | 379.1 | 39.5 | 326.7 | 12.1 | 345.6 | 33.2 | 334.4 | 25.6 | 340.2 | 51.8 | 307.6 | 22.4 | 313.6 | 35.5 | 310.9 | 17.6 |
| LPC 18:1 | 116.3 | 20.8 | 110.4 | 14.0 | 105.5 | 26.6 | 103.7 | 11.9 | 110.4 | 29.2 | 90.0 | 16.9 | 98.3 | 10.2 | 95.3 | 18.2 |
| LPC 18:2 | 251.0 | 23.0 | 223.6 | 5.5 | 230.2 | 33.6 | 226.5 | 19.0 | 234.4 | 29.1 | 207.1 | 20.4 | 210.4 | 32.8 | 210.6 | 23.4 |
| LPC 18:3 | 6.1 | 1.4 | 6.1 | 0.8 | 5.0 | 1.4 | 6.0 | 0.4 | 4.8 | 1.3 | 4.3 | 0.7 | 4.6 | 0.6 | 4.4 | 0.9 |
| LPC 20:1 | 2.4 | 0.6 | 2.3 | 0.8 | 2.0 | 0.6 | 2.2 | 0.4 | 2.3 | 0.6 | 1.6 | 0.4 | 1.9 | 0.3 | 1.8 | 0.2 |
| LPC 20:2 | 4.3 | 0.6 | 4.2 | 1.4 | 3.7 | 1.2 | 3.6 | 0.9 | 4.3 | 1.0 | 3.1 | 1.1 | 3.2 | 0.2 | 3.3 | 0.6 |
| LPC 20:3 | 5.5 | 0.7 | 5.5 | 1.1 | 4.8 | 1.3 | 5.0 | 0.8 | 5.2 | 1.1 | 4.5 | 1.1 | 4.5 | 0.7 | 4.8 | 1.6 |
| LPC 20:4 | 12.9 | 1.7 | 12.4 | 0.2 | 11.7 | 1.4 | 13.0 | 1.9 | 12.6 | 2.2 | 12.9 | 0.8 | 12.7 | 3.1 | 14.6 | 3.6 |

| | | | | | | | | | | | | | | | | |
|---|---|---|---|---|---|---|---|---|---|---|---|---|---|---|---|---|
| LPC 20:5 | 0.3 | 0.2 | 0.4 | 0.0 | 0.3 | 0.1 | 0.2 | 0.1 | 0.3 | 0.1 | 0.2 | 0.1 | 0.2 | 0.1 | 0.2 | 0.1 |
| LPC 22:0 | 0.9 | 0.1 | 0.8 | 0.2 | 0.8 | 0.2 | 0.8 | 0.1 | 0.7 | 0.2 | 0.5 | 0.3 | 0.7 | 0.3 | 0.7 | 0.3 |
| LPC 22:4 | 1.4 | 0.3 | 1.3 | 0.2 | 1.3 | 0.3 | 1.2 | 0.3 | 1.3 | 0.4 | 1.3 | 0.3 | 1.5 | 0.3 | 1.6 | 0.3 |
| LPC 22:5 | 1.7 | 0.4 | 1.7 | 0.4 | 1.7 | 0.3 | 1.6 | 0.2 | 1.5 | 0.3 | 1.5 | 0.4 | 1.6 | 0.8 | 2.4 | 0.9 |
| LPC 22:6 | 0.7 | 0.1 | 0.5 | 0.2 | 0.6 | 0.2 | 0.6 | 0.0 | 0.6 | 0.2 | 0.6 | 0.1 | 0.7 | 0.2 | 0.8 | 0.3 |
| LPC total | 1056.1 | 89.0 | 951.8 | 37.7 | 961.3 | 125.5 | 930.5 | 63.6 | 962.4 | 148.3 | 844.0 | 85.9 | 900.7 | 39.6 | 882.3 | 77.2 |
| LPE 16:0 | 3.1 | 1.0 | 2.3 | 0.7 | 2.4 | 0.6 | 1.8 | 0.6 | 2.7 | 1.2 | 1.8 | 0.3 | 2.7 | 0.8 | 1.4 | 0.5 |
| LPE 18:0 | 13.4 | 3.0 | 10.2 | 2.1 | 10.3 | 2.3 | 9.8 | 2.1 | 10.8 | 3.6 | 8.1 | 0.5 | 11.3 | 5.5 | 6.8 | 1.2 |
| LPE 18:1 | 4.1 | 1.0 | 4.0 | 0.7 | 3.2 | 1.5 | 3.0 | 1.0 | 3.8 | 2.1 | 2.7 | 1.3 | 4.1 | 1.4 | 3.3 | 0.8 |
| LPE 18:2 | 12.8 | 3.3 | 11.7 | 1.9 | 12.1 | 3.8 | 9.8 | 1.2 | 12.3 | 2.8 | 7.8 | 1.0 | 12.4 | 2.7 | 10.5 | 2.4 |
| LPE total | 33.3 | 7.3 | 28.2 | 4.6 | 27.9 | 7.9 | 24.3 | 3.8 | 29.5 | 8.1 | 20.4 | 2.4 | 30.5 | 9.6 | 21.9 | 3.8 |
| PC 16:0_16:0 | 92.9 | 13.1 | 107.0 | 9.0 | 81.5 | 15.4 | 82.6 | 7.3 | 85.3 | 31.6 | 61.5 | 11.7 | 91.1 | 42.5 | 78.3 | 14.2 |
| PC 16:0_16:1 | 11.7 | 6.6 | 13.0 | 2.3 | 9.4 | 4.0 | 11.4 | 2.5 | 12.6 | 7.3 | 10.9 | 4.2 | 12.6 | 9.2 | 12.2 | 2.6 |
| PC 16:0_18:0 | 163.8 | 29.5 | 162.2 | 11.2 | 144.6 | 23.4 | 137.8 | 22.4 | 137.7 | 43.7 | 112.0 | 21.6 | 140.9 | 44.1 | 117.6 | 33.3 |
| PC 16:0_18:1 | 900.1 | 247.6 | 882.6 | 126.3 | 765.7 | 195.0 | 712.1 | 106.1 | 829.8 | 329.9 | 588.6 | 179.5 | 831.9 | 310.4 | 674.5 | 265.8 |
| PC 16:0_18:2 | 2202.7 | 363.1 | 2173.4 | 162.6 | 1932.7 | 279.8 | 1757.1 | 104.1 | 1901.4 | 457.7 | 1570.4 | 169.1 | 1893.4 | 429.5 | 1683.8 | 231.0 |
| PC 16:0_18:3 | 79.6 | 20.0 | 93.3 | 4.7 | 68.4 | 17.4 | 64.9 | 6.0 | 54.7 | 24.9 | 45.1 | 10.0 | 63.1 | 23.9 | 52.7 | 15.0 |
| PC 16:0_20:0 | 9.5 | 1.9 | 10.1 | 4.9 | 7.5 | 2.9 | 8.5 | 2.3 | 6.8 | 2.2 | 6.1 | 4.2 | 7.4 | 1.9 | 5.6 | 1.3 |
| PC 16:0_20:2 | 25.4 | 6.9 | 19.6 | 3.2 | 18.8 | 7.0 | 15.6 | 6.3 | 22.9 | 9.0 | 13.9 | 6.8 | 21.9 | 14.3 | 16.8 | 11.6 |
| PC 16:0_20:3 | 55.5 | 10.9 | 55.9 | 4.3 | 42.1 | 9.3 | 48.2 | 9.8 | 52.3 | 28.8 | 44.0 | 16.3 | 57.9 | 40.4 | 48.8 | 30.8 |
| PC 16:0_20:4 | 110.5 | 25.0 | 122.0 | 9.9 | 101.7 | 23.0 | 102.1 | 14.6 | 105.5 | 42.6 | 103.1 | 15.7 | 128.2 | 87.1 | 116.0 | 38.3 |
| PC 16:0_22:4 | 11.0 | 3.2 | 12.1 | 3.2 | 10.5 | 5.1 | 11.7 | 1.9 | 12.0 | 5.8 | 10.7 | 4.4 | 14.0 | 13.0 | 15.7 | 7.8 |
| PC 16:0_22:5 | 18.1 | 4.3 | 14.9 | 4.0 | 13.7 | 4.1 | 12.6 | 5.1 | 13.3 | 5.6 | 13.1 | 2.1 | 27.7 | 29.3 | 20.0 | 13.5 |
| PC 16:1_18:1 | 15.5 | 10.5 | 15.8 | 1.6 | 11.2 | 5.5 | 11.8 | 1.9 | 12.2 | 9.5 | 14.3 | 5.0 | 14.0 | 8.3 | 12.5 | 3.2 |
| PC 16:1_18:2 | 32.4 | 8.6 | 30.7 | 6.1 | 26.9 | 8.1 | 30.7 | 6.7 | 29.8 | 14.1 | 26.4 | 4.0 | 32.9 | 13.9 | 36.7 | 4.6 |

| | | | | | | | | | | | | | | | | |
|---|---|---|---|---|---|---|---|---|---|---|---|---|---|---|---|---|
| PC 18:0_18:0 | 53.3 | 9.0 | 50.0 | 4.3 | 47.6 | 13.8 | 42.5 | 11.4 | 49.0 | 14.6 | 36.4 | 6.4 | 46.7 | 9.5 | 41.3 | 14.0 |
| PC 18:0_18:1 | 686.3 | 102.5 | 658.0 | 58.4 | 602.5 | 150.8 | 591.9 | 81.1 | 648.7 | 202.7 | 525.3 | 67.0 | 585.4 | 134.1 | 556.0 | 101.5 |
| PC 18:0_18:2 | 2300.7 | 242.2 | 2154.0 | 161.1 | 2096.5 | 288.5 | 1989.5 | 163.6 | 2148.9 | 278.6 | 1957.2 | 59.6 | 1968.5 | 291.5 | 1958.9 | 97.2 |
| PC 18:0_18:3 | 54.9 | 15.2 | 59.5 | 8.3 | 47.8 | 16.3 | 48.7 | 5.8 | 40.7 | 17.9 | 37.6 | 8.0 | 44.4 | 13.3 | 39.5 | 10.7 |
| PC 18:0_20:2 | 31.3 | 5.4 | 24.7 | 2.1 | 24.5 | 9.5 | 25.8 | 10.8 | 28.2 | 8.4 | 19.7 | 10.7 | 28.2 | 9.4 | 22.0 | 9.4 |
| PC 18:0_20:3 | 70.7 | 7.9 | 68.9 | 1.5 | 57.0 | 12.0 | 64.0 | 6.6 | 63.1 | 19.2 | 53.6 | 13.8 | 64.9 | 47.6 | 68.3 | 31.1 |
| PC 18:0_20:4 | 147.4 | 18.3 | 143.4 | 21.0 | 143.7 | 22.7 | 155.7 | 17.1 | 143.7 | 40.7 | 154.1 | 17.5 | 163.8 | 90.1 | 168.5 | 35.4 |
| PC 18:0_22:4 | 11.9 | 4.2 | 11.5 | 3.0 | 9.9 | 2.3 | 7.8 | 4.1 | 10.1 | 5.7 | 10.8 | 3.9 | 11.6 | 6.5 | 13.0 | 2.8 |
| PC 18:0_22:5 | 12.2 | 2.5 | 12.5 | 3.3 | 12.1 | 3.0 | 8.9 | 3.1 | 12.6 | 7.3 | 14.5 | 2.3 | 17.8 | 12.4 | 17.1 | 4.3 |
| PC 18:1_18:1 | 77.7 | 19.8 | 79.0 | 10.7 | 73.2 | 18.9 | 68.2 | 14.2 | 82.8 | 40.6 | 57.4 | 11.0 | 80.3 | 19.1 | 72.8 | 20.5 |
| PC 18:1_18:2 | 445.5 | 64.4 | 427.8 | 25.5 | 404.3 | 95.4 | 385.5 | 23.9 | 410.4 | 120.8 | 336.6 | 32.0 | 369.2 | 44.7 | 371.1 | 55.7 |
| PC 18:1_18:3 | 26.1 | 10.5 | 19.4 | 4.2 | 15.8 | 6.6 | 16.5 | 1.3 | 15.8 | 6.0 | 13.3 | 6.0 | 14.6 | 3.2 | 14.3 | 4.1 |
| PC 18:1_20:4 | 19.2 | 4.9 | 19.4 | 5.4 | 15.5 | 2.9 | 18.9 | 1.7 | 17.3 | 6.8 | 16.6 | 2.9 | 14.9 | 10.1 | 25.7 | 8.2 |
| PC 18:2_18:2 | 432.5 | 63.4 | 395.8 | 29.0 | 370.1 | 63.4 | 331.8 | 33.6 | 364.1 | 74.7 | 293.2 | 16.5 | 296.1 | 26.2 | 288.1 | 22.6 |
| PC 18:2_18:3 | 34.6 | 5.6 | 32.6 | 3.5 | 26.3 | 7.2 | 19.2 | 5.5 | 19.6 | 8.0 | 16.9 | 4.0 | 21.8 | 3.9 | 18.6 | 6.4 |
| PC 18:2_20:1 | 12.7 | 4.9 | 9.7 | 1.5 | 8.8 | 4.4 | 7.5 | 2.9 | 9.4 | 2.4 | 7.8 | 4.1 | 6.3 | 2.4 | 7.1 | 4.5 |
| PC 18:2_20:2 | 22.9 | 5.8 | 20.1 | 9.4 | 20.3 | 9.6 | 17.4 | 7.7 | 20.2 | 7.2 | 15.8 | 4.4 | 17.5 | 5.0 | 14.3 | 2.4 |
| PC 18:2_20:3 | 18.5 | 4.7 | 16.0 | 5.0 | 18.0 | 5.9 | 17.3 | 4.5 | 16.8 | 6.7 | 11.7 | 4.5 | 14.3 | 4.5 | 17.8 | 7.2 |
| PC 18:2_20:4 | 28.1 | 4.9 | 29.2 | 5.0 | 28.5 | 5.1 | 26.6 | 4.6 | 30.8 | 13.1 | 25.4 | 6.8 | 26.0 | 10.2 | 26.9 | 5.0 |
| PC total | 8215.1 | 1207.9 | 7944.1 | 302.2 | 7257.2 | 1219.7 | 6850.8 | 579.4 | 7408.5 | 1782.8 | 6224.1 | 522.4 | 7129.4 | 1604.6 | 6632.7 | 970.7 |
| PC O-16:1/16:0 (PC P-16:0/16:0) | 25.8 | 6.2 | 36.7 | 8.0 | 31.1 | 6.3 | 28.3 | 7.6 | 28.9 | 11.7 | 25.2 | 5.5 | 24.0 | 11.3 | 29.2 | 7.0 |
| PC O-16:1/18:1 (PC P-16:0/18:1) | 115.2 | 24.8 | 137.8 | 18.3 | 120.2 | 18.7 | 150.7 | 23.9 | 97.0 | 17.4 | 99.2 | 24.3 | 98.3 | 10.8 | 121.7 | 10.6 |
| PC O-16:1/18:2 (PC P-16:0/18:2) | 68.8 | 14.8 | 88.0 | 9.1 | 76.2 | 14.9 | 75.8 | 21.3 | 72.3 | 16.8 | 77.4 | 4.2 | 66.3 | 11.2 | 83.7 | 13.0 |
| PC O-16:1/20:4 (PC P-16:0/20:4) | 78.2 | 20.3 | 95.1 | 14.5 | 85.7 | 25.7 | 94.2 | 25.5 | 78.0 | 21.2 | 83.0 | 23.1 | 74.3 | 31.9 | 94.5 | 28.0 |
| PC P-16:1/20:5 | 25.1 | 5.1 | 28.8 | 7.1 | 27.2 | 5.9 | 34.2 | 13.3 | 21.9 | 11.0 | 26.5 | 4.1 | 22.4 | 7.9 | 21.3 | 9.6 |

| | | | | | | | | | | | | | | | | |
|---|---|---|---|---|---|---|---|---|---|---|---|---|---|---|---|---|
| PC O-18:1/18:1 (PC P-18:0/18:1) | 5.8 | 3.2 | 9.8 | 4.3 | 9.5 | 8.8 | 8.3 | 6.8 | 4.9 | 3.0 | 5.3 | 5.3 | 5.8 | 2.8 | 5.8 | 1.5 |
| PC O-18:1/18:2 (PC P-18:0/18:2) | 36.8 | 10.4 | 33.7 | 10.9 | 29.0 | 9.0 | 32.8 | 1.1 | 28.1 | 8.4 | 28.3 | 3.5 | 25.4 | 9.7 | 31.8 | 13.9 |
| PC O-18:1/20:4 (PC P-18:0/20:4) | 33.2 | 6.7 | 46.5 | 6.6 | 32.3 | 15.7 | 31.0 | 9.3 | 30.5 | 8.4 | 39.1 | 9.5 | 25.4 | 7.4 | 36.6 | 10.8 |
| PC O-18:2/18:2 (PC P-18:1/18:2) | 9.3 | 5.6 | 12.6 | 3.0 | 9.3 | 3.8 | 11.0 | 3.4 | 6.7 | 5.0 | 9.9 | 1.2 | 6.6 | 4.2 | 9.0 | 1.5 |
| PC O-18:2/20:4 (PC P-18:1/20:4) | 10.7 | 6.0 | 10.5 | 2.6 | 10.4 | 2.8 | 7.8 | 3.6 | 12.1 | 4.6 | 16.9 | 6.6 | 13.5 | 11.4 | 14.2 | 9.6 |
| PC P-18:2/20:5 | 306.6 | 52.4 | 308.8 | 49.3 | 299.0 | 65.4 | 304.8 | 65.8 | 329.3 | 81.7 | 263.2 | 42.5 | 296.2 | 62.4 | 328.5 | 128.3 |
| EtherPC total | 715.6 | 54.2 | 808.5 | 71.5 | 726.7 | 42.4 | 778.7 | 50.1 | 709.7 | 83.9 | 669.8 | 40.6 | 657.1 | 129.0 | 776.3 | 191.1 |
| PE 16:0_18:1 | 20.3 | 8.0 | 14.1 | 7.5 | 13.4 | 4.7 | 7.4 | 2.1 | 15.9 | 10.7 | 5.5 | 1.6 | 13.6 | 9.9 | 5.7 | 4.9 |
| PE 16:0_18:2 | 94.8 | 33.1 | 66.7 | 10.6 | 92.8 | 26.9 | 58.1 | 13.6 | 73.4 | 25.2 | 43.2 | 20.5 | 62.2 | 30.7 | 41.5 | 10.4 |
| PE 16:0_20:4 | 20.4 | 8.8 | 13.0 | 4.4 | 14.4 | 7.2 | 13.2 | 2.3 | 18.0 | 10.6 | 8.6 | 2.9 | 15.2 | 11.2 | 7.3 | 1.9 |
| PE 18:0_18:1 | 29.3 | 10.6 | 18.9 | 5.6 | 25.3 | 8.5 | 21.1 | 4.9 | 18.6 | 10.6 | 13.9 | 5.0 | 27.8 | 17.2 | 13.6 | 2.8 |
| PE 18:0_18:2 | 364.5 | 114.8 | 255.1 | 52.6 | 316.8 | 103.3 | 223.0 | 22.8 | 274.5 | 81.2 | 230.1 | 59.7 | 270.5 | 102.9 | 183.0 | 44.2 |
| PE 18:0_18:3 | 8.7 | 5.7 | 7.1 | 2.4 | 6.7 | 3.4 | 4.8 | 3.4 | 5.9 | 3.6 | 5.3 | 3.4 | 6.7 | 3.7 | 5.1 | 3.5 |
| PE 18:0_20:4 | 112.9 | 35.8 | 88.1 | 12.4 | 97.2 | 24.2 | 68.2 | 10.1 | 88.4 | 26.9 | 87.5 | 15.9 | 100.3 | 50.9 | 67.1 | 10.7 |
| PE 18:1_18:2 | 55.2 | 24.5 | 31.0 | 8.8 | 38.5 | 17.2 | 25.2 | 9.9 | 31.8 | 21.1 | 22.9 | 11.8 | 39.1 | 20.4 | 15.5 | 4.2 |
| PE 18:2_18:2 | 20.6 | 10.4 | 11.8 | 3.8 | 14.6 | 7.4 | 9.9 | 5.8 | 10.2 | 9.2 | 7.5 | 5.0 | 13.2 | 8.5 | 5.6 | 2.9 |
| PE total | 726.8 | 240.8 | 505.8 | 95.8 | 619.8 | 178.1 | 430.8 | 50.7 | 536.7 | 181.5 | 424.6 | 111.6 | 548.6 | 228.0 | 344.3 | 77.4 |
| PE P-16:0/16:0 | 2.1 | 1.2 | 3.0 | 2.3 | 2.3 | 1.7 | 3.3 | 2.8 | 1.0 | 0.6 | 2.1 | 1.0 | 1.3 | 1.2 | 1.5 | 1.2 |
| PE P-16:0/18:1 | 50.4 | 9.1 | 48.9 | 11.2 | 45.6 | 9.9 | 48.0 | 10.0 | 39.4 | 8.3 | 31.7 | 6.1 | 33.3 | 10.3 | 33.0 | 5.9 |
| PE P-16:0/18:2 | 276.0 | 52.1 | 295.1 | 75.6 | 245.7 | 55.7 | 298.1 | 57.6 | 226.6 | 31.4 | 214.7 | 28.9 | 168.1 | 43.2 | 172.0 | 37.1 |
| PE P-16:0/18:3 | 12.9 | 2.2 | 17.3 | 2.5 | 11.2 | 3.2 | 12.7 | 3.0 | 8.6 | 4.0 | 8.2 | 0.7 | 6.6 | 2.5 | 6.8 | 3.2 |
| PE P-16:0/20:3 | 5.2 | 2.1 | 5.8 | 1.6 | 4.5 | 0.9 | 3.5 | 1.0 | 4.5 | 4.2 | 3.7 | 0.7 | 5.1 | 4.0 | 4.8 | 1.6 |
| PE P-16:0/20:4 | 69.2 | 13.9 | 71.5 | 8.9 | 59.3 | 12.1 | 59.7 | 7.1 | 58.7 | 35.5 | 49.2 | 10.2 | 66.3 | 44.2 | 50.2 | 13.8 |
| PE P-16:0/20:5 | 1.4 | 0.7 | 0.9 | 0.2 | 0.6 | 0.5 | 1.2 | 0.6 | 1.2 | 0.9 | 0.8 | 0.6 | 1.2 | 1.3 | 0.9 | 0.8 |
| PE P-16:0/22:4 | 19.5 | 5.0 | 16.7 | 1.1 | 19.3 | 6.0 | 13.0 | 2.1 | 22.0 | 10.8 | 12.8 | 3.0 | 22.3 | 10.9 | 13.1 | 5.4 |

| | | | | | | | | | | | | | | | | |
|---|---|---|---|---|---|---|---|---|---|---|---|---|---|---|---|---|
| PE P-16:0/22:5 | 46.0 | 11.5 | 42.4 | 4.6 | 39.6 | 9.7 | 32.3 | 7.7 | 45.2 | 12.5 | 29.3 | 3.0 | 45.4 | 22.9 | 34.7 | 8.5 |
| PE P-16:0/22:6 | 19.4 | 3.2 | 16.5 | 5.2 | 16.1 | 4.2 | 12.1 | 3.0 | 14.8 | 4.6 | 10.0 | 0.7 | 18.3 | 10.5 | 9.7 | 3.2 |
| PE P-16:1/20:2 | 2.1 | 1.3 | 2.5 | 1.1 | 1.7 | 0.9 | 2.5 | 1.1 | 2.2 | 1.7 | 1.7 | 0.5 | 2.1 | 1.7 | 1.8 | 0.9 |
| PE P-16:1/22:2 | 4.0 | 1.2 | 5.0 | 1.7 | 3.6 | 1.6 | 2.8 | 1.2 | 4.7 | 3.2 | 2.9 | 1.5 | 4.8 | 3.7 | 4.1 | 2.1 |
| PE P-16:1/22:3 | 1.3 | 0.3 | 2.3 | 0.8 | 1.0 | 0.3 | 2.0 | 0.9 | 1.3 | 0.9 | 2.0 | 1.2 | 2.1 | 1.7 | 2.6 | 1.2 |
| PE P-18:0/16:0 | 3.9 | 1.1 | 5.4 | 2.1 | 4.4 | 2.4 | 5.5 | 1.3 | 2.5 | 1.7 | 2.3 | 0.8 | 2.5 | 1.8 | 3.1 | 0.9 |
| PE P-18:0/16:1 | 1.3 | 0.7 | 1.4 | 0.4 | 0.9 | 0.7 | 0.9 | 0.9 | 1.1 | 0.7 | 1.4 | 0.6 | 1.0 | 0.7 | 1.4 | 0.8 |
| PE P-18:0/18:0 | 0.9 | 0.5 | 1.3 | 0.8 | 1.0 | 0.7 | 1.1 | 1.1 | 1.1 | 0.4 | 1.5 | 1.4 | 0.9 | 0.3 | 0.6 | 0.5 |
| PE P-18:0/18:1 | 90.3 | 16.0 | 80.2 | 11.2 | 76.9 | 12.5 | 76.5 | 8.6 | 69.6 | 17.6 | 62.1 | 4.6 | 63.0 | 13.8 | 59.7 | 11.8 |
| PE P-18:0/18:2 | 859.2 | 81.1 | 839.2 | 72.8 | 734.2 | 86.3 | 716.3 | 86.0 | 669.1 | 80.2 | 594.4 | 21.9 | 563.2 | 105.1 | 556.5 | 55.1 |
| PE P-18:0/18:3 | 50.8 | 7.0 | 53.3 | 7.7 | 36.2 | 7.0 | 40.3 | 9.8 | 29.5 | 7.3 | 28.1 | 4.9 | 27.5 | 5.6 | 34.3 | 5.8 |
| PE P-18:0/20:3 | 9.2 | 1.9 | 10.1 | 1.8 | 8.2 | 2.0 | 8.6 | 1.9 | 8.9 | 4.5 | 7.1 | 1.1 | 7.9 | 5.0 | 7.3 | 5.6 |
| PE P-18:0/20:4 | 108.1 | 14.1 | 104.7 | 3.4 | 92.2 | 18.7 | 91.2 | 8.7 | 90.0 | 36.7 | 85.2 | 11.7 | 94.2 | 48.9 | 98.2 | 20.8 |
| PE P-18:0/20:5 | 3.4 | 1.4 | 3.5 | 1.9 | 1.5 | 1.1 | 2.4 | 0.8 | 3.0 | 2.1 | 1.2 | 0.7 | 2.0 | 1.8 | 1.5 | 0.8 |
| PE P-18:0/22:4 | 11.9 | 1.2 | 11.6 | 1.2 | 9.2 | 1.7 | 8.5 | 2.1 | 9.9 | 4.1 | 8.2 | 0.9 | 9.8 | 6.3 | 12.0 | 7.1 |
| PE P-18:0/22:5 | 29.6 | 2.6 | 30.7 | 4.7 | 25.5 | 3.1 | 26.7 | 9.5 | 26.6 | 7.5 | 21.6 | 2.7 | 27.6 | 17.9 | 25.3 | 16.2 |
| PE P-18:0/22:6 | 20.2 | 2.5 | 17.2 | 3.3 | 14.6 | 2.2 | 15.2 | 2.8 | 13.4 | 4.9 | 14.4 | 2.3 | 14.2 | 7.2 | 16.5 | 4.9 |
| PE P-18:1/16:0 | 12.7 | 3.9 | 13.0 | 5.2 | 10.4 | 4.1 | 14.4 | 4.2 | 10.0 | 2.7 | 10.3 | 3.1 | 9.0 | 3.7 | 10.9 | 6.8 |
| PE P-18:1/18:1 | 49.9 | 13.8 | 52.9 | 16.7 | 49.3 | 18.4 | 57.3 | 6.1 | 43.0 | 9.8 | 42.9 | 5.9 | 37.5 | 9.2 | 36.2 | 9.0 |
| PE P-18:1/18:2 | 305.6 | 81.9 | 332.5 | 99.3 | 309.8 | 110.5 | 344.4 | 53.0 | 283.6 | 46.1 | 280.0 | 34.6 | 217.2 | 38.5 | 237.8 | 71.1 |
| PE P-18:1/18:3 | 18.5 | 5.2 | 18.2 | 6.4 | 16.0 | 6.1 | 15.3 | 5.0 | 9.7 | 3.8 | 13.4 | 1.8 | 11.3 | 2.3 | 13.9 | 8.2 |
| PE P-18:1/20:2 | 1.4 | 0.5 | 1.1 | 0.4 | 1.3 | 0.4 | 0.9 | 0.3 | 0.8 | 0.6 | 1.0 | 0.7 | 1.5 | 0.6 | 0.3 | 0.2 |
| PE P-18:1/20:3 | 3.7 | 1.8 | 4.5 | 2.7 | 3.1 | 0.8 | 4.9 | 1.5 | 4.0 | 2.8 | 3.9 | 1.8 | 4.7 | 3.2 | 3.7 | 4.2 |
| PE P-18:1/20:4 | 48.7 | 8.7 | 56.2 | 15.2 | 47.1 | 6.0 | 47.1 | 4.9 | 44.3 | 19.0 | 48.5 | 6.0 | 55.1 | 31.8 | 54.4 | 13.5 |
| PE P-18:1/22:4 | 3.5 | 1.2 | 4.5 | 1.7 | 4.3 | 0.8 | 4.2 | 1.1 | 4.0 | 2.8 | 4.0 | 0.9 | 4.6 | 2.6 | 3.8 | 1.3 |

| | | | | | | | | | | | | | | | | |
|---|---|---|---|---|---|---|---|---|---|---|---|---|---|---|---|---|
| PE P-18:1/22:5 | 12.2 | 2.9 | 12.6 | 3.6 | 9.4 | 1.8 | 12.3 | 2.4 | 11.5 | 5.1 | 10.6 | 1.4 | 13.0 | 10.8 | 11.1 | 2.5 |
| PE P-18:1/22:6 | 5.9 | 2.0 | 6.0 | 1.7 | 5.1 | 1.1 | 4.5 | 1.7 | 3.9 | 1.7 | 6.4 | 2.0 | 5.2 | 4.9 | 7.2 | 2.7 |
| PE P-18:2/18:1 | 3.8 | 1.7 | 3.0 | 1.7 | 3.5 | 2.3 | 4.1 | 1.3 | 2.8 | 0.9 | 2.8 | 2.2 | 2.5 | 1.0 | 2.6 | 0.6 |
| PE P-18:2/18:2 | 27.4 | 11.5 | 23.2 | 7.4 | 22.1 | 10.6 | 26.8 | 2.5 | 23.8 | 2.0 | 26.6 | 8.0 | 17.8 | 2.4 | 15.9 | 5.3 |
| PE P-18:2/20:2 | 1.5 | 1.1 | 0.9 | 0.1 | 1.6 | 0.7 | 1.0 | 0.4 | 1.2 | 1.4 | 1.3 | 1.0 | 1.1 | 0.7 | 0.5 | 0.4 |
| PE P-18:2/20:4 | 3.6 | 1.8 | 3.3 | 1.1 | 2.6 | 1.5 | 1.7 | 0.9 | 2.9 | 1.2 | 3.8 | 2.7 | 4.2 | 4.5 | 2.9 | 2.0 |
| PE P-18:2/22:2 | 1.3 | 1.0 | 1.0 | 0.7 | 0.4 | 0.5 | 0.5 | 0.7 | 0.8 | 0.5 | 0.5 | 0.5 | 0.4 | 0.1 | 0.7 | 0.5 |
| EtherPE total | 2197.6 | 292.1 | 2218.9 | 337.3 | 1940.5 | 305.4 | 2023.6 | 157.1 | 1799.6 | 265.5 | 1651.8 | 107.2 | 1575.0 | 416.0 | 1553.2 | 258.5 |
| PI 16:0_16:0 | 441.5 | 269.8 | 476.1 | 53.8 | 219.7 | 60.8 | 315.5 | 70.9 | 354.2 | 223.5 | 157.1 | 175.6 | 338.6 | 236.7 | 243.3 | 280.6 |
| PI 16:0_18:1 | 3146.0 | 766.8 | 3477.1 | 529.1 | 2011.4 | 436.0 | 2353.1 | 116.5 | 2449.6 | 1234.6 | 2076.9 | 979.3 | 1964.8 | 1109.7 | 2276.5 | 1408.3 |
| PI 16:0_18:2 | 5162.7 | 915.4 | 6336.5 | 930.4 | 4588.8 | 110.1 | 4228.6 | 1182.3 | 3651.4 | 959.4 | 3561.7 | 1017.1 | 3216.9 | 737.1 | 3743.6 | 506.5 |
| PI 18:0_18:1 | 8527.3 | 1554.8 | 10457.1 | 1339.7 | 6978.2 | 2028.3 | 7341.3 | 531.0 | 8301.5 | 4383.3 | 6694.5 | 2666.6 | 6911.9 | 3274.8 | 6836.8 | 3611.9 |
| PI 18:0_18:2 | 23404.8 | 2245.0 | 27613.1 | 2602.5 | 20654.6 | 1445.7 | 21519.9 | 1425.0 | 18695.7 | 2877.8 | 19805.7 | 2358.8 | 17161.9 | 2874.7 | 17276.2 | 2749.8 |
| PI 18:0_20:2 | 2847.1 | 314.1 | 2084.3 | 811.9 | 2159.4 | 800.0 | 1557.9 | 588.0 | 2309.7 | 350.8 | 1540.1 | 907.8 | 1532.2 | 384.9 | 1389.1 | 649.1 |
| PI 18:0_20:3 | 2738.8 | 446.6 | 2106.6 | 380.3 | 2027.1 | 611.5 | 1835.1 | 286.0 | 1891.2 | 644.1 | 1660.4 | 568.9 | 1900.8 | 807.4 | 1822.9 | 796.4 |
| PI 18:0_20:4 | 8677.6 | 1352.5 | 8733.0 | 1193.6 | 7689.6 | 1914.3 | 7099.4 | 1604.6 | 7242.9 | 1176.5 | 7716.0 | 1018.7 | 7204.6 | 2108.4 | 7922.3 | 1738.5 |
| PI 18:1_18:1 | 445.9 | 177.5 | 712.7 | 316.2 | 360.6 | 229.5 | 458.1 | 118.2 | 528.7 | 361.7 | 490.5 | 238.2 | 487.5 | 328.5 | 426.3 | 329.7 |
| PI 18:1_18:2 | 721.4 | 125.9 | 899.7 | 231.2 | 592.0 | 276.8 | 605.3 | 211.9 | 603.1 | 248.5 | 680.2 | 202.6 | 675.9 | 308.1 | 652.7 | 351.5 |
| PI total | 56112.9 | 4711.7 | 62896.2 | 5648.2 | 47281.5 | 2253.1 | 47314.2 | 3035.5 | 46028.0 | 11374.2 | 44383.0 | 8902.7 | 41395.2 | 11126.0 | 42589.8 | 11336.5 |
| SM 18:1;O2/14:0 | 0.7 | 0.1 | 0.6 | 0.0 | 0.6 | 0.0 | 0.6 | 0.1 | 0.6 | 0.1 | 0.5 | 0.1 | 0.6 | 0.0 | 0.5 | 0.1 |
| SM 18:1;O2/14:1 | 0.0 | 0.0 | 0.0 | 0.0 | 0.0 | 0.0 | 0.0 | 0.0 | 0.0 | 0.0 | 0.0 | 0.0 | 0.0 | 0.0 | 0.0 | 0.0 |
| SM 18:1;O2/16:0 | 61.1 | 6.1 | 56.2 | 4.6 | 57.1 | 2.9 | 56.1 | 3.4 | 52.5 | 7.6 | 53.5 | 3.8 | 56.6 | 6.4 | 54.8 | 6.6 |
| SM 18:1;O2/16:1 | 3.7 | 0.4 | 3.4 | 0.3 | 3.3 | 0.4 | 3.3 | 0.4 | 3.1 | 0.4 | 3.2 | 0.2 | 3.3 | 0.4 | 3.2 | 0.6 |
| SM 18:1;O2/18:0 | 13.1 | 1.7 | 11.6 | 0.9 | 11.9 | 0.8 | 11.2 | 1.6 | 11.2 | 1.1 | 11.0 | 0.4 | 12.2 | 0.7 | 11.3 | 1.2 |
| SM 18:1;O2/18:1 | 3.9 | 0.5 | 3.4 | 0.3 | 3.5 | 0.3 | 3.5 | 0.2 | 3.2 | 0.1 | 3.3 | 0.1 | 3.0 | 0.3 | 3.2 | 0.2 |

| | | | | | | | | | | | | | | | | |
|---|---|---|---|---|---|---|---|---|---|---|---|---|---|---|---|---|
| SM 18:1;O2/18:2 | 0.0 | 0.0 | 0.1 | 0.0 | 0.0 | 0.0 | 0.1 | 0.0 | 0.0 | 0.0 | 0.0 | 0.0 | 0.0 | 0.0 | 0.1 | 0.0 |
| SM 18:1;O2/20:0 | 4.6 | 0.7 | 4.4 | 0.4 | 4.3 | 0.5 | 4.3 | 0.4 | 4.1 | 0.5 | 3.9 | 0.1 | 4.2 | 0.5 | 4.0 | 0.2 |
| SM 18:1;O2/20:1 | 2.6 | 0.3 | 2.8 | 0.3 | 2.5 | 0.2 | 2.7 | 0.3 | 2.4 | 0.3 | 2.6 | 0.1 | 2.2 | 0.3 | 2.4 | 0.2 |
| SM 18:1;O2/20:2 | 0.1 | 0.0 | 0.1 | 0.0 | 0.1 | 0.0 | 0.1 | 0.0 | 0.1 | 0.0 | 0.1 | 0.0 | 0.1 | 0.0 | 0.1 | 0.0 |
| SM 18:1;O2/22:0 | 5.5 | 0.8 | 5.5 | 0.8 | 5.1 | 0.6 | 5.2 | 0.5 | 4.9 | 0.8 | 4.8 | 0.2 | 5.0 | 0.4 | 4.7 | 0.5 |
| SM 18:1;O2/22:1 | 3.5 | 0.4 | 3.7 | 0.3 | 3.3 | 0.2 | 3.7 | 0.3 | 3.1 | 0.4 | 3.4 | 0.1 | 3.1 | 0.3 | 3.2 | 0.3 |
| SM 18:1;O2/22:2 | 0.8 | 0.1 | 0.8 | 0.2 | 0.8 | 0.1 | 0.7 | 0.1 | 0.7 | 0.1 | 0.7 | 0.1 | 0.8 | 0.1 | 0.7 | 0.1 |
| SM 18:1;O2/22:3 | 0.2 | 0.0 | 0.2 | 0.0 | 0.1 | 0.0 | 0.1 | 0.0 | 0.1 | 0.0 | 0.1 | 0.0 | 0.2 | 0.0 | 0.2 | 0.0 |
| SM total | 99.7 | 10.8 | 92.8 | 7.8 | 92.7 | 5.0 | 91.7 | 6.5 | 85.9 | 8.6 | 87.4 | 4.5 | 91.2 | 8.0 | 88.2 | 9.3 |
| Cer 18:1;O2/16:0 | 5.0 | 0.7 | 4.4 | 1.2 | 4.5 | 0.5 | 3.3 | 0.6 | 4.0 | 0.8 | 3.4 | 0.3 | 3.7 | 1.1 | 2.9 | 0.4 |
| Cer 18:1;O2/18:0 | 5.9 | 1.6 | 3.5 | 0.6 | 4.7 | 1.2 | 3.0 | 0.5 | 4.6 | 0.9 | 3.3 | 0.4 | 4.8 | 1.2 | 3.0 | 0.6 |
| Cer 18:1;O2/18:2 | 2.2 | 0.5 | 1.4 | 0.3 | 2.0 | 0.3 | 1.4 | 0.2 | 1.9 | 0.3 | 1.5 | 0.4 | 1.6 | 0.4 | 1.3 | 0.4 |
| Cer 18:1;O2/18:4 | 0.1 | 0.1 | 0.1 | 0.0 | 0.1 | 0.0 | 0.1 | 0.0 | 0.1 | 0.0 | 0.1 | 0.0 | 0.1 | 0.0 | 0.1 | 0.0 |
| Cer 18:1;O2/20:0 | 6.8 | 1.2 | 4.5 | 0.3 | 5.9 | 1.2 | 4.4 | 1.2 | 5.5 | 1.4 | 4.6 | 1.3 | 4.9 | 1.2 | 3.7 | 1.0 |
| Cer 18:1;O2/20:1 | 0.1 | 0.0 | 0.1 | 0.1 | 0.1 | 0.0 | 0.1 | 0.0 | 0.1 | 0.0 | 0.1 | 0.0 | 0.1 | 0.0 | 0.1 | 0.0 |
| Cer 18:1;O2/20:2 | 4.2 | 0.9 | 3.0 | 0.3 | 3.7 | 0.7 | 2.9 | 0.6 | 3.4 | 0.9 | 2.8 | 0.6 | 2.7 | 0.7 | 2.1 | 0.4 |
| Cer 18:1;O2/20:4 | 0.5 | 0.1 | 0.3 | 0.1 | 0.5 | 0.1 | 0.4 | 0.1 | 0.5 | 0.1 | 0.4 | 0.1 | 0.4 | 0.1 | 0.3 | 0.1 |
| Cer 18:1;O2/22:0 | 23.3 | 6.2 | 15.2 | 2.8 | 20.5 | 4.5 | 16.5 | 3.2 | 18.2 | 3.6 | 15.9 | 2.8 | 17.2 | 2.7 | 14.7 | 4.2 |
| Cer 18:1;O2/22:1 | 2.1 | 0.5 | 1.6 | 0.1 | 2.0 | 0.4 | 1.7 | 0.3 | 1.8 | 0.5 | 1.6 | 0.3 | 1.5 | 0.3 | 1.3 | 0.4 |
| Cer 18:1;O2/22:2 | 56.9 | 13.7 | 41.2 | 3.0 | 52.3 | 9.6 | 42.3 | 6.3 | 49.8 | 12.6 | 41.4 | 6.1 | 37.7 | 4.5 | 34.0 | 6.6 |
| Cer 18:1;O2/22:3 | 0.5 | 0.1 | 0.4 | 0.1 | 0.4 | 0.1 | 0.3 | 0.1 | 0.4 | 0.2 | 0.4 | 0.1 | 0.4 | 0.2 | 0.4 | 0.1 |
| Cer 18:1;O2/22:4 | 0.6 | 0.1 | 0.4 | 0.1 | 0.6 | 0.2 | 0.5 | 0.1 | 0.5 | 0.2 | 0.4 | 0.1 | 0.4 | 0.1 | 0.4 | 0.1 |
| Cer 18:1;O2/22:5 | 0.3 | 0.1 | 0.2 | 0.0 | 0.2 | 0.1 | 0.2 | 0.0 | 0.2 | 0.1 | 0.1 | 0.0 | 0.2 | 0.1 | 0.2 | 0.1 |
| Cer total | 108.5 | 25.2 | 76.2 | 7.7 | 97.5 | 17.6 | 77.0 | 12.2 | 91.1 | 20.3 | 76.1 | 11.8 | 75.7 | 10.9 | 64.5 | 13.7 |
| CE 14:0 | 46.7 | 13.0 | 43.3 | 11.7 | 33.3 | 9.3 | 34.3 | 4.7 | 31.1 | 11.4 | 31.5 | 5.1 | 33.5 | 10.3 | 28.4 | 12.6 |

| | | | | | | | | | | | | | | | | |
|---|---|---|---|---|---|---|---|---|---|---|---|---|---|---|---|---|
| CE 14:1 | 18.5 | 8.6 | 23.2 | 4.0 | 12.5 | 7.7 | 19.1 | 3.5 | 12.5 | 7.1 | 18.1 | 3.8 | 11.9 | 6.5 | 18.4 | 6.3 |
| CE 16:0 | 3693.2 | 428.3 | 3286.5 | 554.9 | 3314.4 | 280.1 | 3037.1 | 257.0 | 2887.0 | 638.3 | 2877.2 | 304.1 | 2749.1 | 366.2 | 2896.8 | 1113.3 |
| CE 16:1 | 3222.9 | 914.5 | 3881.1 | 985.9 | 3061.2 | 1039.0 | 3382.6 | 299.7 | 3271.0 | 1350.8 | 3322.4 | 493.7 | 3363.7 | 1619.9 | 4434.6 | 1127.5 |
| CE 18:0 | 1567.1 | 300.9 | 1833.5 | 319.6 | 1310.8 | 387.1 | 1131.4 | 261.8 | 1408.4 | 323.2 | 1057.2 | 214.8 | 1850.4 | 629.6 | 1502.2 | 718.1 |
| CE 18:1 | 33861.3 | 9580.5 | 39842.2 | 11016.5 | 26384.0 | 9119.3 | 23877.4 | 4584.8 | 29724.0 | 9077.7 | 20301.7 | 5001.1 | 36167.1 | 10473.0 | 29338.5 | 11009.9 |
| CE 18:2 | 361715.0 | 154527.8 | 370462.1 | 134852.8 | 241580.7 | 65500.7 | 220508.9 | 12184.9 | 238156.1 | 61444.1 | 193811.9 | 27744.5 | 304717.5 | 103733.7 | 216251.9 | 49926.2 |
| CE 18:3 | 61766.3 | 24738.4 | 60153.1 | 18757.1 | 38088.3 | 10527.2 | 34595.5 | 3745.4 | 34207.5 | 11996.4 | 30460.2 | 1998.3 | 44433.2 | 15006.1 | 37973.7 | 9532.7 |
| CE 18:4 | 233.6 | 69.3 | 231.6 | 66.8 | 145.1 | 30.9 | 139.7 | 38.4 | 125.7 | 47.5 | 105.8 | 12.2 | 255.7 | 277.8 | 170.5 | 143.9 |
| CE 20:1 | 124.2 | 67.7 | 125.4 | 54.3 | 83.1 | 42.0 | 70.4 | 12.7 | 91.8 | 39.0 | 65.3 | 23.4 | 103.4 | 45.1 | 79.9 | 57.6 |
| CE 20:2 | 598.4 | 333.4 | 479.7 | 219.0 | 355.2 | 132.3 | 291.8 | 41.7 | 413.4 | 144.9 | 309.3 | 112.8 | 530.8 | 274.1 | 354.5 | 189.0 |
| CE 20:3 | 2536.7 | 1201.4 | 2307.6 | 817.1 | 1470.8 | 321.6 | 1335.0 | 193.9 | 1431.6 | 398.6 | 1209.4 | 282.6 | 1986.5 | 1048.2 | 1525.4 | 863.1 |
| CE 20:4 | 19404.3 | 8205.7 | 20266.5 | 5600.3 | 13024.9 | 1615.7 | 12323.1 | 2412.5 | 11406.4 | 2317.6 | 10790.9 | 1182.7 | 17355.2 | 9312.0 | 14254.9 | 5067.1 |
| CE 20:5 | 1155.4 | 567.5 | 1269.7 | 534.9 | 614.7 | 194.0 | 616.5 | 115.3 | 519.6 | 237.1 | 452.6 | 46.8 | 718.0 | 519.7 | 492.4 | 280.0 |
| CE 22:1 | 21.8 | 12.5 | 21.8 | 11.3 | 10.7 | 5.6 | 11.0 | 1.7 | 10.1 | 4.9 | 7.6 | 1.5 | 13.7 | 6.2 | 8.6 | 2.8 |
| CE 22:2 | 19.5 | 8.8 | 21.1 | 5.4 | 11.7 | 4.3 | 12.2 | 1.5 | 12.9 | 5.4 | 10.6 | 2.2 | 16.9 | 10.1 | 11.8 | 6.7 |
| CE 22:3 | 21.6 | 9.9 | 26.2 | 12.0 | 12.4 | 2.8 | 13.2 | 3.4 | 12.6 | 4.9 | 10.4 | 1.8 | 22.1 | 13.2 | 16.8 | 9.7 |
| CE 22:4 | 140.4 | 63.3 | 156.5 | 55.5 | 87.6 | 14.7 | 89.3 | 18.9 | 84.9 | 22.3 | 76.6 | 5.8 | 134.2 | 78.0 | 101.7 | 47.2 |
| CE 22:5 | 301.1 | 137.3 | 334.9 | 104.8 | 203.0 | 45.6 | 198.1 | 46.9 | 175.5 | 36.5 | 177.0 | 12.5 | 286.0 | 173.2 | 260.9 | 148.7 |
| CE 22:6 | 352.7 | 187.3 | 299.5 | 115.7 | 204.2 | 36.2 | 181.1 | 41.3 | 197.0 | 56.5 | 184.3 | 28.8 | 270.8 | 146.6 | 263.9 | 120.5 |
| CE total | 490800.6 | 198111.3 | 505065.4 | 172729.8 | 330008.7 | 86410.1 | 301867.8 | 13885.1 | 324179.1 | 84001.2 | 265279.9 | 35752.0 | 415019.7 | 139400.9 | 309985.8 | 78857.6 |
| MG 16:0 | 7.6 | 5.3 | 7.5 | 3.5 | 6.3 | 1.8 | 5.2 | 1.3 | 6.0 | 1.0 | 5.9 | 0.4 | 6.2 | 1.9 | 5.5 | 0.4 |
| MG 18:0 | 48.2 | 71.5 | 27.8 | 20.7 | 12.6 | 3.6 | 14.4 | 7.8 | 12.2 | 3.6 | 9.4 | 2.0 | 14.9 | 8.6 | 12.4 | 1.5 |
| MG total | 55.8 | 76.8 | 35.3 | 22.4 | 18.8 | 2.9 | 19.5 | 8.6 | 18.3 | 4.4 | 15.3 | 2.3 | 21.1 | 8.8 | 17.9 | 1.9 |
| DG 14:0_16:0 | 0.1 | 0.0 | 0.1 | 0.0 | 0.1 | 0.1 | 0.1 | 0.0 | 0.1 | 0.0 | 0.1 | 0.0 | 0.1 | 0.0 | 0.1 | 0.1 |
| DG 14:0_16:1 | 0.0 | 0.0 | 0.0 | 0.0 | 0.0 | 0.0 | 0.0 | 0.0 | 0.0 | 0.0 | 0.0 | 0.0 | 0.0 | 0.0 | 0.0 | 0.0 |

| | | | | | | | | | | | | | | | | |
|---|---|---|---|---|---|---|---|---|---|---|---|---|---|---|---|---|
| DG 14:0_18:0 | 0.0 | 0.0 | 0.0 | 0.0 | 0.0 | 0.0 | 0.0 | 0.0 | 0.0 | 0.0 | 0.0 | 0.0 | 0.0 | 0.0 | 0.0 | 0.0 |
| DG 14:0_18:1 | 0.4 | 0.2 | 0.4 | 0.2 | 0.4 | 0.3 | 0.3 | 0.0 | 0.4 | 0.3 | 0.3 | 0.1 | 0.4 | 0.2 | 0.3 | 0.1 |
| DG 14:0_18:2 | 0.4 | 0.2 | 0.3 | 0.1 | 0.4 | 0.3 | 0.4 | 0.1 | 0.4 | 0.2 | 0.4 | 0.1 | 0.5 | 0.3 | 0.4 | 0.1 |
| DG 14:0_18:3 | 0.1 | 0.0 | 0.1 | 0.0 | 0.1 | 0.1 | 0.1 | 0.0 | 0.1 | 0.1 | 0.1 | 0.0 | 0.1 | 0.0 | 0.1 | 0.0 |
| DG 14:0_20:0 | 0.0 | 0.0 | 0.0 | 0.0 | 0.0 | 0.0 | 0.0 | 0.0 | 0.0 | 0.0 | 0.0 | 0.0 | 0.0 | 0.0 | 0.0 | 0.0 |
| DG 14:1_18:1 | 0.0 | 0.0 | 0.0 | 0.0 | 0.0 | 0.0 | 0.1 | 0.0 | 0.0 | 0.0 | 0.0 | 0.0 | 0.0 | 0.0 | 0.0 | 0.0 |
| DG 14:1_18:2 | 0.0 | 0.0 | 0.0 | 0.0 | 0.1 | 0.1 | 0.1 | 0.0 | 0.1 | 0.1 | 0.1 | 0.0 | 0.1 | 0.1 | 0.1 | 0.0 |
| DG 14:1_22:2 | 0.0 | 0.0 | 0.0 | 0.0 | 0.0 | 0.0 | 0.0 | 0.0 | 0.1 | 0.0 | 0.0 | 0.0 | 0.1 | 0.0 | 0.0 | 0.0 |
| DG 16:0_16:0 | 0.6 | 0.2 | 0.6 | 0.1 | 0.6 | 0.2 | 0.5 | 0.1 | 0.7 | 0.3 | 0.5 | 0.2 | 0.8 | 0.5 | 0.6 | 0.2 |
| DG 16:0_16:1 | 0.6 | 0.3 | 0.5 | 0.1 | 0.6 | 0.2 | 0.6 | 0.1 | 0.7 | 0.4 | 0.8 | 0.2 | 0.9 | 0.8 | 0.8 | 0.3 |
| DG 16:0_18:0 | 0.4 | 0.2 | 0.4 | 0.1 | 0.4 | 0.1 | 0.4 | 0.0 | 0.4 | 0.2 | 0.3 | 0.1 | 0.5 | 0.3 | 0.3 | 0.1 |
| DG 16:0_18:1 | 5.6 | 2.4 | 4.2 | 0.8 | 5.3 | 1.5 | 4.3 | 0.4 | 5.6 | 2.5 | 4.2 | 1.6 | 6.5 | 3.5 | 4.6 | 1.3 |
| DG 16:0_18:2 | 8.6 | 2.5 | 6.5 | 0.7 | 9.6 | 2.7 | 7.8 | 1.0 | 10.5 | 3.3 | 8.6 | 1.9 | 12.7 | 6.9 | 9.2 | 1.9 |
| DG 16:0_18:3 | 1.4 | 0.4 | 1.3 | 0.1 | 1.4 | 0.7 | 1.4 | 0.1 | 1.4 | 0.5 | 1.4 | 0.4 | 1.7 | 1.0 | 1.6 | 0.4 |
| DG 16:0_20:2 | 0.1 | 0.0 | 0.1 | 0.0 | 0.1 | 0.1 | 0.1 | 0.1 | 0.1 | 0.1 | 0.1 | 0.1 | 0.1 | 0.1 | 0.1 | 0.0 |
| DG 16:0_20:3 | 0.1 | 0.0 | 0.1 | 0.1 | 0.1 | 0.1 | 0.1 | 0.0 | 0.2 | 0.1 | 0.1 | 0.1 | 0.2 | 0.2 | 0.2 | 0.1 |
| DG 16:0_20:4 | 0.2 | 0.0 | 0.2 | 0.0 | 0.2 | 0.0 | 0.2 | 0.0 | 0.2 | 0.1 | 0.2 | 0.0 | 0.4 | 0.3 | 0.3 | 0.1 |
| DG 16:0_22:4 | 0.1 | 0.0 | 0.1 | 0.0 | 0.0 | 0.0 | 0.1 | 0.0 | 0.1 | 0.1 | 0.1 | 0.0 | 0.2 | 0.2 | 0.1 | 0.1 |
| DG 16:0_22:5 | 0.1 | 0.0 | 0.0 | 0.0 | 0.1 | 0.0 | 0.0 | 0.0 | 0.1 | 0.0 | 0.1 | 0.0 | 0.1 | 0.2 | 0.1 | 0.1 |
| DG 16:1_16:1 | 0.1 | 0.0 | 0.1 | 0.0 | 0.1 | 0.0 | 0.1 | 0.0 | 0.1 | 0.0 | 0.1 | 0.0 | 0.1 | 0.1 | 0.1 | 0.0 |
| DG 16:1_18:0 | 0.1 | 0.1 | 0.1 | 0.0 | 0.1 | 0.1 | 0.1 | 0.0 | 0.1 | 0.1 | 0.1 | 0.1 | 0.2 | 0.1 | 0.1 | 0.0 |
| DG 16:1_18:1 | 1.4 | 0.7 | 1.2 | 0.2 | 1.5 | 0.7 | 1.5 | 0.2 | 1.7 | 1.0 | 1.8 | 0.7 | 1.8 | 1.2 | 1.9 | 0.5 |
| DG 16:1_18:2 | 2.1 | 0.8 | 1.9 | 0.4 | 2.3 | 0.9 | 2.7 | 0.3 | 2.9 | 1.4 | 3.6 | 0.8 | 3.3 | 1.9 | 3.7 | 0.6 |
| DG 16:1_18:3 | 0.2 | 0.1 | 0.3 | 0.1 | 0.3 | 0.2 | 0.3 | 0.0 | 0.3 | 0.2 | 0.5 | 0.1 | 0.3 | 0.2 | 0.4 | 0.1 |
| DG 16:1_20:2 | 0.1 | 0.0 | 0.1 | 0.0 | 0.1 | 0.1 | 0.1 | 0.0 | 0.1 | 0.0 | 0.1 | 0.0 | 0.1 | 0.1 | 0.1 | 0.0 |

| | | | | | | | | | | | | | | | | |
|---|---|---|---|---|---|---|---|---|---|---|---|---|---|---|---|---|
| DG 16:1_20:3 | 0.1 | 0.0 | 0.0 | 0.0 | 0.1 | 0.0 | 0.1 | 0.0 | 0.1 | 0.0 | 0.1 | 0.0 | 0.1 | 0.1 | 0.1 | 0.0 |
| DG 16:1_20:4 | 0.0 | 0.0 | 0.1 | 0.0 | 0.0 | 0.0 | 0.1 | 0.0 | 0.1 | 0.0 | 0.1 | 0.0 | 0.1 | 0.1 | 0.1 | 0.0 |
| DG 18:0_18:0 | 0.1 | 0.0 | 0.1 | 0.0 | 0.1 | 0.0 | 0.0 | 0.0 | 0.1 | 0.0 | 0.0 | 0.0 | 0.1 | 0.0 | 0.1 | 0.0 |
| DG 18:0_18:1 | 1.5 | 0.5 | 1.3 | 0.4 | 1.6 | 0.5 | 1.2 | 0.1 | 1.7 | 0.8 | 1.3 | 0.5 | 2.0 | 1.1 | 1.3 | 0.2 |
| DG 18:0_18:2 | 3.4 | 0.9 | 3.0 | 0.3 | 3.6 | 1.1 | 3.4 | 0.5 | 3.9 | 1.1 | 3.2 | 0.6 | 4.9 | 2.6 | 3.4 | 0.7 |
| DG 18:0_18:3 | 0.3 | 0.1 | 0.4 | 0.1 | 0.3 | 0.1 | 0.3 | 0.0 | 0.3 | 0.1 | 0.3 | 0.0 | 0.4 | 0.2 | 0.4 | 0.1 |
| DG 18:0_20:4 | 0.1 | 0.0 | 0.1 | 0.0 | 0.1 | 0.0 | 0.1 | 0.0 | 0.1 | 0.0 | 0.1 | 0.0 | 0.1 | 0.1 | 0.1 | 0.0 |
| DG 18:1_18:1 | 8.4 | 3.9 | 6.7 | 1.9 | 8.9 | 3.6 | 7.8 | 1.4 | 10.3 | 5.0 | 8.1 | 3.0 | 10.2 | 4.9 | 7.6 | 2.3 |
| DG 18:1_18:2 | 18.2 | 7.3 | 13.9 | 3.4 | 20.1 | 6.7 | 18.2 | 2.9 | 22.3 | 8.8 | 20.5 | 5.2 | 25.1 | 11.5 | 21.8 | 2.9 |
| DG 18:1_18:3 | 2.4 | 1.0 | 2.0 | 0.4 | 2.0 | 0.8 | 2.0 | 0.4 | 2.2 | 1.1 | 2.2 | 0.5 | 2.6 | 1.6 | 2.4 | 0.5 |
| DG 18:1_18:4 | 0.0 | 0.0 | 0.0 | 0.0 | 0.0 | 0.0 | 0.0 | 0.0 | 0.0 | 0.0 | 0.0 | 0.0 | 0.1 | 0.1 | 0.0 | 0.0 |
| DG 18:1_20:2 | 0.2 | 0.1 | 0.2 | 0.1 | 0.2 | 0.1 | 0.1 | 0.1 | 0.2 | 0.1 | 0.1 | 0.1 | 0.3 | 0.2 | 0.2 | 0.1 |
| DG 18:1_20:3 | 0.3 | 0.1 | 0.2 | 0.0 | 0.2 | 0.1 | 0.2 | 0.1 | 0.2 | 0.2 | 0.2 | 0.1 | 0.4 | 0.3 | 0.2 | 0.1 |
| DG 18:1_20:4 | 0.4 | 0.2 | 0.3 | 0.1 | 0.3 | 0.1 | 0.3 | 0.0 | 0.4 | 0.2 | 0.3 | 0.0 | 0.6 | 0.6 | 0.4 | 0.2 |
| DG 18:1_22:4 | 0.0 | 0.0 | 0.0 | 0.0 | 0.0 | 0.0 | 0.0 | 0.0 | 0.0 | 0.0 | 0.0 | 0.0 | 0.0 | 0.1 | 0.0 | 0.0 |
| DG 18:1_22:5 | 0.0 | 0.0 | 0.0 | 0.0 | 0.0 | 0.0 | 0.0 | 0.0 | 0.0 | 0.0 | 0.0 | 0.0 | 0.1 | 0.1 | 0.1 | 0.0 |
| DG 18:2_18:2 | 5.8 | 2.2 | 4.7 | 1.3 | 5.9 | 1.9 | 5.2 | 0.9 | 6.4 | 2.7 | 6.3 | 1.3 | 8.1 | 5.1 | 7.1 | 1.4 |
| DG 18:2_18:3 | 0.9 | 0.3 | 0.8 | 0.2 | 0.9 | 0.5 | 0.9 | 0.2 | 0.8 | 0.4 | 1.0 | 0.1 | 1.1 | 0.8 | 1.0 | 0.2 |
| DG 18:2_20:2 | 0.1 | 0.0 | 0.1 | 0.0 | 0.1 | 0.1 | 0.1 | 0.0 | 0.1 | 0.1 | 0.1 | 0.0 | 0.2 | 0.2 | 0.1 | 0.0 |
| DG 18:2_20:3 | 0.1 | 0.0 | 0.1 | 0.0 | 0.1 | 0.1 | 0.1 | 0.0 | 0.1 | 0.1 | 0.1 | 0.0 | 0.2 | 0.2 | 0.2 | 0.1 |
| DG 18:2_20:4 | 0.2 | 0.1 | 0.2 | 0.0 | 0.2 | 0.1 | 0.3 | 0.0 | 0.2 | 0.1 | 0.3 | 0.1 | 0.4 | 0.4 | 0.3 | 0.1 |
| DG 18:2_22:4 | 0.0 | 0.0 | 0.0 | 0.0 | 0.0 | 0.0 | 0.0 | 0.0 | 0.0 | 0.0 | 0.0 | 0.0 | 0.0 | 0.1 | 0.0 | 0.0 |
| DG 18:3_18:3 | 0.0 | 0.0 | 0.0 | 0.0 | 0.0 | 0.0 | 0.0 | 0.0 | 0.0 | 0.0 | 0.0 | 0.0 | 0.0 | 0.0 | 0.1 | 0.0 |
| DG 18:3_20:4 | 0.0 | 0.0 | 0.0 | 0.0 | 0.0 | 0.0 | 0.0 | 0.0 | 0.0 | 0.0 | 0.0 | 0.0 | 0.0 | 0.0 | 0.0 | 0.0 |
| DG total | 65.7 | 25.0 | 52.9 | 9.0 | 68.9 | 22.1 | 62.1 | 8.0 | 76.1 | 30.6 | 68.2 | 17.8 | 88.6 | 47.9 | 72.4 | 13.6 |

Lipidomics data were normalized using the ion abundance of internal standards added before lipid extraction via the modified Bligh and Dyer method.

### Supplementary Table 6 Quantitative lipidomics for VLDL fraction.

| Lipids | Quantification (nmol $mL^{-1}$) | | | | | | | | | | | | | | | |
|---|---|---|---|---|---|---|---|---|---|---|---|---|---|---|---|---|
| | 8 months | | | | 12 months | | | | 16 months | | | | 20 months | | | |
| | Severe | | Mild | | Severe | | Mild | | Severe | | Mild | | Severe | | Mild | |
| | Mean | SD | Mean | SD | Mean | SD | Mean | SD | Mean | SD | Mean | SD | Mean | SD | Mean | SD |
| FA 14:0 | 5.9 | 4.5 | 7.1 | 5.4 | 7.8 | 5.9 | 10.3 | 2.1 | 9.0 | 3.1 | 7.4 | 5.0 | 4.3 | 3.4 | 5.2 | 3.1 |
| FA 16:0 | 72.8 | 44.3 | 79.2 | 9.7 | 79.8 | 44.8 | 69.2 | 20.5 | 47.5 | 26.3 | 56.1 | 6.2 | 45.6 | 26.9 | 62.1 | 12.9 |
| FA 16:1 | 2.4 | 1.0 | 3.6 | 0.7 | 3.5 | 2.3 | 3.7 | 1.8 | 1.9 | 1.5 | 2.2 | 0.7 | 3.4 | 0.7 | 2.1 | 1.4 |
| FA 18:0 | 83.7 | 32.3 | 81.3 | 15.2 | 80.0 | 12.2 | 74.9 | 8.7 | 74.0 | 11.5 | 61.6 | 5.0 | 54.4 | 7.6 | 59.5 | 13.2 |
| FA 18:1 | 6.0 | 3.1 | 6.1 | 2.5 | 10.9 | 5.6 | 8.5 | 1.7 | 3.9 | 3.2 | 5.0 | 2.3 | 8.3 | 4.1 | 6.2 | 1.9 |
| FA 18:2 | 0.8 | 0.5 | 0.8 | 0.7 | 1.9 | 1.3 | 1.2 | 0.9 | 0.7 | 0.4 | 0.3 | 0.4 | 1.3 | 1.1 | 0.9 | 0.5 |
| FA 18:3 | 0.8 | 0.4 | 0.8 | 0.7 | 0.6 | 0.4 | 0.8 | 0.7 | 0.6 | 0.6 | 0.4 | 0.2 | 0.3 | 0.3 | 0.4 | 0.3 |
| FA 20:4 | 0.1 | 0.1 | 0.4 | 0.5 | 0.2 | 0.2 | 0.2 | 0.1 | 0.5 | 0.3 | 0.1 | 0.1 | 0.1 | 0.2 | 0.1 | 0.1 |
| FA total | 172.4 | 68.7 | 179.3 | 9.1 | 184.6 | 52.1 | 168.7 | 31.2 | 138.1 | 28.0 | 133.0 | 8.9 | 117.8 | 34.2 | 136.3 | 18.8 |
| LPC 14:0 | n.d. | n.d. | 0.1 | n.d. | n.d. | n.d. | n.d. | n.d. | n.d. | n.d. | n.d. | n.d. | 0.2 | n.d. | n.d. | n.d. |
| LPC 16:0 | 5.8 | 3.7 | 6.0 | 2.9 | 5.5 | 4.2 | 3.9 | 1.1 | 8.7 | 6.9 | 5.9 | 0.8 | 10.9 | 18.1 | 7.4 | 4.1 |
| LPC 16:1 | 0.0 | 0.0 | 0.0 | n.d. | 0.0 | 0.0 | 0.0 | n.d. | 0.1 | 0.1 | 0.1 | n.d. | 0.4 | n.d. | 0.1 | 0.0 |
| LPC 18:0 | 13.9 | 7.5 | 14.3 | 6.0 | 14.1 | 11.1 | 12.2 | 2.5 | 20.5 | 13.6 | 16.1 | 1.6 | 18.5 | 23.4 | 17.6 | 9.9 |
| LPC 18:1 | 1.7 | 1.1 | 1.6 | 1.0 | 2.1 | 2.0 | 1.3 | 0.2 | 3.0 | 2.4 | 1.8 | 0.1 | 2.8 | 4.1 | 2.0 | 1.2 |
| LPC 18:2 | 2.5 | 1.3 | 2.1 | 0.7 | 2.9 | 2.5 | 2.0 | 0.4 | 4.4 | 3.6 | 2.9 | 0.7 | 3.2 | 4.2 | 3.0 | 2.1 |
| LPC 18:3 | 0.0 | 0.0 | 0.0 | n.d. | 0.1 | 0.0 | 0.0 | 0.0 | 0.2 | 0.0 | 0.1 | n.d. | 0.1 | 0.0 | 0.1 | n.d. |
| LPC 20:0 | 0.1 | 0.0 | 0.0 | 0.0 | 0.1 | 0.0 | 0.1 | 0.0 | 0.1 | 0.1 | 0.1 | 0.0 | 0.1 | 0.2 | 0.1 | 0.0 |
| LPC 20:1 | 0.0 | 0.0 | 0.1 | n.d. | 0.1 | 0.0 | n.d. | n.d. | 0.1 | 0.1 | 0.0 | 0.0 | 0.3 | n.d. | 0.0 | 0.0 |
| LPC 20:2 | 0.0 | 0.0 | 0.1 | 0.2 | 0.1 | 0.1 | 0.1 | 0.1 | 0.2 | 0.2 | 0.1 | n.d. | 0.1 | n.d. | 0.1 | 0.1 |
| LPC 20:3 | 0.1 | 0.0 | 0.0 | 0.0 | 0.1 | 0.0 | 0.0 | n.d. | 0.2 | 0.0 | 0.1 | 0.1 | 0.1 | 0.1 | 0.1 | n.d. |

| | | | | | | | | | | | | | | | | |
|---|---|---|---|---|---|---|---|---|---|---|---|---|---|---|---|---|
| LPC 20:4 | 0.1 | 0.1 | 0.1 | 0.0 | 0.2 | 0.1 | 0.1 | 0.0 | 0.2 | 0.3 | 0.1 | 0.0 | 0.5 | 0.7 | 0.2 | 0.2 |
| LPC 20:5 | n.d. | n.d. | n.d. | n.d. | n.d. | n.d. | n.d. | n.d. | n.d. | n.d. | 0.0 | n.d. | n.d. | n.d. | n.d. | n.d. |
| LPC 22:0 | n.d. | n.d. | n.d. | n.d. | n.d. | n.d. | n.d. | n.d. | n.d. | n.d. | n.d. | n.d. | n.d. | n.d. | 0.0 | 0.0 |
| LPC 22:4 | 0.0 | n.d. | n.d. | n.d. | 0.0 | n.d. | 0.0 | n.d. | 0.1 | n.d. | n.d. | n.d. | 0.0 | n.d. | 0.0 | n.d. |
| LPC 22:5 | 0.1 | n.d. | 0.0 | n.d. | 0.0 | n.d. | 0.0 | n.d. | 0.1 | 0.1 | n.d. | n.d. | 0.3 | n.d. | n.d. | n.d. |
| LPC 22:6 | 0.1 | 0.0 | 0.1 | n.d. | n.d. | n.d. | n.d. | n.d. | 0.0 | n.d. | n.d. | n.d. | 0.0 | 0.0 | 0.0 | 0.0 |
| LPC total | 24.1 | 13.6 | 24.3 | 10.7 | 25.1 | 20.1 | 19.5 | 4.0 | 37.2 | 26.8 | 26.9 | 2.4 | 36.1 | 51.0 | 30.4 | 17.6 |
| LPE 16:0 | n.d. | n.d. | n.d. | n.d. | 0.2 | 0.1 | n.d. | n.d. | 0.5 | n.d. | 0.4 | n.d. | 0.3 | 0.1 | n.d. | n.d. |
| LPE 18:0 | 0.2 | 0.0 | 0.8 | 1.0 | 0.4 | 0.1 | n.d. | n.d. | 0.2 | 0.1 | 0.8 | n.d. | 1.7 | n.d. | 0.3 | n.d. |
| LPE 18:1 | n.d. | n.d. | 0.1 | n.d. | 0.4 | n.d. | 0.2 | n.d. | n.d. | n.d. | n.d. | n.d. | 0.5 | 0.2 | n.d. | n.d. |
| LPE 18:2 | 0.2 | 0.2 | n.d. | n.d. | 0.4 | 0.2 | n.d. | n.d. | n.d. | n.d. | 0.3 | 0.3 | 0.2 | n.d. | 0.2 | n.d. |
| LPE total | 0.2 | 0.2 | 0.5 | 0.8 | 0.5 | 0.7 | 0.0 | 0.1 | 0.2 | 0.2 | 0.4 | 0.4 | 0.7 | 1.2 | 0.1 | 0.1 |
| PC 16:0_16:0 | 3.0 | 3.1 | 2.6 | 2.6 | 3.0 | 1.3 | 1.1 | 0.5 | 5.5 | 6.8 | 2.9 | 1.7 | 6.4 | 9.8 | 2.8 | 1.0 |
| PC 16:0_16:1 | 0.3 | 0.1 | n.d. | n.d. | n.d. | n.d. | 0.1 | n.d. | 0.9 | n.d. | n.d. | n.d. | 3.0 | n.d. | 0.6 | 0.5 |
| PC 16:0_18:0 | 8.8 | 6.2 | 6.1 | 6.4 | 6.3 | 5.6 | 4.2 | 2.6 | 14.6 | 13.2 | 9.1 | 3.5 | 15.4 | 23.6 | 15.9 | 7.8 |
| PC 16:0_18:1 | 47.6 | 33.3 | 40.8 | 22.6 | 45.1 | 40.8 | 36.3 | 2.6 | 74.0 | 55.6 | 48.1 | 12.8 | 72.0 | 101.9 | 81.3 | 48.7 |
| PC 16:0_18:2 | 80.5 | 46.6 | 74.4 | 34.2 | 75.8 | 61.0 | 65.6 | 21.4 | 120.0 | 85.2 | 88.3 | 12.7 | 113.8 | 150.2 | 124.1 | 64.2 |
| PC 16:0_18:3 | 1.8 | 1.7 | 2.7 | 1.4 | 1.0 | 1.3 | 1.1 | 1.0 | 2.1 | 1.7 | 1.3 | 0.8 | 8.6 | n.d. | 2.5 | 1.8 |
| PC 16:0_20:2 | 0.3 | 0.0 | 0.4 | n.d. | 0.7 | 0.3 | 0.1 | n.d. | 0.2 | 0.1 | 1.9 | n.d. | 5.9 | n.d. | 0.8 | 0.8 |
| PC 16:0_20:3 | 1.3 | 0.7 | 1.3 | 1.3 | 2.2 | 2.2 | 1.3 | 0.9 | 4.0 | 2.8 | 4.4 | 1.1 | 7.2 | 15.2 | 4.1 | 3.8 |
| PC 16:0_20:4 | 2.9 | 1.9 | 2.2 | 1.3 | 2.4 | 2.5 | 2.2 | 1.1 | 3.6 | 5.3 | 1.1 | 0.9 | 9.0 | 17.0 | 4.6 | 3.1 |
| PC 16:0_22:4 | 3.9 | n.d. | n.d. | n.d. | 0.6 | 0.2 | 0.7 | n.d. | 0.1 | n.d. | n.d. | n.d. | 10.4 | n.d. | 0.5 | 0.5 |
| PC 16:0_22:5 | n.d. | n.d. | n.d. | n.d. | 0.1 | n.d. | n.d. | n.d. | 1.0 | n.d. | 0.5 | n.d. | 3.3 | n.d. | 0.4 | 0.4 |
| PC 16:1_18:1 | 0.7 | 0.7 | 0.1 | n.d. | 0.5 | n.d. | n.d. | n.d. | 0.7 | n.d. | n.d. | n.d. | 1.7 | n.d. | 1.1 | 1.3 |
| PC 16:1_18:2 | 0.7 | n.d. | n.d. | n.d. | 0.6 | 0.1 | 0.4 | n.d. | 1.0 | n.d. | 0.5 | 0.1 | 1.3 | 1.7 | 1.1 | n.d. |

| | | | | | | | | | | | | | | | | |
|---|---|---|---|---|---|---|---|---|---|---|---|---|---|---|---|---|
| PC 18:0_18:0 | 4.0 | 3.2 | 2.1 | 1.4 | 3.5 | 3.3 | 0.9 | 1.0 | 6.2 | 5.8 | 3.2 | 1.2 | 5.9 | 10.0 | 4.5 | 3.3 |
| PC 18:0_18:1 | 38.9 | 31.9 | 28.1 | 12.7 | 34.3 | 30.8 | 34.8 | 12.8 | 54.8 | 36.5 | 46.8 | 12.2 | 46.6 | 59.4 | 64.0 | 32.3 |
| PC 18:0_18:2 | 104.9 | 67.9 | 95.5 | 30.7 | 104.1 | 85.6 | 102.1 | 24.0 | 143.9 | 91.8 | 124.5 | 19.8 | 125.5 | 137.2 | 168.2 | 92.8 |
| PC 18:0_18:3 | 0.9 | 0.7 | 0.7 | n.d. | 2.7 | 2.2 | 1.3 | n.d. | 2.9 | 0.6 | 0.7 | 1.0 | 2.3 | 1.9 | 4.6 | 5.9 |
| PC 18:0_20:2 | 0.5 | 0.3 | 0.3 | 0.0 | 2.9 | n.d. | 0.4 | 0.0 | 0.4 | 0.4 | 1.1 | 1.1 | 2.4 | 3.2 | 1.0 | 0.9 |
| PC 18:0_20:3 | 3.0 | 2.2 | 2.4 | 2.1 | 4.7 | 5.9 | 2.1 | 2.2 | 4.0 | 4.1 | 4.0 | 1.6 | 10.3 | 18.3 | 7.9 | 2.9 |
| PC 18:0_20:4 | 6.4 | 7.1 | 2.0 | 2.6 | 4.6 | 5.1 | 6.1 | 7.0 | 5.1 | 4.6 | 5.8 | 1.7 | 12.4 | 22.8 | 12.0 | 7.3 |
| PC 18:0_22:4 | n.d. | n.d. | 0.4 | n.d. | 0.2 | 0.1 | 0.4 | n.d. | 0.3 | 0.1 | 0.6 | 0.7 | 0.9 | n.d. | 1.7 | n.d. |
| PC 18:0_22:5 | n.d. | n.d. | n.d. | n.d. | n.d. | n.d. | n.d. | n.d. | n.d. | n.d. | n.d. | n.d. | 2.8 | 3.6 | n.d. | n.d. |
| PC 18:1_18:1 | 3.9 | 3.5 | 1.8 | 1.8 | 4.7 | 4.8 | 2.6 | 1.2 | 5.4 | 4.0 | 4.6 | 2.6 | 4.4 | 7.5 | 7.3 | 6.5 |
| PC 18:1_18:2 | 16.5 | 11.2 | 12.8 | 8.3 | 17.7 | 17.8 | 11.4 | 1.9 | 22.3 | 20.4 | 16.6 | 5.3 | 19.3 | 23.7 | 22.8 | 13.0 |
| PC 18:1_18:3 | 0.1 | n.d. | n.d. | n.d. | 0.2 | n.d. | n.d. | n.d. | n.d. | n.d. | 0.2 | 0.0 | 0.1 | n.d. | 0.4 | 0.1 |
| PC 18:1_20:2 | n.d. | n.d. | n.d. | n.d. | 0.6 | n.d. | n.d. | n.d. | 0.4 | n.d. | n.d. | n.d. | 0.7 | n.d. | n.d. | n.d. |
| PC 18:1_20:4 | n.d. | n.d. | n.d. | n.d. | 1.4 | 1.8 | 0.7 | 0.3 | n.d. | n.d. | n.d. | n.d. | 1.4 | n.d. | 0.3 | n.d. |
| PC 18:2_18:2 | 10.2 | 4.3 | 6.4 | 1.7 | 11.6 | 6.8 | 8.2 | 0.7 | 14.9 | 11.3 | 6.9 | 1.2 | 8.1 | 9.4 | 11.0 | 8.8 |
| PC 18:2_18:3 | 0.1 | n.d. | 0.4 | 0.1 | 1.3 | 0.4 | 0.1 | 0.0 | 0.3 | 0.2 | n.d. | n.d. | 0.8 | n.d. | 1.2 | n.d. |
| PC 18:2_20:1 | n.d. | n.d. | n.d. | n.d. | 0.4 | n.d. | 0.5 | n.d. | n.d. | n.d. | n.d. | n.d. | n.d. | n.d. | 0.9 | n.d. |
| PC 18:2_20:2 | 0.2 | n.d. | 0.1 | n.d. | 0.6 | n.d. | n.d. | n.d. | 1.3 | n.d. | n.d. | n.d. | 0.4 | n.d. | n.d. | n.d. |
| PC 18:2_20:3 | 0.8 | 0.3 | n.d. | n.d. | 0.3 | 0.2 | 0.1 | n.d. | 3.0 | n.d. | n.d. | n.d. | 2.7 | 3.6 | 0.5 | 0.4 |
| PC 18:2_20:4 | 0.4 | 0.1 | n.d. | n.d. | 0.4 | n.d. | 0.5 | n.d. | 0.2 | n.d. | 0.2 | 0.0 | 0.4 | n.d. | 1.5 | 2.0 |
| PC total | 330.2 | 224.0 | 280.3 | 121.9 | 316.2 | 276.9 | 281.3 | 57.3 | 479.3 | 341.7 | 370.3 | 56.5 | 461.3 | 621.3 | 539.5 | 300.2 |
| PC O-16:1/16:0 (PC P-16:0/16:0) | n.d. | n.d. | n.d. | n.d. | n.d. | n.d. | n.d. | n.d. | n.d. | n.d. | n.d. | n.d. | n.d. | n.d. | n.d. | n.d. |
| PC O-16:1/18:1 (PC P-16:0/18:1) | 2.0 | n.d. | 2.4 | n.d. | n.d. | n.d. | n.d. | n.d. | n.d. | n.d. | n.d. | n.d. | 2.0 | n.d. | 7.1 | n.d. |
| PC O-16:1/18:2 (PC P-16:0/18:2) | n.d. | n.d. | n.d. | n.d. | 1.2 | n.d. | n.d. | n.d. | 6.9 | 0.3 | 0.4 | n.d. | n.d. | n.d. | 0.7 | n.d. |
| PC O-16:1/20:4 (PC P-16:0/20:4) | 1.6 | n.d. | n.d. | n.d. | 0.9 | n.d. | n.d. | n.d. | n.d. | n.d. | 2.9 | n.d. | 0.3 | n.d. | 3.8 | n.d. |

| | | | | | | | | | | | | | | | | |
|---|---|---|---|---|---|---|---|---|---|---|---|---|---|---|---|---|
| PC O-18:1/18:2 (PC P-18:0/18:2) | n.d. | n.d. | n.d. | n.d. | n.d. | n.d. | n.d. | n.d. | n.d. | n.d. | n.d. | n.d. | 1.0 | n.d. | n.d. | n.d. |
| PC O-18:1/20:4 (PC P-18:0/20:4) | n.d. | n.d. | n.d. | n.d. | n.d. | n.d. | n.d. | n.d. | n.d. | n.d. | n.d. | n.d. | n.d. | n.d. | n.d. | n.d. |
| PC P-18:2/20:5 | 0.3 | n.d. | n.d. | n.d. | 2.8 | 2.4 | 2.9 | 0.2 | 4.4 | 4.6 | 2.2 | n.d. | 7.1 | 5.5 | 4.5 | 3.8 |
| EtherPC total | 1.6 | 2.5 | 0.6 | 1.2 | 1.7 | 2.6 | 1.5 | 1.7 | 5.4 | 6.9 | 1.4 | 1.7 | 4.9 | 6.3 | 6.3 | 5.2 |
| PE 14:1_18:2 | n.d. | n.d. | n.d. | n.d. | 0.1 | n.d. | n.d. | n.d. | n.d. | n.d. | n.d. | n.d. | n.d. | n.d. | n.d. | n.d. |
| PE 16:0_18:1 | 0.5 | 0.5 | n.d. | n.d. | 0.2 | n.d. | n.d. | n.d. | n.d. | n.d. | n.d. | n.d. | n.d. | n.d. | 0.2 | n.d. |
| PE 16:0_18:2 | 0.8 | 0.4 | n.d. | n.d. | 0.8 | 0.9 | n.d. | n.d. | 0.3 | 0.2 | 0.4 | 0.2 | 0.8 | 0.8 | 1.6 | n.d. |
| PE 16:0_20:4 | n.d. | n.d. | n.d. | n.d. | n.d. | n.d. | n.d. | n.d. | 0.3 | n.d. | n.d. | n.d. | 0.2 | n.d. | 0.6 | n.d. |
| PE 16:1_18:2 | 0.5 | n.d. | n.d. | n.d. | 0.1 | 0.0 | 0.2 | 0.2 | n.d. | n.d. | 0.3 | n.d. | 0.2 | n.d. | 0.1 | 0.0 |
| PE 16:1_20:4 | n.d. | n.d. | n.d. | n.d. | n.d. | n.d. | 0.1 | n.d. | n.d. | n.d. | n.d. | n.d. | n.d. | n.d. | n.d. | n.d. |
| PE 18:0_18:1 | 0.7 | 0.5 | 0.3 | 0.3 | 0.3 | 0.2 | 0.3 | 0.1 | 0.8 | 0.8 | 0.9 | n.d. | 0.2 | n.d. | 0.2 | 0.1 |
| PE 18:0_18:2 | 4.4 | 3.7 | 2.3 | 1.1 | 4.3 | 3.9 | 3.4 | 2.1 | 7.1 | 6.1 | 3.7 | 1.1 | 4.7 | 3.8 | 3.8 | 2.4 |
| PE 18:0_18:3 | n.d. | n.d. | n.d. | n.d. | 0.3 | 0.3 | n.d. | n.d. | n.d. | n.d. | n.d. | n.d. | n.d. | n.d. | n.d. | n.d. |
| PE 18:0_20:3 | n.d. | n.d. | n.d. | n.d. | n.d. | n.d. | 0.1 | n.d. | n.d. | n.d. | n.d. | n.d. | n.d. | n.d. | 0.1 | 0.1 |
| PE 18:0_20:4 | 1.3 | 0.8 | 0.7 | 0.7 | 1.4 | 1.9 | 0.4 | 0.4 | 0.8 | 1.2 | 0.7 | 0.5 | 1.5 | 2.5 | 1.8 | 1.6 |
| PE 18:0_22:4 | n.d. | n.d. | n.d. | n.d. | n.d. | n.d. | 0.2 | n.d. | n.d. | n.d. | 0.1 | n.d. | n.d. | n.d. | 0.1 | n.d. |
| PE 18:1_18:1 | 0.3 | 0.3 | n.d. | n.d. | 0.0 | n.d. | 0.1 | n.d. | n.d. | n.d. | 0.2 | 0.0 | 0.2 | n.d. | 0.1 | n.d. |
| PE 18:1_18:2 | 0.1 | n.d. | n.d. | n.d. | 0.1 | 0.0 | n.d. | n.d. | 0.6 | n.d. | n.d. | n.d. | 0.6 | 0.5 | 0.2 | 0.2 |
| PE 18:1_20:4 | n.d. | n.d. | n.d. | n.d. | n.d. | n.d. | n.d. | n.d. | n.d. | n.d. | n.d. | n.d. | n.d. | n.d. | n.d. | n.d. |
| PE 18:2_18:2 | 0.3 | n.d. | n.d. | n.d. | n.d. | n.d. | n.d. | n.d. | n.d. | n.d. | n.d. | n.d. | 0.1 | n.d. | n.d. | n.d. |
| PE 18:2_20:0 | n.d. | n.d. | n.d. | n.d. | n.d. | n.d. | n.d. | n.d. | n.d. | n.d. | n.d. | n.d. | n.d. | n.d. | n.d. | n.d. |
| PE 18:2_20:1 | n.d. | n.d. | n.d. | n.d. | n.d. | n.d. | 0.0 | n.d. | n.d. | n.d. | n.d. | n.d. | n.d. | n.d. | n.d. | n.d. |
| PE total | 6.6 | 5.2 | 2.7 | 1.8 | 6.0 | 6.4 | 4.2 | 2.7 | 8.6 | 7.8 | 4.7 | 1.4 | 7.3 | 7.7 | 6.2 | 5.2 |
| PE P-16:0/16:0 | n.d. | n.d. | n.d. | n.d. | n.d. | n.d. | n.d. | n.d. | n.d. | n.d. | n.d. | n.d. | n.d. | n.d. | n.d. | n.d. |
| PE P-16:0/18:1 | 0.4 | 0.1 | 0.3 | n.d. | n.d. | n.d. | 1.0 | 0.9 | n.d. | n.d. | 1.1 | n.d. | 4.5 | n.d. | 1.0 | 0.4 |

| | | | | | | | | | | | | | | | | |
|---|---|---|---|---|---|---|---|---|---|---|---|---|---|---|---|---|
| PE P-16:0/18:2 | 5.1 | 3.9 | 5.9 | 1.5 | 7.8 | 6.0 | 4.0 | 3.8 | 8.1 | 6.7 | 6.4 | 2.4 | 3.5 | 5.4 | 7.3 | 3.1 |
| PE P-16:0/18:3 | n.d. | n.d. | n.d. | n.d. | n.d. | n.d. | n.d. | n.d. | n.d. | n.d. | n.d. | n.d. | n.d. | n.d. | n.d. | n.d. |
| PE P-16:0/20:3 | n.d. | n.d. | n.d. | n.d. | 1.1 | n.d. | n.d. | n.d. | n.d. | n.d. | n.d. | n.d. | n.d. | n.d. | 1.6 | n.d. |
| PE P-16:0/20:4 | 1.7 | 1.5 | 1.3 | 1.2 | 2.4 | 2.0 | 1.2 | 0.4 | 6.0 | n.d. | 0.8 | 0.7 | 3.6 | 4.3 | 2.6 | 2.6 |
| PE P-16:0/22:4 | n.d. | n.d. | n.d. | n.d. | n.d. | n.d. | n.d. | n.d. | 0.6 | n.d. | 0.4 | n.d. | 1.9 | n.d. | n.d. | n.d. |
| PE P-16:0/22:5 | n.d. | n.d. | n.d. | n.d. | n.d. | n.d. | n.d. | n.d. | n.d. | n.d. | 0.6 | n.d. | 2.3 | 2.0 | 0.6 | 0.2 |
| PE P-16:0/22:6 | n.d. | n.d. | n.d. | n.d. | 0.9 | n.d. | n.d. | n.d. | n.d. | n.d. | n.d. | n.d. | 3.0 | n.d. | n.d. | n.d. |
| PE P-16:1/22:2 | n.d. | n.d. | n.d. | n.d. | n.d. | n.d. | n.d. | n.d. | n.d. | n.d. | n.d. | n.d. | n.d. | n.d. | n.d. | n.d. |
| PE P-18:0/16:0 | n.d. | n.d. | n.d. | n.d. | n.d. | n.d. | 2.3 | n.d. | n.d. | n.d. | n.d. | n.d. | n.d. | n.d. | n.d. | n.d. |
| PE P-18:0/18:0 | n.d. | n.d. | n.d. | n.d. | n.d. | n.d. | n.d. | n.d. | n.d. | n.d. | n.d. | n.d. | n.d. | n.d. | n.d. | n.d. |
| PE P-18:0/18:1 | 1.4 | 0.9 | 1.1 | 1.2 | 5.3 | 3.5 | 0.7 | n.d. | 3.8 | 2.1 | 1.4 | n.d. | 2.3 | 2.4 | 1.2 | 1.4 |
| PE P-18:0/18:2 | 27.5 | 22.9 | 19.6 | 1.8 | 23.0 | 21.4 | 18.0 | 7.0 | 26.6 | 12.9 | 23.7 | 13.0 | 19.0 | 20.9 | 28.5 | 12.2 |
| PE P-18:0/18:3 | n.d. | n.d. | n.d. | n.d. | 0.4 | n.d. | n.d. | n.d. | n.d. | n.d. | 0.7 | n.d. | n.d. | n.d. | 2.0 | 0.4 |
| PE P-18:0/20:3 | 0.8 | n.d. | n.d. | n.d. | n.d. | n.d. | n.d. | n.d. | n.d. | n.d. | n.d. | n.d. | 0.5 | n.d. | n.d. | n.d. |
| PE P-18:0/20:4 | 6.1 | 7.8 | 1.1 | 0.6 | 1.9 | 0.5 | 0.9 | 1.0 | 5.3 | 5.8 | 1.4 | 1.1 | 5.3 | 8.2 | 5.7 | 3.0 |
| PE P-18:0/22:4 | 2.7 | n.d. | n.d. | n.d. | n.d. | n.d. | 0.7 | n.d. | 0.3 | n.d. | 0.7 | n.d. | 1.3 | n.d. | n.d. | n.d. |
| PE P-18:0/22:5 | 1.3 | n.d. | 1.1 | n.d. | 0.4 | 0.1 | n.d. | n.d. | 0.3 | n.d. | n.d. | n.d. | 4.0 | n.d. | 1.4 | n.d. |
| PE P-18:0/22:6 | n.d. | n.d. | n.d. | n.d. | n.d. | n.d. | n.d. | n.d. | 2.4 | n.d. | n.d. | n.d. | 1.2 | n.d. | 0.8 | 0.2 |
| PE P-18:1/16:0 | n.d. | n.d. | 0.6 | n.d. | 1.5 | n.d. | n.d. | n.d. | n.d. | n.d. | n.d. | n.d. | n.d. | n.d. | 0.2 | n.d. |
| PE P-18:1/18:1 | 0.8 | 0.7 | 0.9 | 0.9 | 0.9 | 1.1 | n.d. | n.d. | 1.2 | 1.2 | 2.1 | n.d. | 1.6 | 1.7 | 1.7 | 0.5 |
| PE P-18:1/18:2 | 5.2 | 3.6 | 3.7 | 3.0 | 3.8 | 3.3 | 2.7 | 3.0 | 5.6 | 4.0 | 8.6 | 3.1 | 5.0 | 4.6 | 8.2 | 4.8 |
| PE P-18:1/18:3 | n.d. | n.d. | n.d. | n.d. | n.d. | n.d. | n.d. | n.d. | n.d. | n.d. | n.d. | n.d. | n.d. | n.d. | n.d. | n.d. |
| PE P-18:1/20:3 | n.d. | n.d. | n.d. | n.d. | n.d. | n.d. | n.d. | n.d. | n.d. | n.d. | n.d. | n.d. | n.d. | n.d. | n.d. | n.d. |
| PE P-18:1/20:4 | 1.8 | 1.1 | n.d. | n.d. | 2.2 | 0.8 | n.d. | n.d. | 1.9 | n.d. | 0.9 | n.d. | 1.7 | 1.7 | 2.3 | 2.2 |
| PE P-18:1/22:4 | n.d. | n.d. | n.d. | n.d. | n.d. | n.d. | n.d. | n.d. | n.d. | n.d. | n.d. | n.d. | n.d. | n.d. | n.d. | n.d. |

| | | | | | | | | | | | | | | | | |
|---|---|---|---|---|---|---|---|---|---|---|---|---|---|---|---|---|
| PE P-18:1/22:5 | 0.2 | n.d. | n.d. | n.d. | n.d. | n.d. | n.d. | n.d. | 0.3 | n.d. | n.d. | n.d. | n.d. | n.d. | n.d. | n.d. |
| PE P-18:1/22:6 | n.d. | n.d. | n.d. | n.d. | n.d. | n.d. | n.d. | n.d. | n.d. | n.d. | n.d. | n.d. | 1.4 | n.d. | n.d. | n.d. |
| PE P-18:2/18:1 | n.d. | n.d. | n.d. | n.d. | n.d. | n.d. | n.d. | n.d. | n.d. | n.d. | n.d. | n.d. | n.d. | n.d. | n.d. | n.d. |
| PE P-18:2/18:2 | n.d. | n.d. | n.d. | n.d. | n.d. | n.d. | n.d. | n.d. | n.d. | n.d. | 0.8 | n.d. | n.d. | n.d. | n.d. | n.d. |
| PE P-18:2/20:4 | n.d. | n.d. | n.d. | n.d. | n.d. | n.d. | n.d. | n.d. | n.d. | n.d. | n.d. | n.d. | 0.5 | n.d. | n.d. | n.d. |
| EtherPE total | 45.0 | 40.0 | 31.3 | 8.7 | 37.4 | 36.0 | 27.4 | 14.0 | 49.5 | 29.7 | 42.8 | 12.0 | 41.6 | 55.3 | 55.4 | 23.3 |
| PI 16:0_16:0 | n.d. | n.d. | n.d. | n.d. | n.d. | n.d. | n.d. | n.d. | 7.8 | 5.3 | n.d. | n.d. | n.d. | n.d. | n.d. | n.d. |
| PI 16:0_18:0 | n.d. | n.d. | n.d. | n.d. | n.d. | n.d. | n.d. | n.d. | n.d. | n.d. | n.d. | n.d. | n.d. | n.d. | n.d. | n.d. |
| PI 16:0_18:1 | 20.4 | 19.3 | 4.5 | n.d. | 6.9 | n.d. | 11.2 | 3.0 | 13.8 | 2.2 | 12.8 | n.d. | 126.7 | n.d. | 21.0 | 15.4 |
| PI 16:0_18:2 | 15.5 | 4.1 | 19.2 | 6.5 | 30.6 | 14.5 | n.d. | n.d. | 68.2 | 69.2 | 28.0 | n.d. | 104.7 | n.d. | 16.3 | 9.0 |
| PI 16:0_20:4 | n.d. | n.d. | n.d. | n.d. | 5.2 | 5.0 | n.d. | n.d. | 20.9 | n.d. | n.d. | n.d. | 32.6 | n.d. | n.d. | n.d. |
| PI 16:1_18:0 | n.d. | n.d. | 14.8 | n.d. | 9.6 | n.d. | n.d. | n.d. | 11.9 | n.d. | n.d. | n.d. | 97.6 | n.d. | n.d. | n.d. |
| PI 18:0_18:0 | 6.7 | n.d. | n.d. | n.d. | n.d. | n.d. | n.d. | n.d. | n.d. | n.d. | n.d. | n.d. | n.d. | n.d. | n.d. | n.d. |
| PI 18:0_18:1 | 30.9 | 23.5 | 42.4 | 23.4 | 99.0 | 143.1 | 14.0 | 6.7 | 102.8 | 159.5 | 63.8 | 19.2 | 212.1 | 350.7 | 119.4 | 85.5 |
| PI 18:0_18:2 | 176.2 | 119.1 | 299.6 | 256.3 | 351.8 | 198.9 | 242.4 | 134.4 | 528.2 | 475.7 | 261.5 | 149.7 | 518.2 | 1017.3 | 328.8 | 148.7 |
| PI 18:0_18:3 | n.d. | n.d. | n.d. | n.d. | n.d. | n.d. | n.d. | n.d. | n.d. | n.d. | n.d. | n.d. | n.d. | n.d. | n.d. | n.d. |
| PI 18:0_20:2 | 10.5 | 6.6 | 16.2 | n.d. | 10.1 | 6.8 | n.d. | n.d. | 5.7 | n.d. | 2.4 | n.d. | 54.2 | n.d. | 10.8 | 1.0 |
| PI 18:0_20:3 | 22.3 | 9.7 | 20.8 | 20.2 | 32.6 | n.d. | 10.6 | 2.4 | 61.9 | n.d. | 13.4 | n.d. | 91.5 | n.d. | 45.1 | 43.0 |
| PI 18:0_20:4 | 67.8 | 57.8 | 28.8 | 10.5 | 57.8 | 67.4 | 74.3 | 46.9 | 107.1 | 95.9 | 55.5 | 15.4 | 347.4 | 464.2 | 93.0 | 69.0 |
| PI 18:1_18:1 | n.d. | n.d. | 5.2 | n.d. | n.d. | n.d. | n.d. | n.d. | n.d. | n.d. | n.d. | n.d. | 2.5 | n.d. | n.d. | n.d. |
| PI 18:1_18:2 | 2.2 | n.d. | n.d. | n.d. | 14.1 | n.d. | n.d. | n.d. | n.d. | n.d. | n.d. | n.d. | n.d. | n.d. | 3.4 | n.d. |
| PI total | 305.4 | 194.8 | 405.7 | 272.5 | 359.7 | 437.2 | 341.6 | 181.7 | 653.2 | 760.2 | 379.0 | 180.3 | 886.4 | 1818.6 | 592.7 | 329.1 |
| SM 18:1;O2/14:0 | 0.1 | 0.1 | 0.2 | 0.1 | 0.1 | 0.0 | 0.1 | 0.1 | 0.2 | 0.1 | 0.1 | 0.0 | 0.1 | 0.1 | 0.1 | 0.1 |
| SM 18:1;O2/14:1 | 0.0 | n.d. | n.d. | n.d. | 0.0 | n.d. | n.d. | n.d. | 0.0 | 0.0 | n.d. | n.d. | n.d. | n.d. | n.d. | n.d. |
| SM 18:1;O2/16:0 | 16.0 | 7.3 | 19.8 | 4.9 | 13.0 | 6.0 | 13.6 | 2.1 | 18.3 | 11.7 | 15.1 | 1.9 | 15.8 | 16.9 | 20.0 | 7.6 |

| | | | | | | | | | | | | | | | | |
|---|---|---|---|---|---|---|---|---|---|---|---|---|---|---|---|---|
| SM 18:1;O2/16:1 | 0.9 | 0.4 | 1.1 | 0.4 | 0.7 | 0.4 | 0.8 | 0.1 | 1.0 | 0.6 | 0.9 | 0.2 | 0.9 | 0.8 | 1.2 | 0.4 |
| SM 18:1;O2/18:0 | 3.0 | 1.1 | 3.9 | 0.8 | 2.5 | 1.3 | 2.5 | 0.4 | 3.2 | 1.6 | 3.0 | 0.3 | 3.0 | 2.9 | 3.5 | 1.1 |
| SM 18:1;O2/18:1 | 0.6 | 0.3 | 0.9 | 0.3 | 0.7 | 0.3 | 0.6 | 0.1 | 0.8 | 0.5 | 0.8 | 0.1 | 0.6 | 0.6 | 0.9 | 0.2 |
| SM 18:1;O2/18:2 | n.d. | n.d. | n.d. | n.d. | n.d. | n.d. | n.d. | n.d. | n.d. | n.d. | n.d. | n.d. | 0.0 | n.d. | 0.0 | 0.0 |
| SM 18:1;O2/20:0 | 1.0 | 0.5 | 1.1 | 0.2 | 0.8 | 0.5 | 0.8 | 0.1 | 1.1 | 0.6 | 0.9 | 0.1 | 0.9 | 0.8 | 1.1 | 0.4 |
| SM 18:1;O2/20:1 | 0.5 | 0.2 | 0.6 | 0.1 | 0.4 | 0.3 | 0.4 | 0.1 | 0.4 | 0.3 | 0.5 | 0.2 | 0.4 | 0.4 | 0.6 | 0.2 |
| SM 18:1;O2/20:2 | 0.0 | n.d. | 0.0 | 0.0 | 0.0 | n.d. | 0.0 | n.d. | 0.0 | 0.0 | 0.0 | n.d. | 0.0 | n.d. | 0.0 | n.d. |
| SM 18:1;O2/22:0 | 1.2 | 0.6 | 1.6 | 0.4 | 1.0 | 0.5 | 1.2 | 0.2 | 1.3 | 0.8 | 1.3 | 0.3 | 1.3 | 1.3 | 1.6 | 0.5 |
| SM 18:1;O2/22:1 | 0.6 | 0.3 | 0.9 | 0.1 | 0.6 | 0.4 | 0.7 | 0.1 | 0.8 | 0.4 | 0.8 | 0.1 | 0.7 | 0.7 | 0.9 | 0.4 |
| SM 18:1;O2/22:2 | 0.1 | 0.1 | 0.1 | 0.1 | 0.1 | 0.0 | 0.1 | 0.0 | 0.1 | 0.1 | 0.1 | 0.1 | 0.1 | 0.1 | 0.1 | 0.0 |
| SM 18:1;O2/22:3 | 0.0 | 0.0 | 0.0 | 0.0 | 0.0 | 0.0 | 0.0 | n.d. | 0.0 | n.d. | 0.0 | n.d. | 0.0 | n.d. | 0.0 | 0.0 |
| SM 18:1;O2/22:5 | n.d. | n.d. | n.d. | n.d. | n.d. | n.d. | n.d. | n.d. | n.d. | n.d. | n.d. | n.d. | n.d. | n.d. | n.d. | n.d. |
| SM total | 24.1 | 10.6 | 30.3 | 7.0 | 19.9 | 9.7 | 20.6 | 3.0 | 27.3 | 16.4 | 23.6 | 3.0 | 23.8 | 24.5 | 30.0 | 10.9 |
| Cer 18:1;O2/16:0 | 0.2 | 0.1 | 0.2 | 0.0 | 0.2 | 0.1 | 0.2 | 0.1 | 0.2 | 0.1 | 0.2 | 0.0 | 0.2 | 0.3 | 0.2 | 0.1 |
| Cer 18:1;O2/18:0 | 0.3 | 0.1 | 0.1 | 0.1 | 0.2 | 0.1 | 0.2 | 0.1 | 0.3 | 0.1 | 0.2 | 0.1 | 0.3 | 0.4 | 0.3 | 0.2 |
| Cer 18:1;O2/18:1 | n.d. | n.d. | n.d. | n.d. | n.d. | n.d. | n.d. | n.d. | n.d. | n.d. | n.d. | n.d. | n.d. | n.d. | n.d. | n.d. |
| Cer 18:1;O2/18:2 | 0.0 | 0.0 | 0.0 | 0.0 | 0.0 | 0.0 | 0.0 | 0.0 | 0.0 | 0.0 | 0.0 | 0.0 | 0.0 | 0.0 | 0.0 | 0.0 |
| Cer 18:1;O2/20:0 | 0.2 | 0.1 | 0.1 | 0.1 | 0.2 | 0.2 | 0.2 | 0.0 | 0.3 | 0.2 | 0.2 | 0.0 | 0.3 | 0.4 | 0.2 | 0.1 |
| Cer 18:1;O2/20:1 | n.d. | n.d. | n.d. | n.d. | n.d. | n.d. | n.d. | n.d. | n.d. | n.d. | n.d. | n.d. | n.d. | n.d. | n.d. | n.d. |
| Cer 18:1;O2/20:2 | 0.0 | 0.0 | 0.0 | n.d. | 0.0 | 0.0 | 0.0 | 0.0 | 0.0 | 0.0 | 0.0 | n.d. | 0.0 | 0.0 | 0.0 | 0.0 |
| Cer 18:1;O2/20:4 | n.d. | n.d. | n.d. | n.d. | n.d. | n.d. | n.d. | n.d. | n.d. | n.d. | n.d. | n.d. | 0.0 | n.d. | n.d. | n.d. |
| Cer 18:1;O2/22:0 | 0.9 | 0.5 | 0.6 | 0.2 | 0.9 | 0.7 | 0.7 | 0.2 | 1.1 | 0.8 | 0.9 | 0.1 | 1.3 | 1.7 | 1.2 | 0.7 |
| Cer 18:1;O2/22:1 | 0.0 | 0.0 | 0.0 | 0.0 | 0.0 | 0.0 | 0.0 | n.d. | 0.0 | 0.0 | 0.0 | n.d. | 0.0 | 0.0 | 0.0 | 0.0 |
| Cer 18:1;O2/22:2 | 0.2 | 0.1 | 0.2 | 0.1 | 0.2 | 0.2 | 0.2 | 0.1 | 0.4 | 0.2 | 0.2 | 0.1 | 0.3 | 0.3 | 0.3 | 0.2 |
| Cer 18:1;O2/22:3 | n.d. | n.d. | n.d. | n.d. | n.d. | n.d. | n.d. | n.d. | n.d. | n.d. | n.d. | n.d. | n.d. | n.d. | n.d. | n.d. |

| | | | | | | | | | | | | | | | | |
|---|---|---|---|---|---|---|---|---|---|---|---|---|---|---|---|---|
| Cer 18:1;O2/22:4 | n.d. | n.d. | n.d. | n.d. | n.d. | n.d. | n.d. | n.d. | n.d. | n.d. | n.d. | n.d. | 0.0 | n.d. | n.d. | n.d. |
| Cer total | 1.9 | 0.9 | 1.2 | 0.4 | 1.7 | 1.2 | 1.5 | 0.5 | 2.3 | 1.5 | 1.7 | 0.3 | 2.4 | 3.2 | 2.3 | 1.3 |
| CE 14:0 | 2.2 | 1.4 | 2.7 | 0.9 | 1.9 | 1.2 | 2.0 | 0.4 | 2.9 | 1.9 | 2.1 | 0.3 | 2.8 | 3.7 | 3.3 | 1.8 |
| CE 14:1 | 0.4 | 0.4 | 0.7 | 0.3 | 0.5 | 0.4 | 0.7 | 0.2 | 0.6 | 0.5 | 0.8 | 0.1 | 0.8 | 1.2 | 0.9 | 0.5 |
| CE 16:0 | 218.1 | 133.9 | 255.0 | 84.2 | 184.9 | 112.4 | 172.1 | 28.5 | 257.2 | 160.3 | 197.6 | 20.1 | 285.1 | 392.7 | 281.7 | 137.9 |
| CE 16:1 | 108.6 | 87.7 | 129.2 | 43.5 | 102.1 | 82.2 | 113.8 | 12.1 | 161.0 | 123.7 | 154.2 | 14.9 | 215.9 | 347.3 | 231.1 | 128.5 |
| CE 18:0 | 82.2 | 54.7 | 87.4 | 23.0 | 69.4 | 46.0 | 64.7 | 7.5 | 106.9 | 76.5 | 77.4 | 7.4 | 128.4 | 184.8 | 129.6 | 81.5 |
| CE 18:1 | 994.4 | 679.6 | 965.5 | 295.3 | 909.2 | 613.4 | 793.8 | 92.3 | 1429.1 | 1024.9 | 1008.0 | 59.4 | 1382.4 | 1797.6 | 1552.4 | 922.0 |
| CE 18:2 | 3718.7 | 1998.4 | 4265.3 | 1544.9 | 3260.3 | 1819.1 | 3183.1 | 543.8 | 4839.1 | 2960.6 | 3878.8 | 790.3 | 3939.2 | 4602.1 | 4770.4 | 2205.5 |
| CE 18:3 | 328.3 | 217.3 | 398.6 | 162.8 | 279.9 | 194.6 | 275.1 | 41.8 | 410.4 | 282.7 | 326.1 | 64.3 | 392.9 | 549.9 | 481.0 | 317.1 |
| CE 18:4 | 1.2 | 0.8 | 1.4 | 0.7 | 0.9 | 0.7 | 0.9 | 0.1 | 1.3 | 0.9 | 1.0 | 0.2 | 6.2 | 13.1 | 2.7 | 1.9 |
| CE 20:0 | 1.5 | 0.9 | 1.8 | 0.6 | 1.3 | 0.6 | 1.3 | 0.2 | 1.7 | 0.9 | 1.2 | 0.1 | 1.3 | 1.4 | 1.5 | 0.8 |
| CE 20:1 | 2.9 | 2.5 | 2.8 | 1.2 | 2.5 | 1.9 | 2.1 | 0.6 | 4.4 | 3.7 | 2.4 | 0.5 | 3.7 | 4.9 | 4.0 | 2.2 |
| CE 20:2 | 3.2 | 2.6 | 3.2 | 1.7 | 2.9 | 2.4 | 2.7 | 0.8 | 5.1 | 4.2 | 3.2 | 0.9 | 5.6 | 8.5 | 5.6 | 3.4 |
| CE 20:3 | 13.2 | 8.5 | 17.1 | 7.0 | 11.0 | 7.3 | 11.1 | 1.8 | 18.2 | 14.4 | 12.8 | 2.0 | 24.2 | 40.5 | 20.7 | 11.8 |
| CE 20:4 | 138.2 | 82.9 | 188.4 | 69.7 | 124.3 | 83.8 | 124.5 | 14.1 | 187.5 | 139.4 | 155.9 | 32.9 | 243.5 | 401.1 | 205.8 | 93.1 |
| CE 20:5 | 6.5 | 4.2 | 9.2 | 3.3 | 5.0 | 3.7 | 5.0 | 1.0 | 7.3 | 5.4 | 5.2 | 1.5 | 12.2 | 22.3 | 7.9 | 4.5 |
| CE 22:0 | 0.5 | 0.4 | 0.7 | 0.3 | 0.4 | 0.3 | 0.3 | 0.1 | 0.6 | 0.4 | 0.4 | 0.1 | 0.4 | 0.4 | 0.4 | 0.2 |
| CE 22:1 | 0.4 | 0.3 | 0.5 | 0.2 | 0.4 | 0.2 | 0.3 | 0.0 | 0.5 | 0.5 | 0.4 | 0.2 | 0.4 | 0.4 | 0.5 | 0.3 |
| CE 22:2 | 0.1 | 0.1 | 0.1 | 0.1 | 0.1 | 0.1 | 0.1 | 0.0 | 0.2 | 0.2 | 0.1 | 0.0 | 0.2 | 0.2 | 0.1 | 0.1 |
| CE 22:3 | 0.1 | 0.1 | 0.1 | 0.1 | 0.1 | 0.0 | 0.1 | 0.0 | 0.1 | 0.1 | 0.0 | 0.0 | 0.1 | 0.3 | 0.2 | 0.1 |
| CE 22:4 | 0.5 | 0.3 | 0.8 | 0.3 | 0.5 | 0.4 | 0.4 | 0.1 | 0.7 | 0.6 | 0.6 | 0.2 | 1.2 | 2.1 | 1.0 | 0.6 |
| CE 22:5 | 1.4 | 0.9 | 2.5 | 1.2 | 1.3 | 1.0 | 1.2 | 0.3 | 2.0 | 1.4 | 1.7 | 0.4 | 3.9 | 7.0 | 3.1 | 1.7 |
| CE 22:6 | 1.8 | 1.2 | 2.5 | 1.4 | 1.4 | 1.0 | 1.2 | 0.3 | 2.2 | 1.3 | 1.8 | 0.6 | 3.7 | 6.4 | 2.9 | 1.1 |
| CE total | 5624.4 | 3264.2 | 6335.6 | 2229.7 | 4960.2 | 2958.1 | 4756.5 | 718.6 | 7438.9 | 4771.0 | 5831.5 | 931.1 | 6654.1 | 8382.1 | 7706.9 | 3889.8 |

| | | | | | | | | | | | | | | | | |
|---|---|---|---|---|---|---|---|---|---|---|---|---|---|---|---|---|
| MG 16:0 | 3.6 | 0.9 | 4.7 | 2.0 | 6.0 | 3.2 | 6.3 | 3.7 | 3.8 | 0.6 | 4.1 | 1.0 | 4.0 | 1.4 | 3.1 | 1.2 |
| MG 18:0 | 27.0 | 5.8 | 23.3 | 5.3 | 21.6 | 6.1 | 25.7 | 6.9 | 24.1 | 2.7 | 23.5 | 12.4 | 19.6 | 3.5 | 18.1 | 4.1 |
| MG 18:1 | 1.7 | 0.4 | 3.2 | 1.3 | 2.9 | 2.1 | 2.9 | 1.6 | 2.0 | 1.0 | 2.0 | 1.0 | 2.9 | 1.1 | 2.5 | 1.3 |
| MG 18:2 | 1.6 | 0.8 | 2.0 | 0.8 | 2.5 | 1.0 | 2.4 | 1.6 | 3.0 | 3.4 | 2.1 | 0.7 | 2.1 | 1.5 | 1.2 | 0.4 |
| MG total | 33.9 | 7.4 | 33.2 | 8.0 | 33.0 | 10.5 | 37.3 | 9.3 | 32.9 | 4.8 | 31.7 | 11.8 | 28.5 | 4.2 | 24.9 | 5.6 |
| DG 14:0_14:0 | 0.0 | 0.0 | 0.0 | 0.0 | 0.0 | 0.0 | 0.0 | n.d. | 0.0 | n.d. | 0.0 | n.d. | 0.0 | n.d. | 0.0 | 0.0 |
| DG 14:0_16:0 | 0.0 | 0.0 | 0.0 | 0.0 | 0.0 | 0.0 | 0.0 | 0.0 | 0.0 | 0.0 | 0.0 | 0.0 | 0.0 | 0.0 | 0.0 | 0.0 |
| DG 14:0_16:1 | 0.0 | 0.0 | 0.0 | 0.0 | 0.0 | 0.0 | 0.0 | 0.0 | 0.0 | 0.0 | 0.0 | 0.0 | 0.0 | 0.0 | 0.0 | 0.0 |
| DG 14:0_18:0 | 0.0 | n.d. | 0.0 | n.d. | 0.0 | n.d. | 0.0 | n.d. | 0.0 | 0.0 | n.d. | n.d. | 0.0 | n.d. | n.d. | n.d. |
| DG 14:0_18:1 | 0.0 | 0.0 | 0.0 | 0.0 | 0.0 | 0.0 | 0.0 | 0.0 | 0.0 | 0.0 | 0.0 | 0.0 | 0.0 | 0.0 | 0.0 | 0.0 |
| DG 14:0_18:2 | 0.0 | 0.0 | 0.0 | 0.0 | 0.0 | 0.0 | 0.0 | 0.0 | 0.0 | 0.0 | 0.0 | 0.0 | 0.0 | 0.0 | 0.0 | 0.0 |
| DG 14:0_18:3 | n.d. | n.d. | n.d. | n.d. | n.d. | n.d. | n.d. | n.d. | 0.0 | n.d. | n.d. | n.d. | n.d. | n.d. | n.d. | n.d. |
| DG 14:0_20:0 | n.d. | n.d. | n.d. | n.d. | n.d. | n.d. | n.d. | n.d. | n.d. | n.d. | n.d. | n.d. | n.d. | n.d. | n.d. | n.d. |
| DG 14:1_18:1 | 0.0 | n.d. | n.d. | n.d. | 0.0 | 0.0 | 0.0 | 0.0 | 0.0 | n.d. | 0.0 | n.d. | 0.0 | 0.0 | 0.0 | 0.0 |
| DG 14:1_18:2 | 0.0 | n.d. | n.d. | n.d. | 0.0 | n.d. | 0.0 | n.d. | 0.0 | n.d. | 0.0 | 0.0 | n.d. | n.d. | 0.0 | n.d. |
| DG 14:1_22:2 | n.d. | n.d. | n.d. | n.d. | n.d. | n.d. | n.d. | n.d. | n.d. | n.d. | n.d. | n.d. | n.d. | n.d. | n.d. | n.d. |
| DG 16:0_16:0 | 0.0 | 0.0 | 0.0 | 0.0 | 0.1 | 0.0 | 0.1 | 0.0 | 0.1 | 0.0 | 0.0 | 0.0 | 0.0 | 0.0 | 0.1 | 0.0 |
| DG 16:0_16:1 | 0.0 | 0.0 | 0.0 | 0.0 | 0.1 | 0.0 | 0.1 | 0.0 | 0.0 | 0.0 | 0.0 | 0.0 | 0.0 | 0.0 | 0.1 | 0.0 |
| DG 16:0_18:0 | 0.0 | 0.0 | 0.0 | 0.0 | 0.0 | 0.0 | 0.0 | 0.0 | 0.0 | 0.0 | 0.0 | 0.0 | 0.0 | 0.0 | 0.0 | 0.0 |
| DG 16:0_18:1 | 0.2 | 0.1 | 0.1 | 0.1 | 0.3 | 0.2 | 0.2 | 0.1 | 0.3 | 0.2 | 0.2 | 0.1 | 0.2 | 0.1 | 0.2 | 0.1 |
| DG 16:0_18:2 | 0.1 | 0.1 | 0.1 | 0.0 | 0.2 | 0.2 | 0.2 | 0.1 | 0.3 | 0.2 | 0.2 | 0.0 | 0.2 | 0.1 | 0.2 | 0.1 |
| DG 16:0_18:3 | 0.0 | 0.0 | 0.0 | 0.0 | 0.0 | 0.0 | 0.0 | 0.0 | 0.0 | 0.0 | 0.0 | 0.0 | 0.0 | 0.0 | 0.0 | 0.0 |
| DG 16:0_20:2 | 0.0 | n.d. | n.d. | n.d. | 0.0 | 0.0 | 0.0 | n.d. | 0.0 | n.d. | 0.0 | n.d. | n.d. | n.d. | 0.0 | n.d. |
| DG 16:0_20:3 | n.d. | n.d. | 0.0 | n.d. | 0.0 | 0.0 | 0.0 | n.d. | 0.0 | n.d. | n.d. | n.d. | n.d. | n.d. | 0.0 | n.d. |
| DG 16:0_20:4 | n.d. | n.d. | n.d. | n.d. | 0.0 | n.d. | n.d. | n.d. | 0.0 | n.d. | 0.0 | n.d. | 0.0 | 0.0 | 0.0 | 0.0 |

| | | | | | | | | | | | | | | | | |
|---|---|---|---|---|---|---|---|---|---|---|---|---|---|---|---|---|
| DG 16:0_22:4 | n.d. | n.d. | n.d. | n.d. | n.d. | n.d. | n.d. | n.d. | 0.0 | n.d. | n.d. | n.d. | 0.0 | 0.0 | 0.0 | n.d. |
| DG 16:0_22:5 | 0.0 | 0.0 | n.d. | n.d. | n.d. | n.d. | 0.0 | n.d. | n.d. | n.d. | n.d. | n.d. | n.d. | n.d. | 0.0 | 0.0 |
| DG 16:1_16:1 | 0.0 | 0.0 | 0.0 | 0.0 | 0.0 | 0.0 | 0.0 | 0.0 | 0.0 | 0.0 | 0.0 | 0.0 | 0.0 | 0.0 | 0.0 | 0.0 |
| DG 16:1_18:0 | 0.0 | 0.0 | 0.0 | 0.0 | 0.0 | 0.0 | 0.0 | n.d. | 0.0 | 0.0 | 0.0 | 0.0 | 0.0 | 0.0 | 0.0 | 0.0 |
| DG 16:1_18:1 | 0.0 | 0.0 | 0.0 | 0.0 | 0.1 | 0.1 | 0.0 | 0.0 | 0.0 | 0.0 | 0.0 | 0.0 | 0.0 | 0.0 | 0.1 | 0.0 |
| DG 16:1_18:2 | 0.0 | 0.0 | 0.0 | 0.0 | 0.0 | 0.1 | 0.0 | 0.0 | 0.0 | 0.0 | 0.1 | 0.0 | 0.0 | 0.0 | 0.0 | 0.0 |
| DG 16:1_18:3 | 0.0 | 0.0 | 0.0 | n.d. | 0.0 | 0.0 | 0.0 | 0.0 | 0.0 | 0.0 | 0.0 | 0.0 | 0.0 | 0.0 | 0.0 | 0.0 |
| DG 16:1_20:2 | n.d. | n.d. | n.d. | n.d. | 0.0 | n.d. | n.d. | n.d. | n.d. | n.d. | n.d. | n.d. | n.d. | n.d. | n.d. | n.d. |
| DG 16:1_20:3 | n.d. | n.d. | n.d. | n.d. | n.d. | n.d. | n.d. | n.d. | 0.0 | n.d. | n.d. | n.d. | n.d. | n.d. | n.d. | n.d. |
| DG 16:1_20:4 | 0.0 | n.d. | n.d. | n.d. | 0.0 | n.d. | 0.0 | n.d. | n.d. | n.d. | n.d. | n.d. | 0.0 | n.d. | n.d. | n.d. |
| DG 18:0_18:0 | 0.0 | 0.0 | 0.0 | 0.0 | 0.0 | 0.0 | 0.0 | 0.0 | 0.0 | 0.0 | 0.0 | 0.0 | 0.0 | 0.0 | 0.0 | 0.0 |
| DG 18:0_18:1 | 0.0 | 0.0 | 0.0 | 0.0 | 0.1 | 0.0 | 0.0 | 0.0 | 0.0 | 0.0 | 0.0 | 0.0 | 0.0 | 0.0 | 0.0 | 0.0 |
| DG 18:0_18:2 | 0.0 | 0.0 | 0.0 | 0.0 | 0.1 | 0.1 | 0.0 | 0.0 | 0.0 | 0.0 | 0.0 | 0.0 | 0.0 | 0.0 | 0.0 | 0.0 |
| DG 18:0_18:3 | 0.0 | n.d. | 0.0 | n.d. | 0.0 | 0.0 | 0.0 | n.d. | 0.0 | 0.0 | 0.0 | 0.0 | 0.0 | 0.0 | 0.0 | n.d. |
| DG 18:0_20:4 | n.d. | n.d. | n.d. | n.d. | n.d. | n.d. | n.d. | n.d. | 0.0 | n.d. | n.d. | n.d. | n.d. | n.d. | n.d. | n.d. |
| DG 18:1_18:1 | 0.1 | 0.1 | 0.1 | 0.1 | 0.2 | 0.2 | 0.1 | 0.1 | 0.2 | 0.2 | 0.1 | 0.0 | 0.1 | 0.1 | 0.2 | 0.1 |
| DG 18:1_18:2 | 0.2 | 0.1 | 0.1 | 0.1 | 0.3 | 0.3 | 0.2 | 0.1 | 0.3 | 0.3 | 0.3 | 0.1 | 0.2 | 0.2 | 0.3 | 0.1 |
| DG 18:1_18:3 | 0.0 | 0.0 | 0.0 | 0.0 | 0.0 | 0.0 | 0.0 | 0.0 | 0.0 | 0.0 | 0.0 | 0.0 | 0.0 | 0.0 | 0.0 | 0.0 |
| DG 18:1_18:4 | 0.0 | n.d. | n.d. | n.d. | 0.0 | n.d. | n.d. | n.d. | 0.0 | n.d. | 0.0 | n.d. | n.d. | n.d. | n.d. | n.d. |
| DG 18:1_20:2 | n.d. | n.d. | n.d. | n.d. | 0.0 | 0.0 | 0.0 | n.d. | 0.0 | 0.0 | 0.0 | n.d. | 0.0 | n.d. | 0.0 | n.d. |
| DG 18:1_20:3 | n.d. | n.d. | 0.0 | n.d. | 0.0 | n.d. | 0.0 | n.d. | 0.0 | 0.0 | 0.0 | 0.0 | 0.0 | 0.0 | 0.0 | n.d. |
| DG 18:1_20:4 | 0.0 | 0.0 | 0.0 | 0.0 | 0.0 | 0.0 | 0.0 | 0.0 | 0.0 | 0.0 | 0.0 | 0.0 | 0.0 | 0.0 | 0.0 | 0.0 |
| DG 18:1_22:5 | n.d. | n.d. | n.d. | n.d. | n.d. | n.d. | n.d. | n.d. | 0.0 | n.d. | 0.0 | n.d. | 0.0 | n.d. | n.d. | n.d. |
| DG 18:2_18:2 | 0.1 | 0.0 | 0.0 | 0.0 | 0.1 | 0.1 | 0.1 | 0.0 | 0.1 | 0.1 | 0.1 | 0.0 | 0.1 | 0.1 | 0.1 | 0.1 |
| DG 18:2_18:3 | 0.0 | 0.0 | 0.0 | 0.0 | 0.0 | 0.0 | 0.0 | 0.0 | 0.0 | 0.0 | 0.0 | 0.0 | 0.0 | 0.0 | 0.0 | 0.0 |

| | | | | | | | | | | | | | | | | |
|---|---|---|---|---|---|---|---|---|---|---|---|---|---|---|---|---|
| DG 18:2_20:2 | n.d. | n.d. | n.d. | n.d. | 0.0 | 0.0 | 0.0 | n.d. | 0.0 | n.d. | 0.0 | n.d. | n.d. | n.d. | 0.0 | n.d. |
| DG 18:2_20:3 | n.d. | n.d. | n.d. | n.d. | 0.0 | n.d. | 0.0 | 0.0 | 0.0 | 0.0 | n.d. | n.d. | 0.0 | n.d. | 0.0 | n.d. |
| DG 18:2_20:4 | 0.0 | n.d. | 0.0 | n.d. | 0.0 | 0.0 | n.d. | n.d. | n.d. | n.d. | 0.0 | 0.0 | 0.0 | 0.0 | 0.0 | 0.0 |
| DG 18:3_18:3 | 0.0 | n.d. | n.d. | n.d. | 0.0 | 0.0 | n.d. | n.d. | n.d. | n.d. | n.d. | n.d. | n.d. | n.d. | 0.0 | n.d. |
| DG total | 0.9 | 0.6 | 0.6 | 0.5 | 1.7 | 1.6 | 1.3 | 0.8 | 1.6 | 1.2 | 1.3 | 0.3 | 1.0 | 0.7 | 1.5 | 0.7 |

Lipidomics data were normalized using the ion abundance of internal standards added before lipid extraction via the modified Bligh and Dyer method.

n.d.: not detected.

### Supplementary Table 7 Quantitative lipidomics for LDL fraction.

| Lipids | Quantification (nmol mL$^{-1}$) | | | | | | | | | | | | | | | |
|---|---|---|---|---|---|---|---|---|---|---|---|---|---|---|---|---|
| | 8 months | | | | 12 months | | | | 16 months | | | | 20 months | | | |
| | Severe | | Mild | | Severe | | Mild | | Severe | | Mild | | Severe | | Mild | |
| | Mean | SD | Mean | SD | Mean | SD | Mean | SD | Mean | SD | Mean | SD | Mean | SD | Mean | SD |
| FA 14:0 | 10.9 | 6.3 | 3.3 | 3.6 | 5.5 | 5.7 | 5.9 | 5.1 | 11.2 | 6.8 | 6.4 | 5.3 | 5.6 | 3.4 | 3.5 | 4.9 |
| FA 16:0 | 60.3 | 46.8 | 53.7 | 34.6 | 80.6 | 18.7 | 98.0 | 25.2 | 75.6 | 8.9 | 91.9 | 16.4 | 53.8 | 49.2 | 68.5 | 11.8 |
| FA 16:1 | 1.0 | 1.1 | 2.7 | 0.8 | 2.4 | 1.0 | 2.1 | 1.8 | 4.0 | 3.8 | 6.0 | 2.8 | 4.2 | 2.7 | 3.0 | 1.9 |
| FA 18:0 | 80.1 | 20.0 | 55.1 | 35.6 | 85.2 | 10.8 | 81.8 | 10.1 | 64.6 | 32.8 | 68.8 | 43.1 | 89.8 | 19.4 | 86.1 | 10.9 |
| FA 18:1 | 10.8 | 4.5 | 10.5 | 7.7 | 11.9 | 9.6 | 8.8 | 6.9 | 9.5 | 8.3 | 15.3 | 10.3 | 15.0 | 11.3 | 14.3 | 0.8 |
| FA 18:2 | 7.2 | 1.6 | 4.6 | 0.7 | 5.4 | 3.9 | 5.0 | 2.0 | 5.9 | 3.9 | 7.0 | 0.9 | 6.5 | 5.5 | 5.1 | 1.3 |
| FA 18:3 | 1.9 | 0.5 | 2.0 | 0.4 | 1.2 | 0.9 | 1.8 | 0.6 | 1.9 | 1.0 | 1.5 | 1.0 | 0.9 | 0.8 | 1.1 | 0.4 |
| FA 20:4 | 0.6 | 0.3 | 0.3 | 0.3 | 0.4 | 0.1 | 0.4 | 0.6 | 0.5 | 0.4 | 0.3 | 0.1 | 0.5 | 0.4 | 0.3 | 0.4 |
| FA total | 172.8 | 33.4 | 132.2 | 59.3 | 192.6 | 36.6 | 203.3 | 22.4 | 173.3 | 33.7 | 197.3 | 60.9 | 176.5 | 85.0 | 181.8 | 8.5 |
| LPC 14:0 | 0.1 | 0.1 | 0.1 | 0.1 | 0.1 | 0.1 | 0.1 | 0.1 | 0.1 | 0.1 | 0.0 | 0.0 | 0.1 | 0.1 | 0.1 | 0.0 |
| LPC 16:0 | 100.7 | 10.5 | 87.6 | 13.7 | 90.7 | 14.1 | 76.8 | 11.2 | 77.1 | 15.9 | 68.5 | 1.5 | 93.2 | 36.6 | 65.2 | 3.8 |
| LPC 16:1 | 0.9 | 0.2 | 1.2 | 0.4 | 1.1 | 0.5 | 1.1 | 0.2 | 0.9 | 0.4 | 1.0 | 0.2 | 1.2 | 0.7 | 0.9 | 0.1 |
| LPC 18:0 | 227.8 | 20.6 | 195.0 | 17.6 | 198.9 | 15.6 | 194.2 | 8.2 | 159.7 | 16.3 | 158.3 | 14.7 | 186.3 | 29.3 | 155.9 | 26.0 |
| LPC 18:1 | 43.3 | 7.9 | 37.1 | 6.4 | 33.1 | 8.0 | 30.8 | 3.1 | 31.5 | 7.2 | 28.0 | 3.1 | 35.0 | 5.6 | 26.4 | 2.5 |
| LPC 18:2 | 68.4 | 7.2 | 60.5 | 6.6 | 56.5 | 11.3 | 51.8 | 3.7 | 50.1 | 8.0 | 48.6 | 5.9 | 51.9 | 9.7 | 40.6 | 9.4 |
| LPC 18:3 | 1.3 | 0.4 | 1.1 | 0.2 | 1.1 | 0.5 | 1.1 | 0.3 | 1.1 | 0.5 | 1.0 | 0.2 | 0.9 | 0.3 | 0.7 | 0.3 |
| LPC 20:0 | 1.2 | 0.1 | 1.2 | 0.2 | 1.0 | 0.2 | 1.0 | 0.2 | 0.7 | 0.3 | 0.7 | 0.2 | 0.8 | 0.2 | 0.7 | 0.1 |
| LPC 20:1 | 1.0 | 0.2 | 0.9 | 0.3 | 0.8 | 0.3 | 0.9 | 0.2 | 0.9 | 0.1 | 0.8 | 0.3 | 0.9 | 0.2 | 0.8 | 0.3 |
| LPC 20:2 | 2.0 | 0.5 | 1.4 | 0.7 | 1.4 | 0.5 | 1.1 | 0.3 | 1.5 | 0.3 | 1.1 | 0.4 | 1.4 | 0.2 | 1.0 | 0.4 |
| LPC 20:3 | 1.7 | 0.4 | 1.8 | 0.6 | 1.4 | 0.5 | 1.5 | 0.3 | 1.4 | 0.5 | 1.3 | 0.3 | 1.5 | 0.4 | 1.2 | 0.2 |

| | | | | | | | | | | | | | | | | |
|---|---|---|---|---|---|---|---|---|---|---|---|---|---|---|---|---|
| LPC 20:4 | 3.6 | 0.3 | 3.5 | 0.5 | 3.4 | 0.8 | 4.7 | 2.0 | 3.5 | 2.0 | 3.1 | 0.9 | 3.4 | 2.2 | 2.8 | 0.5 |
| LPC 20:5 | 0.0 | 0.0 | 0.1 | 0.0 | 0.0 | 0.0 | 0.0 | 0.0 | 0.1 | 0.1 | 0.0 | 0.0 | 0.1 | 0.1 | 0.0 | 0.0 |
| LPC 22:0 | 0.3 | 0.2 | 0.2 | 0.0 | 0.2 | 0.1 | 0.1 | 0.0 | 0.1 | 0.1 | 0.1 | 0.1 | 0.2 | 0.1 | 0.2 | 0.1 |
| LPC 22:4 | 0.5 | 0.2 | 0.5 | 0.2 | 0.4 | 0.1 | 0.4 | 0.1 | 0.5 | 0.3 | 0.4 | 0.2 | 0.5 | 0.3 | 0.4 | 0.2 |
| LPC 22:5 | 0.6 | 0.1 | 0.7 | 0.1 | 0.4 | 0.1 | 0.6 | 0.1 | 0.5 | 0.4 | 0.4 | 0.1 | 0.5 | 0.2 | 0.4 | 0.1 |
| LPC 22:6 | 0.1 | 0.0 | 0.2 | 0.1 | 0.2 | 0.1 | 0.1 | 0.1 | 0.1 | 0.0 | 0.1 | 0.1 | 0.2 | 0.1 | 0.1 | 0.1 |
| LPC total | 453.4 | 40.7 | 393.1 | 44.8 | 390.6 | 48.5 | 366.3 | 19.7 | 329.7 | 40.3 | 313.6 | 24.1 | 378.2 | 77.8 | 297.5 | 41.4 |
| LPE 16:0 | 1.5 | 0.8 | 1.2 | 0.7 | 0.7 | 0.3 | 0.8 | 0.6 | 1.0 | 1.1 | 0.4 | 0.2 | 1.3 | 1.1 | 1.0 | 1.0 |
| LPE 18:0 | 7.4 | 2.5 | 5.3 | 1.0 | 4.7 | 1.7 | 4.8 | 2.9 | 4.6 | 2.0 | 4.7 | 1.5 | 6.0 | 3.3 | 3.3 | 2.2 |
| LPE 18:1 | 2.2 | 0.9 | 2.2 | 1.2 | 1.1 | 0.8 | 1.2 | 0.5 | 1.4 | 1.0 | 0.1 | 0.1 | 1.5 | 1.0 | 0.5 | 0.4 |
| LPE 18:2 | 5.9 | 1.9 | 3.5 | 1.3 | 3.9 | 1.1 | 9.3 | 11.2 | 7.4 | 10.9 | 2.1 | 0.6 | 4.0 | 1.1 | 2.1 | 0.7 |
| LPE total | 16.7 | 4.6 | 12.1 | 3.6 | 10.4 | 2.8 | 15.8 | 10.6 | 14.2 | 10.0 | 7.2 | 1.7 | 12.6 | 5.7 | 6.8 | 2.9 |
| PC 16:0_16:0 | 64.2 | 14.3 | 53.9 | 10.6 | 41.3 | 12.1 | 35.2 | 6.5 | 38.1 | 10.4 | 36.9 | 2.7 | 43.9 | 18.9 | 32.0 | 4.0 |
| PC 16:0_16:1 | 8.0 | 3.5 | 5.8 | 2.2 | 3.1 | 2.8 | 6.3 | 1.1 | 4.1 | 3.3 | 6.5 | 2.1 | 7.8 | 6.4 | 4.3 | 1.5 |
| PC 16:0_18:0 | 207.0 | 36.7 | 186.4 | 33.7 | 138.5 | 27.3 | 121.1 | 17.1 | 130.0 | 29.5 | 109.5 | 23.7 | 180.8 | 48.7 | 119.4 | 13.6 |
| PC 16:0_18:1 | 1007.6 | 167.2 | 909.3 | 122.6 | 675.5 | 155.3 | 567.8 | 111.5 | 682.6 | 141.3 | 570.1 | 98.1 | 862.7 | 175.5 | 608.7 | 62.3 |
| PC 16:0_18:2 | 1668.4 | 203.6 | 1520.5 | 200.1 | 1178.8 | 198.7 | 975.4 | 184.3 | 1121.4 | 130.5 | 1020.8 | 82.4 | 1318.3 | 237.5 | 1010.8 | 175.4 |
| PC 16:0_18:3 | 61.9 | 12.2 | 54.7 | 15.0 | 32.7 | 12.7 | 29.4 | 12.2 | 27.3 | 7.3 | 28.6 | 7.4 | 37.3 | 5.5 | 22.5 | 7.2 |
| PC 16:0_20:2 | 16.6 | 8.1 | 13.6 | 4.3 | 13.4 | 3.6 | 11.1 | 3.8 | 14.2 | 3.7 | 10.2 | 4.1 | 18.0 | 6.6 | 15.2 | 1.4 |
| PC 16:0_20:3 | 63.1 | 13.4 | 59.5 | 14.4 | 40.9 | 8.8 | 40.2 | 6.3 | 36.1 | 7.9 | 34.8 | 6.2 | 55.6 | 36.5 | 46.6 | 18.8 |
| PC 16:0_20:4 | 83.6 | 12.6 | 77.6 | 9.1 | 58.3 | 18.3 | 54.0 | 12.2 | 46.6 | 9.7 | 52.2 | 16.5 | 81.5 | 40.7 | 64.3 | 15.7 |
| PC 16:0_22:4 | 10.5 | 4.0 | 11.4 | 2.9 | 7.4 | 1.9 | 8.2 | 3.1 | 4.5 | 1.0 | 5.5 | 4.3 | 14.3 | 11.7 | 10.7 | 2.3 |
| PC 16:0_22:5 | 10.4 | 3.0 | 9.7 | 4.8 | 7.9 | 6.2 | 7.3 | 2.4 | 5.5 | 1.2 | 5.4 | 1.1 | 13.1 | 14.6 | 7.8 | 3.3 |
| PC 16:1_18:1 | 8.1 | 4.2 | 9.6 | 3.2 | 6.7 | 4.8 | 6.7 | 3.0 | 5.6 | 3.3 | 4.1 | 3.5 | 8.1 | 3.4 | 7.1 | 2.3 |
| PC 16:1_18:2 | 18.8 | 6.0 | 22.1 | 6.8 | 12.1 | 4.9 | 13.2 | 3.4 | 12.1 | 5.5 | 12.6 | 5.8 | 16.4 | 6.0 | 17.3 | 7.2 |

| | | | | | | | | | | | | | | | | |
|---|---|---|---|---|---|---|---|---|---|---|---|---|---|---|---|---|
| PC 18:0_18:0 | 93.1 | 23.0 | 75.6 | 7.6 | 65.6 | 12.0 | 63.4 | 8.0 | 62.2 | 13.1 | 52.7 | 9.7 | 74.3 | 12.7 | 64.0 | 8.7 |
| PC 18:0_18:1 | 779.1 | 92.2 | 647.3 | 77.3 | 553.7 | 104.0 | 555.4 | 46.1 | 506.0 | 61.5 | 515.7 | 88.4 | 635.3 | 96.1 | 523.8 | 94.9 |
| PC 18:0_18:2 | 1858.8 | 108.0 | 1641.4 | 193.4 | 1358.7 | 165.9 | 1375.1 | 133.0 | 1266.6 | 162.9 | 1307.3 | 193.1 | 1460.6 | 188.8 | 1303.7 | 273.5 |
| PC 18:0_18:3 | 38.8 | 10.3 | 37.6 | 3.7 | 25.3 | 10.8 | 26.2 | 5.3 | 20.8 | 8.3 | 21.8 | 11.2 | 28.4 | 6.4 | 23.5 | 11.8 |
| PC 18:0_20:2 | 27.9 | 10.9 | 17.6 | 4.2 | 15.1 | 4.2 | 17.5 | 4.8 | 14.6 | 7.4 | 12.3 | 4.6 | 22.9 | 6.5 | 17.9 | 3.3 |
| PC 18:0_20:3 | 80.1 | 12.0 | 75.3 | 11.5 | 55.2 | 17.7 | 68.6 | 17.5 | 47.1 | 6.4 | 59.1 | 13.8 | 72.9 | 39.2 | 77.6 | 15.6 |
| PC 18:0_20:4 | 112.7 | 15.5 | 108.3 | 11.8 | 93.8 | 24.4 | 96.9 | 16.7 | 67.8 | 10.4 | 91.0 | 20.2 | 107.7 | 52.7 | 105.1 | 14.9 |
| PC 18:0_22:4 | 7.6 | 2.9 | 7.9 | 3.5 | 8.2 | 1.3 | 4.3 | 1.3 | 6.6 | 4.0 | 5.3 | 1.5 | 6.2 | 7.3 | 8.8 | 4.9 |
| PC 18:0_22:5 | 7.2 | 3.5 | 7.7 | 1.4 | 5.7 | 1.5 | 4.4 | 1.8 | 3.7 | 1.1 | 6.6 | 2.6 | 6.6 | 8.9 | 7.0 | 2.0 |
| PC 18:1_18:1 | 105.5 | 11.6 | 76.3 | 14.0 | 69.7 | 23.0 | 60.6 | 12.5 | 70.1 | 12.1 | 66.5 | 12.6 | 81.7 | 17.2 | 73.3 | 14.6 |
| PC 18:1_18:2 | 454.7 | 55.3 | 392.5 | 67.7 | 326.2 | 72.3 | 294.5 | 59.7 | 315.0 | 51.2 | 310.7 | 46.6 | 342.7 | 68.1 | 280.1 | 70.2 |
| PC 18:1_18:3 | 18.2 | 4.9 | 17.7 | 5.1 | 8.5 | 3.2 | 8.2 | 4.8 | 7.5 | 3.4 | 8.4 | 1.1 | 12.8 | 5.1 | 7.1 | 3.9 |
| PC 18:1_20:2 | 3.3 | 2.9 | 4.2 | 4.8 | 2.5 | 1.8 | 1.4 | 1.7 | 5.2 | 1.4 | 1.9 | 0.8 | 3.9 | 1.5 | 2.4 | 2.6 |
| PC 18:1_20:4 | 12.3 | 3.6 | 8.3 | 2.4 | 7.4 | 3.2 | 9.3 | 2.8 | 5.2 | 3.9 | 6.7 | 4.4 | 10.2 | 5.3 | 9.8 | 3.7 |
| PC 18:2_18:2 | 334.4 | 58.8 | 289.6 | 39.2 | 228.7 | 49.4 | 200.2 | 29.1 | 205.0 | 38.8 | 198.0 | 24.5 | 215.9 | 56.2 | 170.2 | 50.1 |
| PC 18:2_18:3 | 25.0 | 3.7 | 24.1 | 5.7 | 12.6 | 5.3 | 16.8 | 6.4 | 9.4 | 5.2 | 13.9 | 7.9 | 11.1 | 6.2 | 9.8 | 5.2 |
| PC 18:2_20:1 | 7.5 | 5.7 | 7.5 | 5.9 | 5.3 | 2.1 | 7.9 | 1.9 | 7.1 | 2.0 | 6.6 | 5.0 | 6.2 | 4.2 | 6.5 | 5.8 |
| PC 18:2_20:2 | 12.7 | 3.7 | 15.6 | 7.5 | 10.1 | 3.0 | 10.3 | 1.5 | 8.3 | 1.4 | 10.0 | 3.9 | 8.4 | 5.1 | 13.4 | 9.5 |
| PC 18:2_20:3 | 20.6 | 6.0 | 18.0 | 8.9 | 13.3 | 6.0 | 11.7 | 2.8 | 9.6 | 3.5 | 12.0 | 5.8 | 11.8 | 2.8 | 17.4 | 4.4 |
| PC 18:2_20:4 | 17.0 | 3.9 | 16.4 | 6.9 | 11.0 | 5.2 | 12.5 | 2.2 | 11.4 | 2.6 | 12.5 | 8.0 | 12.9 | 3.6 | 10.6 | 4.2 |
| PC total | 7244.6 | 782.6 | 6423.2 | 805.2 | 5093.1 | 809.4 | 4721.0 | 581.3 | 4774.1 | 559.5 | 4615.9 | 611.0 | 5790.2 | 690.4 | 4698.7 | 777.1 |
| PC O-16:1/16:0 (PC P-16:0/16:0) | 9.7 | 10.1 | 10.9 | 3.7 | 6.8 | 4.0 | 11.0 | 5.1 | 4.1 | 1.1 | 6.8 | 3.1 | 7.0 | 4.4 | 5.4 | 4.5 |
| PC O-16:1/18:1 (PC P-16:0/18:1) | 43.2 | 22.6 | 60.0 | 11.5 | 39.4 | 8.2 | 29.9 | 11.3 | 23.8 | 12.7 | 30.3 | 11.3 | 37.4 | 6.8 | 32.0 | 15.9 |
| PC O-16:1/18:2 (PC P-16:0/18:2) | 51.0 | 12.9 | 40.0 | 7.2 | 15.8 | 6.7 | 27.8 | 14.0 | 28.8 | 15.4 | 19.3 | 12.7 | 38.8 | 5.2 | 26.0 | 12.8 |
| PC O-16:1/20:4 (PC P-16:0/20:4) | 37.9 | 11.5 | 45.0 | 11.3 | 34.2 | 7.0 | 31.6 | 17.7 | 24.1 | 19.3 | 26.5 | 15.2 | 36.5 | 19.0 | 39.1 | 10.5 |

| | | | | | | | | | | | | | | | | |
|---|---|---|---|---|---|---|---|---|---|---|---|---|---|---|---|---|
| PC O-18:1/18:2 (PC P-18:0/18:2) | 10.0 | 4.7 | 10.5 | 10.4 | 7.6 | 6.0 | 5.2 | 4.0 | 5.7 | 1.4 | 7.0 | 2.9 | 7.2 | 4.0 | 4.6 | 4.9 |
| PC O-18:1/20:4 (PC P-18:0/20:4) | 11.2 | 4.7 | 10.1 | 6.1 | 5.8 | 3.3 | 6.1 | 3.3 | 5.3 | 3.4 | 8.6 | 6.9 | 10.0 | 8.1 | 5.8 | 0.8 |
| PC P-18:2/20:5 | 120.6 | 41.1 | 96.1 | 16.6 | 70.2 | 16.2 | 64.6 | 23.5 | 64.1 | 22.5 | 82.3 | 47.0 | 116.1 | 16.1 | 86.3 | 23.8 |
| EtherPC total | 282.2 | 74.4 | 277.2 | 46.5 | 183.2 | 18.5 | 172.1 | 72.1 | 155.0 | 66.8 | 181.5 | 89.9 | 250.2 | 31.2 | 199.6 | 56.2 |
| PE 14:1_18:2 | 2.1 | 1.8 | 2.4 | 0.9 | 1.4 | 0.9 | 1.3 | 0.7 | 2.7 | 2.2 | 1.1 | 0.4 | 0.6 | 0.4 | 0.9 | 1.1 |
| PE 16:0_18:1 | 5.1 | 3.7 | 4.2 | 1.0 | 3.1 | 2.9 | 1.8 | 2.0 | 3.3 | 1.5 | 1.7 | 0.8 | 3.1 | 2.4 | 1.6 | 1.2 |
| PE 16:0_18:2 | 28.9 | 9.8 | 21.4 | 3.5 | 17.8 | 7.9 | 11.6 | 3.8 | 15.9 | 5.4 | 9.3 | 2.1 | 21.3 | 13.2 | 6.1 | 3.5 |
| PE 16:0_20:4 | 4.0 | 3.5 | 3.9 | 1.0 | 2.4 | 1.4 | 2.9 | 1.8 | 2.1 | 1.1 | 1.4 | 0.9 | 3.0 | 2.4 | 1.3 | 1.1 |
| PE 16:1_18:2 | 12.0 | 3.1 | 10.7 | 4.2 | 6.6 | 3.2 | 5.4 | 2.7 | 4.8 | 2.4 | 7.1 | 3.1 | 6.1 | 3.0 | 4.8 | 2.4 |
| PE 16:1_20:4 | 1.0 | 0.6 | 2.0 | 2.4 | 0.7 | 0.3 | 1.3 | 0.3 | 0.8 | 0.7 | 0.8 | 0.4 | 1.1 | 0.7 | 0.7 | 0.5 |
| PE 18:0_18:1 | 11.2 | 4.8 | 7.5 | 2.2 | 6.8 | 5.8 | 4.1 | 2.6 | 6.0 | 3.4 | 3.4 | 1.5 | 8.3 | 5.4 | 3.4 | 1.3 |
| PE 18:0_18:2 | 102.5 | 27.8 | 84.5 | 10.6 | 71.4 | 30.0 | 48.0 | 9.9 | 61.2 | 12.2 | 47.9 | 9.1 | 75.8 | 29.6 | 42.2 | 14.8 |
| PE 18:0_18:3 | 1.4 | 0.6 | 2.4 | 1.7 | 0.8 | 0.4 | 0.7 | 0.4 | 0.7 | 0.6 | 0.8 | 0.2 | 1.0 | 0.8 | 0.4 | 0.3 |
| PE 18:0_20:3 | 0.8 | 0.6 | 1.1 | 1.0 | 0.7 | 0.6 | 1.2 | 1.4 | 0.3 | 0.2 | 0.5 | 0.6 | 1.1 | 1.0 | 0.6 | 0.3 |
| PE 18:0_20:4 | 30.5 | 11.7 | 29.4 | 4.7 | 23.1 | 8.2 | 18.5 | 4.7 | 16.5 | 5.4 | 14.6 | 2.5 | 27.9 | 13.2 | 14.4 | 3.7 |
| PE 18:0_22:4 | 1.0 | 0.8 | 0.5 | 0.3 | 0.6 | 0.3 | 0.4 | 0.3 | 0.9 | 0.6 | 0.2 | 0.1 | 0.4 | 0.3 | 0.3 | 0.1 |
| PE 18:1_18:1 | 3.1 | 2.6 | 2.3 | 0.5 | 1.3 | 0.7 | 0.7 | 0.4 | 1.5 | 1.3 | 0.9 | 0.1 | 2.4 | 1.9 | 0.8 | 0.3 |
| PE 18:1_18:2 | 17.5 | 6.1 | 13.2 | 1.8 | 10.3 | 5.9 | 6.1 | 4.7 | 7.1 | 5.0 | 5.9 | 1.9 | 12.1 | 7.0 | 6.9 | 2.2 |
| PE 18:1_20:4 | 2.3 | 1.4 | 0.6 | 0.2 | 1.0 | 1.0 | 0.7 | 0.2 | 0.9 | 0.6 | 0.6 | 0.5 | 1.0 | 0.6 | 0.7 | 0.9 |
| PE 18:2_18:2 | 2.7 | 1.7 | 1.7 | 0.8 | 1.5 | 1.7 | 1.8 | 0.5 | 1.7 | 1.3 | 1.7 | 0.8 | 2.0 | 1.5 | 0.7 | 0.6 |
| PE 18:2_20:0 | 0.8 | 0.7 | 1.6 | 1.0 | 0.7 | 0.6 | 1.0 | 0.2 | 0.6 | 0.3 | 0.2 | 0.2 | 0.2 | 0.2 | 1.0 | 0.9 |
| PE 18:2_20:1 | 1.7 | 0.8 | 2.3 | 1.5 | 1.1 | 0.6 | 1.0 | 1.0 | 0.4 | 0.4 | 1.0 | 0.3 | 1.2 | 0.8 | 1.8 | 1.2 |
| PE total | 228.2 | 73.3 | 190.9 | 28.1 | 150.3 | 63.7 | 105.4 | 34.9 | 126.2 | 33.3 | 97.7 | 6.5 | 168.0 | 75.2 | 86.3 | 31.3 |
| PE P-16:0/16:0 | 2.4 | 1.2 | 1.5 | 0.8 | 2.1 | 1.1 | 1.8 | n.d. | 1.2 | n.d. | 0.8 | 0.4 | 1.2 | n.d. | 0.7 | n.d. |
| PE P-16:0/18:1 | 33.6 | 14.4 | 36.7 | 2.2 | 19.4 | 5.4 | 19.0 | 11.2 | 20.2 | 9.0 | 23.3 | 5.4 | 24.1 | 8.4 | 18.5 | 6.0 |

| | | | | | | | | | | | | | | | | |
|---|---|---|---|---|---|---|---|---|---|---|---|---|---|---|---|---|
| PE P-16:0/18:2 | 228.1 | 85.0 | 199.1 | 29.8 | 146.6 | 27.8 | 135.8 | 42.6 | 122.2 | 51.4 | 129.2 | 35.5 | 116.5 | 55.0 | 102.4 | 26.9 |
| PE P-16:0/18:3 | 11.0 | 7.0 | 8.0 | 1.7 | 2.7 | 2.4 | 3.9 | 3.0 | 5.8 | 4.0 | 3.5 | 1.6 | 5.1 | 2.0 | 4.0 | 4.1 |
| PE P-16:0/20:3 | 2.0 | 0.7 | 2.6 | 2.0 | 1.8 | 0.9 | 1.7 | 0.5 | 1.9 | 0.5 | 2.0 | 1.1 | 1.2 | 0.6 | 1.9 | 1.3 |
| PE P-16:0/20:4 | 47.3 | 5.9 | 37.1 | 5.2 | 28.3 | 6.6 | 33.6 | 11.1 | 22.7 | 6.3 | 27.0 | 3.7 | 39.5 | 19.9 | 35.0 | 1.6 |
| PE P-16:0/22:4 | 12.5 | 4.4 | 8.9 | 3.4 | 7.4 | 4.4 | 4.6 | 3.8 | 9.4 | 3.7 | 5.5 | 3.4 | 9.8 | 4.1 | 6.7 | 2.1 |
| PE P-16:0/22:5 | 28.3 | 6.5 | 21.7 | 2.1 | 16.8 | 4.0 | 13.9 | 0.4 | 18.5 | 8.4 | 11.3 | 5.3 | 31.0 | 13.4 | 13.6 | 4.3 |
| PE P-16:0/22:6 | 11.7 | 4.4 | 3.9 | 1.8 | 4.3 | 1.5 | 2.2 | 1.5 | 5.7 | 4.6 | 2.2 | 2.2 | 7.5 | 4.3 | 8.3 | 5.7 |
| PE P-16:1/22:2 | 2.6 | 2.0 | 1.7 | 1.2 | 0.9 | 0.6 | 0.6 | 0.1 | 1.1 | 1.0 | 1.0 | 1.2 | 1.7 | 1.4 | 1.5 | 0.9 |
| PE P-18:0/16:0 | 3.1 | 4.1 | 4.8 | 3.7 | 3.9 | 3.1 | 2.9 | 1.0 | 6.1 | 2.5 | 1.8 | 1.8 | 2.4 | 0.9 | 2.8 | 2.2 |
| PE P-18:0/18:0 | 2.0 | 1.2 | 0.5 | 0.0 | 0.5 | 0.2 | 1.8 | 1.7 | 0.3 | 0.2 | 0.7 | 0.3 | 1.8 | 0.5 | 0.6 | 0.7 |
| PE P-18:0/18:1 | 74.0 | 24.3 | 67.6 | 15.2 | 51.4 | 11.9 | 45.1 | 9.7 | 41.4 | 19.6 | 50.6 | 8.3 | 45.3 | 24.8 | 36.1 | 9.1 |
| PE P-18:0/18:2 | 681.4 | 170.6 | 629.0 | 92.6 | 488.8 | 78.5 | 400.3 | 128.0 | 390.9 | 123.5 | 379.7 | 113.8 | 462.6 | 176.7 | 383.2 | 80.5 |
| PE P-18:0/18:3 | 31.7 | 6.9 | 31.2 | 4.7 | 16.3 | 5.0 | 18.2 | 10.6 | 13.8 | 9.0 | 15.1 | 7.1 | 23.2 | 10.1 | 17.8 | 4.8 |
| PE P-18:0/20:3 | 5.3 | 4.3 | 4.4 | 3.3 | 3.5 | 2.4 | 3.6 | 0.5 | 3.8 | 2.2 | 2.5 | 1.4 | 5.9 | 6.1 | 3.9 | 3.2 |
| PE P-18:0/20:4 | 74.4 | 18.8 | 80.7 | 7.9 | 55.7 | 16.5 | 46.4 | 7.7 | 44.8 | 17.5 | 55.2 | 7.6 | 68.8 | 44.5 | 62.6 | 13.4 |
| PE P-18:0/22:4 | 4.0 | 2.3 | 5.0 | 1.9 | 4.0 | 2.4 | 3.4 | 1.0 | 4.1 | 2.6 | 3.2 | 2.4 | 8.3 | 2.8 | 4.5 | 3.3 |
| PE P-18:0/22:5 | 20.4 | 7.7 | 15.2 | 4.6 | 11.2 | 6.7 | 14.2 | 6.5 | 11.1 | 4.1 | 10.3 | 4.3 | 14.8 | 11.5 | 15.3 | 6.7 |
| PE P-18:0/22:6 | 13.1 | 4.9 | 12.4 | 7.2 | 5.0 | 2.5 | 7.4 | 3.0 | 6.1 | 2.4 | 5.8 | 4.9 | 9.2 | 4.3 | 8.3 | 3.9 |
| PE P-18:1/16:0 | 8.8 | 4.8 | 8.4 | 2.2 | 4.7 | 2.9 | 7.3 | 2.3 | 6.3 | 7.2 | 6.7 | 2.2 | 7.4 | 7.2 | 5.8 | 3.3 |
| PE P-18:1/18:1 | 40.7 | 13.0 | 42.0 | 13.8 | 25.4 | 10.8 | 31.4 | 3.8 | 30.5 | 13.5 | 31.0 | 15.5 | 31.3 | 9.0 | 21.2 | 8.4 |
| PE P-18:1/18:2 | 231.3 | 88.6 | 218.1 | 33.9 | 166.4 | 42.7 | 171.2 | 27.5 | 152.0 | 76.2 | 157.5 | 42.7 | 179.3 | 103.6 | 159.8 | 61.0 |
| PE P-18:1/18:3 | 9.1 | 5.3 | 7.3 | 3.4 | 6.1 | 3.0 | 5.8 | 1.7 | 4.2 | 3.2 | 5.7 | 2.3 | 5.6 | 4.2 | 6.7 | 4.1 |
| PE P-18:1/20:3 | 1.8 | 1.8 | 1.0 | 1.1 | 1.6 | 1.0 | 1.5 | 0.5 | 1.5 | 1.1 | 0.9 | 0.5 | 3.0 | 2.5 | 1.2 | 0.8 |
| PE P-18:1/20:4 | 33.9 | 11.8 | 31.7 | 7.9 | 16.5 | 3.7 | 21.5 | 3.9 | 17.7 | 3.3 | 21.9 | 8.9 | 31.9 | 20.2 | 26.6 | 2.0 |
| PE P-18:1/22:4 | 2.4 | 1.7 | 2.0 | 2.0 | 3.3 | 3.8 | 1.0 | 0.8 | 0.7 | 0.2 | 1.4 | 1.3 | 3.4 | 3.4 | 1.2 | 0.8 |

| | | | | | | | | | | | | | | | | |
|---|---|---|---|---|---|---|---|---|---|---|---|---|---|---|---|---|
| PE P-18:1/22:5 | 5.8 | 5.5 | 7.8 | 0.6 | 4.4 | 2.7 | 3.7 | 2.5 | 3.4 | 2.4 | 6.2 | 3.9 | 6.4 | 4.7 | 3.5 | 1.5 |
| PE P-18:1/22:6 | 2.4 | 1.8 | 2.5 | 0.9 | 0.9 | 0.8 | 1.2 | 1.1 | 0.3 | n.d. | 2.4 | 1.4 | 1.8 | 2.4 | 1.3 | 0.3 |
| PE P-18:2/18:1 | 2.5 | 2.1 | 2.0 | 1.9 | 1.2 | 1.3 | 1.9 | 0.2 | 1.1 | 1.6 | 0.9 | 0.9 | 0.5 | 0.0 | 1.0 | 0.4 |
| PE P-18:2/18:2 | 14.1 | 10.1 | 19.3 | 7.7 | 7.7 | 5.8 | 8.0 | 2.2 | 9.2 | 7.0 | 11.0 | 8.2 | 7.8 | 6.2 | 8.7 | 7.7 |
| PE P-18:2/20:4 | 1.3 | 0.5 | 1.1 | 1.0 | 1.9 | n.d. | 1.6 | 0.6 | 1.5 | 0.1 | 1.2 | 1.1 | 2.4 | 1.4 | 1.1 | 0.2 |
| EtherPE total | 1636.2 | 448.2 | 1511.2 | 119.4 | 1103.5 | 179.1 | 1007.8 | 228.8 | 946.7 | 344.9 | 972.2 | 205.1 | 1151.9 | 413.1 | 956.4 | 222.6 |
| PI 16:0_16:0 | 175.4 | 70.1 | 200.9 | 169.0 | 80.0 | 43.8 | 80.0 | 57.7 | 85.4 | 71.5 | 138.0 | 81.9 | 124.6 | 97.7 | 107.5 | 44.2 |
| PI 16:0_18:0 | 176.1 | 54.9 | 185.7 | 165.1 | 50.4 | 23.8 | 54.1 | 35.8 | 37.5 | 28.4 | 53.7 | 42.5 | 90.0 | 74.5 | 68.2 | 19.0 |
| PI 16:0_18:1 | 1660.3 | 534.7 | 1705.8 | 703.4 | 853.4 | 278.7 | 727.3 | 255.3 | 715.0 | 212.5 | 591.4 | 174.3 | 1137.9 | 469.8 | 856.0 | 339.1 |
| PI 16:0_18:2 | 3227.1 | 651.3 | 3989.5 | 1214.7 | 1697.8 | 256.2 | 1595.2 | 757.6 | 1698.0 | 326.7 | 1724.8 | 561.5 | 1968.0 | 670.8 | 1299.6 | 581.8 |
| PI 16:0_20:4 | 228.8 | 108.6 | 259.5 | 108.2 | 143.6 | 96.7 | 88.8 | 75.1 | 72.7 | 64.2 | 29.9 | 16.5 | 102.6 | 88.2 | 134.2 | 119.2 |
| PI 16:1_18:0 | 129.7 | 74.0 | 161.0 | 88.3 | 97.6 | 89.5 | 99.7 | 61.4 | 79.7 | 91.4 | 115.3 | 98.2 | 145.6 | 137.9 | 103.3 | 72.8 |
| PI 18:0_18:0 | 62.0 | 51.7 | 76.8 | 95.8 | 79.2 | 39.5 | 40.0 | 26.6 | 33.2 | 24.1 | 43.1 | 13.2 | 71.9 | 72.5 | 58.6 | 56.4 |
| PI 18:0_18:1 | 5122.5 | 1125.3 | 6340.0 | 2179.4 | 3413.8 | 913.7 | 3189.1 | 350.3 | 2694.5 | 878.2 | 2740.0 | 1047.7 | 4202.1 | 2281.5 | 3164.8 | 942.7 |
| PI 18:0_18:2 | 15646.6 | 2547.5 | 18469.9 | 6469.6 | 10601.2 | 1706.3 | 10363.3 | 2138.3 | 9303.0 | 785.8 | 9167.3 | 1963.8 | 10556.3 | 2278.6 | 8913.4 | 3356.3 |
| PI 18:0_18:3 | 60.2 | 33.8 | 177.1 | 88.7 | 48.8 | 62.4 | 80.4 | 39.0 | 33.1 | 9.9 | 24.4 | 10.7 | 48.8 | 21.8 | 52.5 | 12.5 |
| PI 18:0_20:2 | 1577.8 | 738.0 | 1092.9 | 879.9 | 977.6 | 341.4 | 668.7 | 339.4 | 687.1 | 308.9 | 468.6 | 316.8 | 843.9 | 452.4 | 505.4 | 440.6 |
| PI 18:0_20:3 | 1675.8 | 279.2 | 1546.4 | 510.4 | 1051.4 | 411.0 | 769.8 | 257.8 | 586.6 | 156.0 | 572.4 | 140.7 | 791.3 | 152.8 | 775.6 | 258.0 |
| PI 18:0_20:4 | 5455.9 | 138.6 | 6079.4 | 1188.2 | 3884.4 | 1346.9 | 3172.3 | 438.5 | 2799.2 | 542.0 | 3087.3 | 227.3 | 3939.7 | 799.1 | 3326.3 | 910.2 |
| PI 18:1_18:1 | 283.4 | 96.2 | 377.4 | 264.2 | 152.7 | 68.5 | 112.5 | 39.7 | 148.3 | 104.3 | 116.4 | 67.7 | 205.4 | 77.2 | 112.7 | 55.5 |
| PI 18:1_18:2 | 294.2 | 98.8 | 484.8 | 280.5 | 213.0 | 89.5 | 230.4 | 181.6 | 143.3 | 58.4 | 179.4 | 83.2 | 261.3 | 158.2 | 206.3 | 178.2 |
| PI total | 35736.1 | 5417.8 | 41147.2 | 13411.5 | 23345.1 | 4435.6 | 21238.1 | 4075.7 | 19094.0 | 2254.2 | 19000.5 | 3185.3 | 24479.7 | 6499.7 | 19641.1 | 6492.3 |
| SM 18:1;O2/14:0 | 0.6 | 0.1 | 0.6 | 0.1 | 0.6 | 0.1 | 0.6 | 0.0 | 0.6 | 0.1 | 0.5 | 0.0 | 0.6 | 0.1 | 0.6 | 0.1 |
| SM 18:1;O2/14:1 | 0.0 | 0.0 | 0.0 | 0.0 | 0.0 | 0.0 | 0.0 | 0.0 | 0.0 | 0.0 | 0.0 | 0.0 | 0.0 | 0.0 | 0.0 | 0.0 |
| SM 18:1;O2/16:0 | 61.6 | 3.8 | 57.7 | 1.6 | 55.4 | 3.3 | 54.8 | 3.3 | 52.1 | 6.4 | 51.4 | 3.9 | 60.0 | 7.6 | 54.2 | 6.3 |

| | | | | | | | | | | | | | | | | |
|---|---|---|---|---|---|---|---|---|---|---|---|---|---|---|---|---|
| SM 18:1;O2/16:1 | 3.8 | 0.3 | 3.7 | 0.2 | 3.6 | 0.4 | 3.8 | 0.2 | 3.4 | 0.2 | 3.5 | 0.3 | 3.6 | 0.2 | 3.7 | 0.3 |
| SM 18:1;O2/18:0 | 12.0 | 1.2 | 10.9 | 0.7 | 10.5 | 0.9 | 10.4 | 0.4 | 10.1 | 1.3 | 9.9 | 1.2 | 11.5 | 0.9 | 9.9 | 1.0 |
| SM 18:1;O2/18:1 | 3.6 | 0.2 | 3.6 | 0.1 | 3.3 | 0.2 | 3.6 | 0.1 | 2.9 | 0.2 | 3.2 | 0.3 | 3.1 | 0.4 | 3.2 | 0.2 |
| SM 18:1;O2/18:2 | 0.0 | 0.0 | 0.0 | 0.0 | 0.0 | 0.0 | 0.0 | 0.0 | 0.0 | 0.0 | 0.0 | 0.0 | 0.0 | 0.0 | 0.0 | 0.0 |
| SM 18:1;O2/20:0 | 4.1 | 0.2 | 4.1 | 0.4 | 3.7 | 0.6 | 3.8 | 0.1 | 3.4 | 0.3 | 3.5 | 0.6 | 3.9 | 0.5 | 3.5 | 0.3 |
| SM 18:1;O2/20:1 | 2.3 | 0.1 | 2.4 | 0.2 | 2.1 | 0.2 | 2.5 | 0.1 | 1.9 | 0.3 | 2.2 | 0.3 | 2.0 | 0.3 | 2.1 | 0.1 |
| SM 18:1;O2/20:2 | 0.1 | 0.0 | 0.1 | 0.0 | 0.1 | 0.0 | 0.1 | 0.0 | 0.1 | 0.0 | 0.1 | 0.0 | 0.1 | 0.0 | 0.1 | 0.0 |
| SM 18:1;O2/22:0 | 5.7 | 0.4 | 5.7 | 0.5 | 5.1 | 0.7 | 5.6 | 0.4 | 4.7 | 0.3 | 5.0 | 0.7 | 5.4 | 0.6 | 5.1 | 0.3 |
| SM 18:1;O2/22:1 | 3.3 | 0.3 | 3.6 | 0.2 | 3.0 | 0.4 | 3.5 | 0.2 | 2.8 | 0.2 | 3.2 | 0.4 | 2.9 | 0.4 | 3.0 | 0.4 |
| SM 18:1;O2/22:2 | 0.7 | 0.1 | 0.7 | 0.0 | 0.6 | 0.1 | 0.6 | 0.1 | 0.6 | 0.0 | 0.6 | 0.0 | 0.6 | 0.1 | 0.6 | 0.1 |
| SM 18:1;O2/22:3 | 0.1 | 0.0 | 0.1 | 0.0 | 0.1 | 0.0 | 0.1 | 0.0 | 0.1 | 0.0 | 0.2 | 0.0 | 0.1 | 0.0 | 0.1 | 0.1 |
| SM 18:1;O2/22:5 | 0.0 | 0.0 | 0.0 | 0.0 | 0.0 | 0.0 | 0.0 | 0.0 | 0.0 | 0.0 | 0.0 | 0.0 | 0.0 | 0.0 | 0.0 | 0.0 |
| SM total | 98.0 | 5.0 | 93.2 | 1.2 | 88.2 | 5.5 | 89.4 | 3.4 | 82.7 | 8.2 | 83.2 | 7.1 | 94.0 | 8.4 | 86.2 | 7.8 |
| Cer 18:1;O2/16:0 | 4.4 | 0.8 | 3.2 | 0.7 | 3.1 | 0.8 | 2.5 | 0.2 | 2.4 | 0.6 | 2.1 | 0.6 | 3.0 | 0.5 | 1.9 | 0.5 |
| Cer 18:1;O2/18:0 | 5.4 | 1.6 | 2.6 | 0.6 | 4.1 | 1.6 | 2.8 | 1.1 | 2.9 | 1.5 | 2.4 | 0.8 | 4.2 | 0.9 | 2.2 | 0.3 |
| Cer 18:1;O2/18:1 | 0.0 | 0.0 | 0.0 | 0.0 | 0.0 | 0.0 | 0.0 | 0.0 | 0.0 | 0.0 | 0.0 | 0.0 | 0.0 | 0.0 | 0.0 | 0.0 |
| Cer 18:1;O2/18:2 | 0.3 | 0.1 | 0.2 | 0.1 | 0.2 | 0.1 | 0.1 | 0.1 | 0.1 | 0.0 | 0.1 | 0.1 | 0.1 | 0.1 | 0.1 | 0.0 |
| Cer 18:1;O2/20:0 | 6.2 | 1.2 | 3.6 | 0.7 | 4.6 | 1.1 | 3.2 | 0.3 | 3.4 | 1.1 | 2.9 | 0.9 | 4.4 | 0.9 | 2.5 | 0.6 |
| Cer 18:1;O2/20:1 | 0.0 | 0.0 | 0.0 | 0.0 | 0.0 | 0.0 | 0.0 | 0.0 | 0.0 | 0.0 | 0.0 | 0.0 | 0.0 | 0.0 | 0.0 | 0.0 |
| Cer 18:1;O2/20:2 | 0.5 | 0.2 | 0.3 | 0.2 | 0.4 | 0.1 | 0.3 | 0.0 | 0.2 | 0.1 | 0.2 | 0.1 | 0.3 | 0.1 | 0.2 | 0.1 |
| Cer 18:1;O2/20:4 | 0.0 | 0.0 | 0.0 | 0.0 | 0.0 | 0.0 | 0.0 | n.d. | 0.0 | 0.0 | 0.0 | n.d. | 0.0 | 0.0 | 0.0 | n.d. |
| Cer 18:1;O2/22:0 | 20.9 | 4.1 | 13.4 | 2.1 | 15.5 | 3.5 | 11.5 | 0.8 | 11.7 | 3.9 | 10.2 | 2.5 | 15.9 | 3.9 | 9.7 | 2.4 |
| Cer 18:1;O2/22:1 | 0.3 | 0.1 | 0.3 | 0.1 | 0.2 | 0.1 | 0.2 | 0.0 | 0.2 | 0.0 | 0.2 | 0.1 | 0.2 | 0.1 | 0.2 | 0.1 |
| Cer 18:1;O2/22:2 | 6.5 | 1.4 | 4.4 | 0.9 | 4.6 | 1.2 | 3.4 | 0.3 | 3.4 | 1.1 | 2.9 | 0.7 | 3.8 | 1.2 | 2.5 | 0.5 |
| Cer 18:1;O2/22:3 | 0.1 | 0.0 | 0.0 | 0.0 | 0.0 | 0.0 | 0.0 | 0.0 | 0.0 | 0.0 | 0.0 | 0.0 | 0.0 | 0.0 | 0.0 | 0.0 |

| | | | | | | | | | | | | | | | | |
|---|---|---|---|---|---|---|---|---|---|---|---|---|---|---|---|---|
| Cer 18:1;O2/22:4 | 0.0 | 0.0 | 0.0 | 0.0 | 0.0 | 0.0 | 0.0 | 0.0 | 0.0 | n.d. | 0.0 | 0.0 | 0.0 | 0.0 | 0.0 | 0.0 |
| Cer total | 44.6 | 8.8 | 27.9 | 4.9 | 32.8 | 8.0 | 24.1 | 2.2 | 24.3 | 8.1 | 21.2 | 5.6 | 31.9 | 6.9 | 19.4 | 4.2 |
| CE 14:0 | 38.0 | 6.1 | 37.9 | 3.3 | 26.6 | 4.0 | 27.4 | 3.2 | 21.7 | 3.7 | 21.3 | 2.2 | 27.2 | 5.0 | 23.0 | 3.0 |
| CE 14:1 | 7.8 | 2.6 | 11.9 | 3.0 | 5.8 | 2.1 | 9.0 | 1.1 | 5.0 | 1.8 | 7.7 | 0.5 | 6.1 | 2.6 | 8.3 | 1.7 |
| CE 16:0 | 2986.2 | 310.4 | 2903.5 | 214.3 | 2236.3 | 354.7 | 2111.9 | 129.5 | 1860.5 | 348.3 | 1756.1 | 133.0 | 2327.4 | 362.8 | 1846.5 | 278.2 |
| CE 16:1 | 1381.6 | 226.4 | 1608.4 | 284.7 | 1066.7 | 231.0 | 1410.8 | 154.0 | 1027.6 | 246.3 | 1410.1 | 115.1 | 1446.2 | 616.4 | 1518.8 | 112.6 |
| CE 18:0 | 742.9 | 73.2 | 734.3 | 68.6 | 586.1 | 79.2 | 566.1 | 25.1 | 562.7 | 104.8 | 536.2 | 78.2 | 747.3 | 211.0 | 607.6 | 105.5 |
| CE 18:1 | 9373.9 | 1360.0 | 8552.9 | 1082.1 | 7888.3 | 1043.3 | 7465.1 | 530.5 | 7830.3 | 1092.0 | 7582.7 | 902.7 | 9574.0 | 1966.4 | 8162.9 | 1168.0 |
| CE 18:2 | 51683.1 | 8527.4 | 50056.8 | 8442.9 | 41060.6 | 6499.3 | 39892.6 | 1764.3 | 36728.3 | 2857.5 | 36384.2 | 2865.8 | 43358.8 | 7713.4 | 35475.4 | 3337.3 |
| CE 18:3 | 7150.9 | 1172.1 | 7352.5 | 1852.9 | 4888.1 | 779.5 | 4862.8 | 662.0 | 3893.3 | 359.1 | 4122.1 | 421.2 | 5568.2 | 930.4 | 4579.5 | 413.3 |
| CE 18:4 | 33.6 | 7.1 | 33.9 | 9.5 | 20.8 | 6.9 | 20.0 | 4.0 | 15.4 | 4.6 | 14.8 | 2.0 | 48.4 | 57.9 | 28.1 | 16.9 |
| CE 20:0 | 13.0 | 2.3 | 14.1 | 1.6 | 10.2 | 2.2 | 9.6 | 1.0 | 9.0 | 1.4 | 7.9 | 0.9 | 10.1 | 1.0 | 7.9 | 0.4 |
| CE 20:1 | 32.9 | 11.3 | 30.6 | 7.8 | 25.1 | 5.5 | 24.5 | 3.6 | 26.6 | 8.5 | 22.5 | 4.8 | 32.0 | 8.4 | 27.8 | 10.1 |
| CE 20:2 | 74.4 | 20.0 | 63.4 | 19.8 | 53.4 | 7.5 | 50.4 | 7.0 | 50.1 | 9.0 | 41.3 | 13.6 | 76.9 | 26.5 | 59.0 | 19.4 |
| CE 20:3 | 367.7 | 85.2 | 365.1 | 65.7 | 241.7 | 51.3 | 240.4 | 6.6 | 181.8 | 33.1 | 178.2 | 33.2 | 328.0 | 137.4 | 242.8 | 77.4 |
| CE 20:4 | 3856.8 | 985.3 | 3951.0 | 631.2 | 2659.7 | 981.4 | 2535.1 | 307.5 | 1848.7 | 419.7 | 2010.7 | 322.8 | 3308.0 | 1323.4 | 2413.0 | 539.9 |
| CE 20:5 | 165.7 | 41.0 | 192.2 | 36.8 | 94.7 | 20.6 | 97.8 | 15.6 | 66.1 | 5.0 | 68.5 | 7.2 | 135.5 | 79.8 | 84.5 | 27.6 |
| CE 22:0 | 7.2 | 1.4 | 8.0 | 0.9 | 5.2 | 0.9 | 5.1 | 0.6 | 4.5 | 0.6 | 4.3 | 0.4 | 4.7 | 0.9 | 4.1 | 0.7 |
| CE 22:1 | 6.3 | 2.1 | 6.2 | 1.6 | 4.0 | 0.5 | 3.9 | 0.4 | 3.7 | 0.6 | 3.3 | 0.3 | 5.3 | 1.5 | 3.8 | 1.0 |
| CE 22:2 | 3.5 | 0.9 | 3.7 | 1.0 | 2.3 | 0.3 | 2.1 | 0.3 | 2.0 | 0.3 | 1.8 | 0.5 | 3.3 | 1.1 | 2.5 | 0.7 |
| CE 22:3 | 2.9 | 0.5 | 3.0 | 0.7 | 1.9 | 0.3 | 1.7 | 0.1 | 1.5 | 0.4 | 1.3 | 0.3 | 2.8 | 1.4 | 2.1 | 0.6 |
| CE 22:4 | 15.5 | 3.9 | 17.2 | 2.9 | 10.8 | 3.5 | 10.3 | 1.5 | 8.6 | 2.1 | 8.7 | 1.2 | 16.3 | 7.9 | 11.8 | 2.9 |
| CE 22:5 | 39.8 | 10.8 | 44.3 | 7.4 | 27.7 | 10.4 | 27.0 | 4.3 | 20.2 | 4.3 | 22.3 | 3.3 | 44.1 | 25.1 | 32.2 | 9.6 |
| CE 22:6 | 46.1 | 12.2 | 44.4 | 14.4 | 28.3 | 9.9 | 27.7 | 6.4 | 22.5 | 3.0 | 25.6 | 5.0 | 45.3 | 20.7 | 34.4 | 8.4 |
| CE total | 78029.8 | 11928.6 | 76035.2 | 12278.1 | 60944.4 | 8484.6 | 59401.3 | 2956.0 | 54190.0 | 2960.1 | 54231.6 | 3461.8 | 67115.9 | 11902.4 | 55176.2 | 3994.2 |

| | | | | | | | | | | | | | | | | |
|---|---|---|---|---|---|---|---|---|---|---|---|---|---|---|---|---|
| MG 16:0 | 8.1 | 3.7 | 5.2 | 0.6 | 9.6 | 3.4 | 10.4 | 7.8 | 8.0 | 4.7 | 6.8 | 2.5 | 7.4 | 2.8 | 6.0 | 1.2 |
| MG 18:0 | 19.4 | 4.4 | 14.1 | 1.4 | 47.6 | 24.6 | 37.4 | 15.3 | 31.3 | 8.2 | 34.3 | 20.4 | 29.3 | 10.4 | 28.2 | 8.0 |
| MG 18:1 | 3.4 | 0.7 | 3.1 | 0.6 | 2.5 | 0.8 | 2.4 | 0.3 | 6.2 | 4.2 | 4.1 | 1.9 | 5.1 | 1.7 | 3.7 | 1.3 |
| MG 18:2 | 4.5 | 2.0 | 2.0 | 1.0 | 6.0 | 4.7 | 6.2 | 3.1 | 2.0 | 0.7 | 2.4 | 0.9 | 1.6 | 0.8 | 1.5 | 0.3 |
| MG total | 35.4 | 7.9 | 24.5 | 1.9 | 65.7 | 25.0 | 56.3 | 15.8 | 47.5 | 12.1 | 47.6 | 25.1 | 43.3 | 13.1 | 39.3 | 10.3 |
| DG 14:0_14:0 | 0.0 | 0.0 | 0.0 | 0.0 | 0.0 | 0.0 | 0.0 | 0.0 | 0.0 | 0.0 | 0.0 | n.d. | 0.0 | n.d. | 0.0 | 0.0 |
| DG 14:0_16:0 | 0.1 | 0.1 | 0.1 | 0.0 | 0.1 | 0.0 | 0.1 | 0.1 | 0.1 | 0.0 | 0.1 | 0.0 | 0.1 | 0.0 | 0.0 | 0.0 |
| DG 14:0_16:1 | 0.0 | 0.0 | 0.0 | 0.0 | 0.0 | 0.0 | 0.0 | 0.0 | 0.0 | 0.0 | 0.0 | 0.0 | 0.0 | 0.0 | 0.0 | 0.0 |
| DG 14:0_18:0 | 0.0 | 0.0 | 0.0 | 0.0 | 0.0 | 0.0 | 0.0 | 0.0 | 0.0 | 0.0 | 0.0 | 0.0 | 0.0 | 0.0 | 0.0 | 0.0 |
| DG 14:0_18:1 | 0.1 | 0.1 | 0.1 | 0.1 | 0.1 | 0.1 | 0.1 | 0.0 | 0.2 | 0.1 | 0.1 | 0.0 | 0.1 | 0.0 | 0.1 | 0.0 |
| DG 14:0_18:2 | 0.1 | 0.0 | 0.1 | 0.0 | 0.1 | 0.0 | 0.1 | 0.0 | 0.1 | 0.0 | 0.1 | 0.0 | 0.1 | 0.0 | 0.1 | 0.0 |
| DG 14:0_18:3 | 0.0 | 0.0 | 0.0 | 0.0 | 0.0 | 0.0 | 0.0 | 0.0 | 0.0 | 0.0 | 0.0 | 0.0 | 0.0 | 0.0 | 0.0 | 0.0 |
| DG 14:0_20:0 | 0.0 | 0.0 | 0.0 | 0.0 | 0.0 | 0.0 | 0.0 | 0.0 | 0.0 | 0.0 | 0.0 | 0.0 | 0.0 | 0.0 | 0.0 | 0.0 |
| DG 14:1_18:1 | 0.0 | 0.0 | 0.0 | 0.0 | 0.0 | 0.0 | 0.0 | 0.0 | 0.0 | 0.0 | 0.0 | 0.0 | 0.0 | 0.0 | 0.0 | 0.0 |
| DG 14:1_18:2 | 0.0 | 0.0 | 0.0 | 0.0 | 0.0 | 0.0 | 0.0 | 0.0 | 0.0 | 0.0 | 0.0 | 0.0 | 0.0 | 0.0 | 0.0 | 0.0 |
| DG 14:1_22:2 | 0.0 | 0.0 | 0.0 | 0.0 | 0.0 | 0.0 | 0.0 | 0.0 | 0.0 | 0.0 | 0.0 | n.d. | 0.0 | 0.0 | 0.0 | 0.0 |
| DG 16:0_16:0 | 0.4 | 0.1 | 0.3 | 0.1 | 0.3 | 0.1 | 0.3 | 0.1 | 0.3 | 0.1 | 0.2 | 0.1 | 0.4 | 0.1 | 0.2 | 0.0 |
| DG 16:0_16:1 | 0.2 | 0.1 | 0.2 | 0.1 | 0.1 | 0.1 | 0.2 | 0.0 | 0.2 | 0.1 | 0.2 | 0.1 | 0.3 | 0.1 | 0.2 | 0.0 |
| DG 16:0_18:0 | 0.2 | 0.1 | 0.1 | 0.0 | 0.1 | 0.1 | 0.1 | 0.0 | 0.1 | 0.1 | 0.1 | 0.0 | 0.2 | 0.1 | 0.1 | 0.0 |
| DG 16:0_18:1 | 1.9 | 0.5 | 1.5 | 0.5 | 1.3 | 0.5 | 1.0 | 0.1 | 1.6 | 0.4 | 1.1 | 0.2 | 2.1 | 0.6 | 1.1 | 0.2 |
| DG 16:0_18:2 | 2.0 | 0.3 | 1.6 | 0.5 | 1.5 | 0.2 | 1.3 | 0.1 | 1.7 | 0.1 | 1.4 | 0.1 | 2.3 | 1.0 | 1.5 | 0.2 |
| DG 16:0_18:3 | 0.3 | 0.1 | 0.2 | 0.1 | 0.2 | 0.0 | 0.2 | 0.0 | 0.2 | 0.0 | 0.2 | 0.1 | 0.3 | 0.1 | 0.2 | 0.0 |
| DG 16:0_20:2 | 0.0 | 0.0 | 0.0 | 0.0 | 0.0 | 0.0 | 0.0 | 0.0 | 0.0 | 0.0 | 0.0 | 0.0 | 0.0 | 0.0 | 0.0 | 0.0 |
| DG 16:0_20:3 | 0.0 | 0.0 | 0.0 | 0.0 | 0.0 | 0.0 | 0.0 | 0.0 | 0.0 | 0.0 | 0.0 | 0.0 | 0.0 | 0.0 | 0.0 | 0.0 |
| DG 16:0_20:4 | 0.0 | 0.0 | 0.0 | 0.0 | 0.0 | 0.0 | 0.0 | 0.0 | 0.0 | 0.0 | 0.0 | 0.0 | 0.0 | 0.0 | 0.0 | 0.0 |

| | | | | | | | | | | | | | | | | |
|---|---|---|---|---|---|---|---|---|---|---|---|---|---|---|---|---|
| DG 16:0_22:4 | 0.0 | 0.0 | 0.0 | 0.0 | 0.0 | 0.0 | 0.0 | 0.0 | 0.0 | 0.0 | 0.0 | 0.0 | 0.0 | 0.0 | 0.0 | 0.0 |
| DG 16:0_22:5 | 0.0 | 0.0 | 0.0 | 0.0 | 0.0 | 0.0 | 0.0 | 0.0 | 0.0 | 0.0 | 0.0 | 0.0 | 0.0 | 0.0 | 0.0 | 0.0 |
| DG 16:1_16:1 | 0.0 | 0.0 | 0.0 | 0.0 | 0.0 | 0.0 | 0.0 | 0.0 | 0.0 | 0.0 | 0.0 | 0.0 | 0.0 | 0.0 | 0.0 | 0.0 |
| DG 16:1_18:0 | 0.0 | 0.0 | 0.0 | 0.0 | 0.0 | 0.0 | 0.0 | 0.0 | 0.0 | 0.0 | 0.0 | 0.0 | 0.0 | 0.0 | 0.0 | 0.0 |
| DG 16:1_18:1 | 0.3 | 0.1 | 0.3 | 0.1 | 0.2 | 0.1 | 0.3 | 0.0 | 0.3 | 0.1 | 0.3 | 0.1 | 0.4 | 0.1 | 0.3 | 0.1 |
| DG 16:1_18:2 | 0.4 | 0.1 | 0.4 | 0.1 | 0.3 | 0.1 | 0.4 | 0.0 | 0.4 | 0.1 | 0.4 | 0.0 | 0.5 | 0.2 | 0.4 | 0.1 |
| DG 16:1_18:3 | 0.0 | 0.0 | 0.1 | 0.0 | 0.0 | 0.0 | 0.1 | 0.0 | 0.0 | 0.0 | 0.0 | 0.0 | 0.0 | 0.0 | 0.0 | 0.0 |
| DG 16:1_20:2 | 0.0 | 0.0 | 0.0 | 0.0 | 0.0 | 0.0 | 0.0 | 0.0 | 0.0 | 0.0 | 0.0 | 0.0 | 0.0 | 0.0 | 0.0 | 0.0 |
| DG 16:1_20:3 | 0.0 | 0.0 | 0.0 | 0.0 | 0.0 | 0.0 | 0.0 | 0.0 | 0.0 | 0.0 | 0.0 | 0.0 | 0.0 | 0.0 | 0.0 | 0.0 |
| DG 16:1_20:4 | 0.0 | 0.0 | 0.0 | 0.0 | 0.0 | 0.0 | 0.0 | 0.0 | 0.0 | 0.0 | 0.0 | 0.0 | 0.0 | 0.0 | 0.0 | 0.0 |
| DG 18:0_18:0 | 0.0 | 0.0 | 0.0 | 0.0 | 0.0 | 0.0 | 0.0 | 0.0 | 0.0 | 0.0 | 0.0 | 0.0 | 0.0 | 0.0 | 0.0 | 0.0 |
| DG 18:0_18:1 | 0.3 | 0.1 | 0.3 | 0.1 | 0.2 | 0.1 | 0.2 | 0.0 | 0.3 | 0.1 | 0.2 | 0.1 | 0.4 | 0.1 | 0.2 | 0.1 |
| DG 18:0_18:2 | 0.6 | 0.1 | 0.5 | 0.2 | 0.4 | 0.1 | 0.4 | 0.1 | 0.5 | 0.1 | 0.4 | 0.1 | 0.6 | 0.2 | 0.4 | 0.1 |
| DG 18:0_18:3 | 0.1 | 0.0 | 0.0 | 0.0 | 0.0 | 0.0 | 0.0 | 0.0 | 0.1 | 0.0 | 0.1 | 0.0 | 0.1 | 0.0 | 0.0 | 0.0 |
| DG 18:0_20:4 | 0.0 | 0.0 | 0.0 | 0.0 | 0.0 | 0.0 | 0.0 | 0.0 | 0.0 | 0.0 | 0.0 | 0.0 | 0.0 | 0.0 | 0.0 | 0.0 |
| DG 18:1_18:1 | 1.7 | 0.6 | 1.3 | 0.4 | 1.2 | 0.6 | 0.9 | 0.1 | 1.4 | 0.4 | 1.0 | 0.2 | 1.6 | 0.3 | 1.0 | 0.2 |
| DG 18:1_18:2 | 3.9 | 0.6 | 3.2 | 0.9 | 2.8 | 0.8 | 2.7 | 0.3 | 3.4 | 0.3 | 2.9 | 0.5 | 4.2 | 1.2 | 3.0 | 0.6 |
| DG 18:1_18:3 | 0.5 | 0.1 | 0.5 | 0.1 | 0.3 | 0.1 | 0.3 | 0.1 | 0.4 | 0.1 | 0.4 | 0.0 | 0.5 | 0.1 | 0.4 | 0.1 |
| DG 18:1_18:4 | 0.0 | 0.0 | 0.0 | 0.0 | 0.0 | n.d. | 0.0 | n.d. | 0.0 | 0.0 | 0.0 | 0.0 | 0.0 | 0.0 | 0.0 | 0.0 |
| DG 18:1_20:2 | 0.0 | 0.0 | 0.0 | 0.0 | 0.0 | 0.0 | 0.0 | 0.0 | 0.0 | 0.0 | 0.0 | 0.0 | 0.0 | 0.0 | 0.0 | 0.0 |
| DG 18:1_20:3 | 0.0 | 0.0 | 0.0 | 0.0 | 0.0 | 0.0 | 0.0 | 0.0 | 0.0 | 0.0 | 0.0 | 0.0 | 0.0 | 0.0 | 0.0 | 0.0 |
| DG 18:1_20:4 | 0.1 | 0.0 | 0.1 | 0.0 | 0.0 | 0.0 | 0.0 | 0.0 | 0.0 | 0.0 | 0.0 | 0.0 | 0.1 | 0.1 | 0.1 | 0.0 |
| DG 18:1_22:5 | 0.0 | 0.0 | 0.0 | 0.0 | 0.0 | 0.0 | 0.0 | 0.0 | 0.0 | 0.0 | 0.0 | n.d. | 0.0 | 0.0 | 0.0 | 0.0 |
| DG 18:2_18:2 | 1.3 | 0.1 | 1.2 | 0.3 | 1.0 | 0.1 | 1.1 | 0.2 | 1.2 | 0.2 | 1.1 | 0.1 | 1.6 | 0.7 | 1.1 | 0.3 |
| DG 18:2_18:3 | 0.2 | 0.0 | 0.2 | 0.1 | 0.1 | 0.1 | 0.2 | 0.0 | 0.2 | 0.0 | 0.2 | 0.0 | 0.2 | 0.1 | 0.2 | 0.0 |

| | | | | | | | | | | | | | | | | |
|---|---|---|---|---|---|---|---|---|---|---|---|---|---|---|---|---|
| DG 18:2_20:2 | 0.0 | 0.0 | 0.0 | 0.0 | 0.0 | 0.0 | 0.0 | 0.0 | 0.0 | 0.0 | 0.0 | 0.0 | 0.0 | 0.0 | 0.0 | 0.0 |
| DG 18:2_20:3 | 0.0 | 0.0 | 0.0 | 0.0 | 0.0 | 0.0 | 0.0 | 0.0 | 0.0 | 0.0 | 0.0 | 0.0 | 0.0 | 0.0 | 0.0 | 0.0 |
| DG 18:2_20:4 | 0.0 | 0.0 | 0.0 | 0.0 | 0.0 | 0.0 | 0.0 | 0.0 | 0.0 | 0.0 | 0.0 | 0.0 | 0.1 | 0.1 | 0.0 | 0.0 |
| DG 18:3_18:3 | 0.0 | 0.0 | 0.0 | 0.0 | 0.0 | 0.0 | 0.0 | 0.0 | 0.0 | 0.0 | 0.0 | 0.0 | 0.0 | 0.0 | 0.0 | 0.0 |
| DG total | 15.3 | 2.5 | 12.6 | 3.4 | 10.8 | 3.1 | 10.1 | 1.0 | 13.1 | 1.7 | 10.7 | 1.5 | 16.6 | 5.1 | 11.1 | 1.9 |

Lipidomics data were normalized using the ion abundance of internal standards added before lipid extraction via the modified Bligh and Dyer method.

n.d.: not detected.

### Supplementary Table 8 Quantitative lipidomics for erythrocytes.

| Lipids | Quantification (amol cell$^{-1}$) | | | | | | | | | | | | | | | |
|---|---|---|---|---|---|---|---|---|---|---|---|---|---|---|---|---|
| | 8 months | | | | 12 months | | | | 16 months | | | | 20 months | | | |
| | Severe | | Mild | | Severe | | Mild | | Severe | | Mild | | Severe | | Mild | |
| | Mean | SD | Mean | SD | Mean | SD | Mean | SD | Mean | SD | Mean | SD | Mean | SD | Mean | SD |
| LPC 16:0 | 15.66 | 2.70 | 16.15 | 1.75 | 14.56 | 2.36 | 14.78 | 2.22 | 14.58 | 1.17 | 16.13 | 1.25 | 15.24 | 1.82 | 14.63 | 1.18 |
| LPC 16:1 | 0.08 | 0.03 | 0.11 | 0.02 | 0.06 | 0.03 | 0.09 | 0.02 | 0.07 | 0.04 | 0.10 | 0.01 | 0.07 | 0.03 | 0.11 | 0.02 |
| LPC 18:0 | 26.63 | 5.39 | 26.27 | 2.52 | 24.68 | 4.01 | 27.44 | 1.39 | 24.83 | 4.68 | 29.48 | 3.27 | 22.95 | 5.74 | 23.57 | 2.06 |
| LPC 18:1 | 3.78 | 0.80 | 4.01 | 0.31 | 3.18 | 0.80 | 3.40 | 0.56 | 3.33 | 0.61 | 3.69 | 0.35 | 3.43 | 0.70 | 3.35 | 0.18 |
| LPC 18:2 | 3.29 | 0.45 | 3.31 | 0.36 | 3.97 | 0.85 | 3.94 | 0.66 | 2.74 | 0.45 | 2.98 | 0.29 | 2.81 | 0.99 | 2.76 | 0.47 |
| LPC 18:3 | 0.02 | 0.01 | 0.02 | 0.01 | 0.05 | 0.03 | 0.05 | 0.03 | 0.02 | 0.01 | 0.02 | 0.00 | 0.02 | 0.01 | 0.03 | 0.02 |
| LPC 20:0 | 0.11 | 0.03 | 0.13 | 0.04 | 0.08 | 0.01 | 0.12 | 0.03 | 0.10 | 0.03 | 0.12 | 0.03 | 0.08 | 0.03 | 0.07 | 0.03 |
| LPC 20:1 | 0.07 | 0.02 | 0.07 | 0.03 | 0.08 | 0.02 | 0.08 | 0.03 | 0.07 | 0.03 | 0.07 | 0.02 | 0.08 | 0.04 | 0.07 | 0.03 |
| LPC 20:2 | 0.07 | 0.02 | 0.07 | 0.04 | 0.12 | 0.05 | 0.12 | 0.04 | 0.06 | 0.02 | 0.05 | 0.02 | 0.06 | 0.02 | 0.04 | 0.03 |
| LPC 20:3 | 0.03 | 0.01 | 0.05 | 0.01 | 0.07 | 0.03 | 0.05 | 0.04 | 0.02 | 0.00 | 0.03 | 0.01 | 0.04 | 0.02 | 0.04 | 0.01 |
| LPC 20:4 | 0.12 | 0.04 | 0.14 | 0.02 | 0.15 | 0.03 | 0.18 | 0.05 | 0.08 | 0.03 | 0.12 | 0.02 | 0.13 | 0.07 | 0.14 | 0.02 |
| LPC total | 49.86 | 9.16 | 50.32 | 4.92 | 47.00 | 7.55 | 50.24 | 4.62 | 45.90 | 5.99 | 52.79 | 4.32 | 44.92 | 7.62 | 44.82 | 1.30 |
| LPE 16:0 | 0.26 | 0.10 | 0.29 | 0.16 | 0.16 | 0.04 | 0.26 | 0.10 | 0.42 | 0.10 | 0.28 | 0.10 | 0.35 | 0.20 | 0.55 | 0.25 |
| LPE 18:0 | 0.69 | 0.25 | 0.82 | 0.25 | 0.59 | 0.19 | 0.55 | 0.18 | 0.64 | 0.16 | 0.67 | 0.15 | 0.74 | 0.20 | 0.58 | 0.10 |
| LPE 18:1 | 0.64 | 0.21 | 0.63 | 0.21 | 0.20 | 0.12 | 0.25 | 0.12 | 0.55 | 0.19 | 0.64 | 0.14 | 0.66 | 0.15 | 0.53 | 0.09 |
| LPE 18:2 | 1.89 | 0.62 | 2.40 | 0.37 | 1.31 | 0.89 | 1.50 | 0.84 | 1.65 | 0.52 | 1.86 | 0.36 | 1.77 | 0.66 | 1.31 | 0.59 |
| LPE total | 3.47 | 0.92 | 4.15 | 0.78 | 2.26 | 1.06 | 2.57 | 0.75 | 3.26 | 0.45 | 3.44 | 0.53 | 3.52 | 0.70 | 2.98 | 0.29 |
| PC 16:0_16:0 | 52.07 | 7.03 | 56.69 | 6.66 | 43.83 | 4.87 | 55.33 | 2.64 | 48.79 | 6.73 | 53.80 | 7.18 | 44.56 | 7.11 | 50.92 | 0.97 |
| PC 16:0_16:1 | 1.92 | 0.39 | 2.67 | 0.25 | 1.78 | 0.82 | 2.47 | 0.38 | 1.84 | 0.71 | 3.00 | 0.33 | 2.23 | 0.87 | 2.20 | 1.00 |
| PC 16:0_18:0 | 33.03 | 3.73 | 29.92 | 4.96 | 33.08 | 6.05 | 34.80 | 3.32 | 29.83 | 4.99 | 30.68 | 2.67 | 29.62 | 4.50 | 29.02 | 2.17 |

| | | | | | | | | | | | | | | | | |
|---|---|---|---|---|---|---|---|---|---|---|---|---|---|---|---|---|
| PC 16:0_18:1 | 90.81 | 12.93 | 102.90 | 14.22 | 98.70 | 16.05 | 103.27 | 5.61 | 80.02 | 9.36 | 89.40 | 12.57 | 88.44 | 17.06 | 81.71 | 11.00 |
| PC 16:0_18:2 | 105.01 | 10.85 | 130.41 | 38.73 | 159.72 | 20.18 | 165.07 | 34.88 | 84.46 | 14.46 | 100.51 | 15.64 | 112.11 | 39.77 | 90.72 | 51.38 |
| PC 16:0_18:3 | 1.20 | 0.46 | 2.06 | 0.68 | 4.12 | 1.91 | 4.31 | 2.32 | 0.78 | 0.45 | 1.09 | 0.45 | 1.40 | 0.74 | 1.43 | 0.68 |
| PC 16:0_20:1 | 2.22 | 0.54 | 1.36 | 0.59 | 2.60 | 0.99 | 2.37 | 1.25 | 1.60 | 0.85 | 1.54 | 0.21 | 1.79 | 0.69 | 1.24 | 0.39 |
| PC 16:0_20:2 | 1.88 | 0.49 | 2.50 | 1.49 | 3.44 | 0.87 | 3.67 | 1.35 | 1.69 | 0.77 | 1.83 | 1.26 | 2.13 | 1.58 | 1.91 | 1.40 |
| PC 16:0_20:3 | 1.15 | 0.50 | 1.36 | 0.92 | 3.07 | 0.85 | 3.33 | 1.20 | 0.57 | 0.28 | 0.77 | 0.52 | 1.57 | 0.73 | 1.01 | 0.63 |
| PC 16:0_20:4 | 1.79 | 0.38 | 2.72 | 0.87 | 4.47 | 1.22 | 6.02 | 2.97 | 1.29 | 0.40 | 1.79 | 0.75 | 2.59 | 1.71 | 1.85 | 1.02 |
| PC 16:1_18:2 | 0.60 | 0.50 | 0.68 | 0.54 | 1.03 | 0.44 | 1.94 | 0.88 | 0.67 | 0.52 | 0.76 | 0.20 | 0.92 | 0.68 | 0.88 | 0.82 |
| PC 18:0_18:0 | 4.04 | 0.90 | 3.30 | 0.51 | 4.03 | 0.67 | 4.31 | 0.51 | 3.17 | 1.36 | 3.67 | 0.60 | 3.43 | 1.10 | 3.71 | 0.83 |
| PC 18:0_18:1 | 31.05 | 4.26 | 29.90 | 3.28 | 41.07 | 7.06 | 44.47 | 4.02 | 29.57 | 5.28 | 30.07 | 5.23 | 32.38 | 10.89 | 28.88 | 7.50 |
| PC 18:0_18:2 | 71.72 | 8.65 | 84.35 | 17.58 | 130.80 | 18.37 | 139.05 | 24.27 | 65.87 | 14.80 | 80.28 | 14.34 | 85.36 | 37.18 | 70.63 | 40.77 |
| PC 18:0_18:3 | 0.98 | 0.33 | 1.05 | 0.42 | 3.30 | 1.23 | 2.98 | 1.52 | 0.42 | 0.27 | 0.73 | 0.51 | 1.17 | 0.84 | 1.79 | 0.62 |
| PC 18:0_20:2 | 0.55 | 0.31 | 0.23 | 0.17 | 0.70 | 0.44 | 0.92 | 0.10 | 0.13 | 0.12 | 0.48 | 0.27 | 0.48 | 0.16 | 0.72 | 0.81 |
| PC 18:0_20:3 | 0.59 | 0.18 | 0.89 | 0.27 | 2.34 | 1.08 | 2.85 | 0.77 | 0.34 | 0.15 | 0.78 | 0.47 | 0.74 | 0.34 | 0.82 | 0.42 |
| PC 18:0_20:4 | 1.81 | 0.55 | 2.66 | 0.42 | 4.80 | 1.09 | 6.27 | 2.82 | 1.02 | 0.47 | 1.92 | 0.87 | 1.97 | 1.32 | 1.91 | 1.29 |
| PC 18:1_18:1 | 5.57 | 1.24 | 4.99 | 1.75 | 7.61 | 1.92 | 8.39 | 1.26 | 4.84 | 1.39 | 5.48 | 0.45 | 5.93 | 2.19 | 4.81 | 1.95 |
| PC 18:1_18:2 | 11.80 | 2.46 | 14.69 | 5.15 | 27.46 | 6.67 | 26.55 | 7.01 | 8.74 | 2.39 | 11.39 | 2.19 | 14.57 | 8.09 | 10.14 | 6.74 |
| PC 18:2_18:2 | 5.99 | 1.43 | 6.73 | 2.57 | 17.51 | 4.46 | 16.15 | 6.83 | 4.33 | 1.21 | 5.22 | 0.90 | 6.52 | 4.29 | 4.18 | 3.31 |
| PC total | 425.78 | 42.96 | 482.07 | 96.71 | 595.47 | 81.83 | 634.51 | 81.93 | 369.96 | 56.40 | 425.18 | 57.75 | 439.79 | 134.06 | 388.69 | 130.46 |
| PC O-16:1/18:1 (PC P-16:0/18:1) | 2.44 | 0.38 | 3.35 | 1.36 | 3.34 | 0.71 | 5.18 | 0.70 | 3.07 | 1.49 | 3.02 | 1.24 | 2.97 | 1.01 | 3.19 | 2.06 |
| PC O-16:1/18:2 (PC P-16:0/18:2) | 1.21 | 0.35 | 1.98 | 0.41 | 2.52 | 0.94 | 3.49 | 1.24 | 0.87 | 0.41 | 1.08 | 0.74 | 1.48 | 1.05 | 1.32 | 1.09 |
| PC P-16:1/20:5 | 3.61 | 1.24 | 3.32 | 1.25 | 3.54 | 1.13 | 3.52 | 0.54 | 3.57 | 0.72 | 3.65 | 0.32 | 3.58 | 1.51 | 3.63 | 0.66 |
| PC O-18:0/18:1 | 0.82 | 0.28 | 0.81 | 0.57 | 0.69 | 0.34 | 1.15 | 0.21 | 0.63 | 0.37 | 0.67 | 0.30 | 0.70 | 0.13 | 0.52 | 0.21 |
| PC O-18:0/18:2 | 1.65 | 0.42 | 1.43 | 0.34 | 3.46 | 0.81 | 3.82 | 1.52 | 1.51 | 0.99 | 2.00 | 0.51 | 2.69 | 1.51 | 2.06 | 1.52 |
| PC P-18:2/20:5 | 10.16 | 1.53 | 9.32 | 2.15 | 11.46 | 2.30 | 11.44 | 2.45 | 9.99 | 1.65 | 10.07 | 1.44 | 11.21 | 3.65 | 9.28 | 2.14 |

| | | | | | | | | | | | | | | | | |
|---|---|---|---|---|---|---|---|---|---|---|---|---|---|---|---|---|
| EtherPC total | 19.89 | 2.60 | 20.20 | 4.63 | 25.02 | 2.82 | 28.59 | 5.15 | 19.64 | 3.47 | 20.49 | 3.77 | 22.62 | 8.02 | 20.00 | 6.16 |
| PE 16:0_16:0 | 3.25 | 0.54 | 2.95 | 0.22 | 2.68 | 0.73 | 3.49 | 0.87 | 2.63 | 0.67 | 2.22 | 0.70 | 3.25 | 0.76 | 2.72 | 0.45 |
| PE 16:0_16:1 | 1.39 | 0.33 | 1.93 | 0.59 | 1.34 | 0.51 | 1.70 | 0.23 | 1.44 | 0.34 | 2.15 | 0.32 | 1.68 | 0.94 | 1.63 | 0.39 |
| PE 16:0_18:0 | 3.68 | 0.18 | 3.69 | 0.41 | 3.90 | 1.03 | 4.11 | 0.87 | 2.81 | 0.29 | 3.38 | 0.34 | 3.16 | 0.41 | 2.65 | 0.29 |
| PE 16:0_18:1 | 47.77 | 7.15 | 53.56 | 5.17 | 54.19 | 10.53 | 65.72 | 8.86 | 44.13 | 3.85 | 48.72 | 3.18 | 47.34 | 14.13 | 44.33 | 9.52 |
| PE 16:0_18:2 | 90.85 | 13.41 | 100.88 | 18.47 | 191.39 | 40.28 | 201.82 | 53.42 | 65.99 | 12.58 | 80.30 | 13.00 | 109.12 | 53.72 | 74.30 | 52.11 |
| PE 16:0_18:3 | 1.14 | 0.51 | 1.73 | 1.34 | 5.57 | 2.64 | 6.08 | 3.97 | 0.27 | 0.12 | 0.68 | 0.15 | 1.08 | 0.91 | 1.24 | 1.42 |
| PE 16:0_20:1 | 0.35 | 0.02 | 0.29 | 0.22 | 0.73 | 0.38 | 0.53 | 0.21 | 0.33 | 0.20 | 0.20 | 0.09 | 0.26 | 0.17 | 0.42 | 0.28 |
| PE 16:0_20:3 | 0.65 | 0.41 | 0.94 | 0.45 | 2.82 | 1.36 | 3.42 | 1.18 | 0.35 | 0.16 | 0.61 | 0.42 | 1.37 | 0.95 | 1.06 | 0.49 |
| PE 16:0_20:4 | 4.80 | 1.34 | 6.75 | 1.57 | 22.09 | 7.15 | 24.60 | 11.99 | 2.68 | 0.60 | 3.73 | 2.08 | 7.37 | 3.96 | 4.02 | 3.64 |
| PE 16:0_22:4 | 0.87 | 0.37 | 1.53 | 0.66 | 5.53 | 1.29 | 6.39 | 4.18 | 0.35 | 0.26 | 0.79 | 0.64 | 1.22 | 0.42 | 0.75 | 0.77 |
| PE 16:1_18:0 | 0.47 | 0.32 | 0.41 | 0.15 | 0.39 | 0.18 | 0.63 | 0.55 | 0.49 | 0.18 | 0.53 | 0.24 | 0.61 | 0.08 | 0.47 | 0.29 |
| PE 16:1_18:1 | 1.21 | 0.32 | 1.49 | 0.50 | 0.86 | 0.31 | 1.71 | 0.65 | 1.02 | 0.50 | 1.65 | 0.18 | 1.21 | 0.57 | 1.48 | 1.18 |
| PE 16:1_18:2 | 1.30 | 0.80 | 1.69 | 0.84 | 2.12 | 1.24 | 3.89 | 2.66 | 0.44 | 0.18 | 0.44 | 0.22 | 1.64 | 0.80 | 1.15 | 0.99 |
| PE 16:1_20:4 | 0.39 | 0.11 | 0.46 | 0.23 | 0.28 | 0.22 | 0.54 | 0.31 | 0.36 | 0.10 | 0.24 | 0.11 | 0.40 | 0.11 | 0.24 | 0.09 |
| PE 18:0_18:0 | 0.32 | 0.19 | 0.56 | 0.17 | 0.26 | 0.12 | 0.48 | 0.18 | 0.27 | 0.24 | 0.41 | 0.16 | 0.52 | 0.29 | 0.30 | 0.16 |
| PE 18:0_18:1 | 13.70 | 2.08 | 15.27 | 2.56 | 15.82 | 3.45 | 18.81 | 4.04 | 12.12 | 1.54 | 13.82 | 3.21 | 12.37 | 4.47 | 10.83 | 3.97 |
| PE 18:0_18:2 | 58.06 | 10.00 | 64.48 | 12.44 | 103.77 | 21.42 | 114.05 | 27.51 | 44.63 | 11.93 | 47.10 | 13.67 | 66.32 | 31.90 | 48.49 | 29.53 |
| PE 18:0_20:3 | 0.86 | 0.27 | 0.78 | 0.42 | 1.92 | 0.61 | 2.28 | 1.59 | 0.55 | 0.27 | 0.67 | 0.18 | 0.84 | 0.73 | 0.92 | 0.54 |
| PE 18:0_20:4 | 7.14 | 1.40 | 7.63 | 1.37 | 19.64 | 5.10 | 19.69 | 10.09 | 4.49 | 1.36 | 5.43 | 1.16 | 8.20 | 3.94 | 5.67 | 1.77 |
| PE 18:0_22:4 | 0.39 | 0.11 | 0.57 | 0.19 | 1.98 | 0.70 | 2.18 | 1.27 | 0.17 | 0.09 | 0.27 | 0.23 | 0.56 | 0.20 | 0.43 | 0.24 |
| PE 18:1_18:1 | 14.55 | 1.13 | 15.95 | 4.47 | 18.75 | 2.90 | 22.62 | 3.41 | 12.64 | 1.41 | 13.70 | 2.43 | 15.14 | 5.54 | 14.18 | 6.58 |
| PE 18:1_18:2 | 34.37 | 3.90 | 39.14 | 12.61 | 94.40 | 20.04 | 94.16 | 32.94 | 23.86 | 6.27 | 27.87 | 7.75 | 46.54 | 28.19 | 30.00 | 24.20 |
| PE 18:1_20:1 | 0.39 | 0.20 | 0.89 | 0.61 | 0.61 | 0.28 | 0.63 | 0.24 | 0.52 | 0.33 | 0.47 | 0.21 | 0.62 | 0.63 | 0.32 | 0.18 |
| PE 18:1_20:2 | 0.48 | 0.24 | 0.57 | 0.46 | 1.03 | 0.47 | 1.11 | 0.82 | 0.22 | 0.05 | 0.32 | 0.16 | 0.46 | 0.16 | 0.90 | 0.01 |

| | | | | | | | | | | | | | | | | |
|---|---|---|---|---|---|---|---|---|---|---|---|---|---|---|---|---|
| PE 18:1_20:4 | 1.45 | 0.65 | 3.03 | 1.42 | 9.56 | 3.13 | 9.95 | 6.66 | 0.67 | 0.35 | 1.33 | 1.08 | 2.42 | 1.78 | 1.32 | 1.28 |
| PE 18:2_18:2 | 6.45 | 1.74 | 7.52 | 3.17 | 32.83 | 11.84 | 36.30 | 23.63 | 3.12 | 1.00 | 4.15 | 1.23 | 9.31 | 6.23 | 5.49 | 5.78 |
| PE 18:2_20:1 | 1.74 | 0.19 | 1.37 | 0.97 | 4.32 | 1.20 | 4.48 | 2.34 | 1.21 | 0.67 | 1.24 | 0.20 | 1.92 | 1.38 | 1.31 | 1.44 |
| PE 18:2_20:2 | 1.23 | 0.68 | 1.59 | 0.65 | 5.20 | 2.15 | 7.69 | 5.66 | 0.58 | 0.52 | 0.62 | 0.12 | 1.17 | 1.15 | 1.13 | 1.05 |
| PE total | 298.64 | 35.13 | 337.07 | 65.34 | 603.42 | 127.94 | 658.48 | 200.31 | 227.73 | 38.91 | 262.93 | 40.55 | 344.74 | 158.41 | 255.23 | 146.12 |
| PE P-16:0/16:0 | 0.16 | 0.14 | 0.15 | 0.03 | 0.24 | 0.13 | 0.23 | 0.16 | 0.11 | 0.06 | 0.11 | 0.05 | 0.17 | 0.13 | 0.13 | 0.04 |
| PE P-16:0/18:1 | 4.11 | 1.15 | 4.82 | 1.35 | 5.72 | 2.00 | 6.73 | 2.20 | 4.36 | 1.03 | 3.91 | 1.28 | 3.91 | 1.53 | 3.69 | 0.96 |
| PE P-16:0/18:2 | 13.22 | 3.22 | 14.44 | 4.63 | 33.03 | 14.88 | 36.25 | 17.78 | 11.83 | 3.48 | 13.07 | 2.26 | 14.25 | 7.00 | 11.26 | 7.64 |
| PE P-16:0/18:3 | 0.54 | 0.28 | 0.79 | 0.57 | 2.49 | 1.76 | 3.15 | 2.66 | 0.21 | 0.22 | 0.33 | 0.26 | 0.48 | 0.29 | 0.39 | 0.37 |
| PE P-16:0/20:3 | 0.69 | 0.24 | 0.51 | 0.21 | 1.14 | 0.62 | 1.32 | 0.52 | 0.48 | 0.31 | 0.57 | 0.36 | 0.45 | 0.17 | 0.45 | 0.11 |
| PE P-16:0/20:4 | 8.76 | 3.03 | 8.42 | 3.61 | 16.31 | 6.05 | 19.13 | 8.98 | 8.85 | 2.64 | 9.31 | 1.46 | 8.43 | 3.32 | 7.19 | 0.87 |
| PE P-16:0/22:4 | 1.30 | 0.36 | 1.63 | 0.77 | 2.52 | 1.20 | 3.28 | 1.30 | 1.40 | 0.16 | 1.69 | 0.27 | 1.40 | 0.89 | 1.63 | 0.32 |
| PE P-16:0/22:5 | 1.76 | 0.85 | 2.05 | 1.06 | 3.75 | 1.47 | 4.45 | 2.36 | 1.88 | 0.41 | 1.28 | 0.11 | 1.89 | 0.55 | 1.94 | 0.40 |
| PE P-16:1/18:2 | 0.15 | 0.08 | 0.33 | 0.14 | 0.79 | 0.63 | 1.15 | 0.90 | 0.10 | 0.08 | 0.04 | 0.02 | 0.43 | 0.19 | 0.36 | 0.15 |
| PE P-16:1/20:2 | 0.37 | 0.17 | 0.36 | 0.14 | 0.55 | 0.38 | 0.73 | 0.29 | 0.35 | 0.15 | 0.30 | 0.20 | 0.40 | 0.17 | 0.39 | 0.17 |
| PE P-16:1/22:2 | 0.26 | 0.12 | 0.32 | 0.23 | 0.42 | 0.28 | 0.50 | 0.38 | 0.38 | 0.25 | 0.20 | 0.18 | 0.17 | 0.14 | 0.30 | 0.19 |
| PE P-16:1/22:3 | 0.09 | 0.08 | 0.18 | 0.06 | 0.37 | 0.29 | 0.40 | 0.22 | 0.18 | 0.04 | 0.17 | 0.16 | 0.14 | 0.05 | 0.20 | 0.03 |
| PE P-18:0/16:0 | 0.16 | 0.14 | 0.10 | 0.05 | 0.16 | 0.02 | 0.13 | 0.07 | 0.10 | 0.03 | 0.12 | 0.11 | 0.13 | 0.09 | 0.11 | n.d. |
| PE P-18:0/18:1 | 3.04 | 1.06 | 3.21 | 0.77 | 4.04 | 1.70 | 4.56 | 1.35 | 2.28 | 0.68 | 2.71 | 0.38 | 2.92 | 1.27 | 2.53 | 0.51 |
| PE P-18:0/18:2 | 14.73 | 4.17 | 16.31 | 3.91 | 33.35 | 11.65 | 35.90 | 13.79 | 12.11 | 3.41 | 13.66 | 2.61 | 17.46 | 8.13 | 12.97 | 7.99 |
| PE P-18:0/18:3 | 0.41 | 0.15 | 0.48 | 0.20 | 2.07 | 0.85 | 2.42 | 1.64 | 0.14 | 0.06 | 0.32 | 0.20 | 0.31 | 0.20 | 0.46 | 0.51 |
| PE P-18:0/20:3 | 0.72 | 0.46 | 0.47 | 0.46 | 0.66 | 0.26 | 0.87 | 0.29 | 0.45 | 0.11 | 0.34 | 0.12 | 0.49 | 0.19 | 0.39 | 0.12 |
| PE P-18:0/20:4 | 7.21 | 2.68 | 7.42 | 2.69 | 11.46 | 4.94 | 14.06 | 4.98 | 7.69 | 1.09 | 8.35 | 1.01 | 7.35 | 2.20 | 7.63 | 1.24 |
| PE P-18:0/22:4 | 0.57 | 0.32 | 0.63 | 0.32 | 0.61 | 0.44 | 1.12 | 0.41 | 0.66 | 0.15 | 0.57 | 0.17 | 0.42 | 0.26 | 0.60 | 0.21 |
| PE P-18:0/22:5 | 0.75 | 0.42 | 0.80 | 0.24 | 1.60 | 0.59 | 1.49 | 0.36 | 0.70 | 0.22 | 0.79 | 0.31 | 0.69 | 0.15 | 0.99 | 0.41 |

| | | | | | | | | | | | | | | | | |
|---|---|---|---|---|---|---|---|---|---|---|---|---|---|---|---|---|
| PE P-18:1/16:0 | 0.72 | 0.40 | 0.96 | 0.42 | 1.01 | 0.44 | 2.00 | 0.64 | 0.78 | 0.22 | 1.28 | 0.49 | 1.01 | 0.54 | 1.39 | 0.43 |
| PE P-18:1/18:1 | 1.98 | 0.81 | 2.44 | 0.79 | 3.77 | 1.97 | 5.67 | 2.17 | 1.89 | 0.55 | 2.32 | 0.57 | 2.54 | 1.24 | 2.25 | 1.39 |
| PE P-18:1/18:2 | 6.19 | 1.61 | 8.53 | 2.61 | 25.05 | 13.51 | 30.80 | 15.17 | 5.78 | 1.70 | 7.57 | 2.97 | 9.14 | 5.54 | 8.39 | 6.85 |
| PE P-18:1/20:3 | 0.29 | 0.14 | 0.44 | 0.25 | 0.65 | 0.25 | 0.71 | 0.47 | 0.32 | 0.19 | 0.38 | 0.11 | 0.23 | 0.15 | 0.14 | 0.10 |
| PE P-18:1/20:4 | 5.07 | 1.57 | 5.07 | 1.93 | 8.69 | 3.23 | 11.61 | 6.22 | 4.97 | 0.72 | 6.25 | 1.48 | 4.60 | 2.45 | 5.18 | 1.60 |
| PE P-18:1/22:4 | 0.10 | 0.06 | 0.17 | 0.12 | 0.25 | 0.14 | 0.36 | 0.23 | 0.11 | 0.09 | 0.48 | 0.06 | 0.13 | 0.09 | 0.16 | 0.10 |
| PE P-18:1/22:5 | 0.15 | 0.11 | 0.14 | 0.10 | 0.40 | 0.22 | 0.49 | 0.41 | 0.10 | 0.07 | 0.23 | 0.03 | 0.10 | 0.05 | 0.15 | 0.12 |
| PE P-18:2/18:1 | 0.13 | 0.09 | 0.25 | 0.17 | 0.60 | 0.35 | 0.71 | 0.63 | 0.09 | 0.03 | 0.14 | 0.08 | 0.22 | 0.15 | 0.30 | 0.04 |
| PE P-18:2/18:2 | 0.84 | 0.39 | 0.80 | 0.24 | 4.30 | 2.79 | 4.16 | 2.85 | 0.47 | 0.18 | 0.75 | 0.57 | 1.04 | 0.73 | 0.86 | 0.78 |
| PE P-18:2/20:4 | 0.64 | 0.31 | 0.64 | 0.45 | 1.07 | 0.39 | 2.03 | 1.57 | 0.49 | 0.10 | 0.42 | 0.23 | 0.39 | 0.33 | 0.33 | 0.13 |
| EtherPE total | 75.04 | 17.72 | 82.67 | 24.46 | 166.87 | 67.55 | 196.40 | 87.18 | 69.00 | 14.15 | 77.13 | 15.06 | 81.00 | 34.70 | 71.99 | 27.14 |
| PI 16:0_18:1 | 28.31 | 9.83 | 50.44 | 20.26 | 27.50 | 12.94 | 42.37 | 18.14 | 20.74 | 11.12 | 27.85 | 12.06 | 29.55 | 16.91 | 43.19 | 17.54 |
| PI 16:0_18:2 | 100.10 | 39.76 | 217.69 | 76.80 | 204.22 | 44.91 | 252.36 | 101.23 | 77.05 | 32.24 | 106.92 | 15.30 | 132.10 | 60.08 | 121.65 | 122.58 |
| PI 18:0_18:2 | 191.52 | 59.48 | 285.38 | 116.90 | 303.27 | 48.54 | 418.00 | 163.46 | 139.78 | 40.80 | 176.15 | 57.15 | 232.54 | 121.60 | 220.25 | 185.23 |
| PI 18:0_20:4 | 100.01 | 39.49 | 108.27 | 47.46 | 173.86 | 43.41 | 180.37 | 104.64 | 62.35 | 20.92 | 82.99 | 22.32 | 95.23 | 42.27 | 89.57 | 20.77 |
| PI total | 419.95 | 97.97 | 661.77 | 237.98 | 708.84 | 118.55 | 893.10 | 371.13 | 299.91 | 47.43 | 393.92 | 68.21 | 489.42 | 207.20 | 474.67 | 327.34 |
| PS 16:0_18:2 | 1.63 | 0.58 | 2.20 | 1.41 | 3.22 | 1.42 | 3.27 | 1.30 | 0.80 | 0.25 | 0.98 | 0.32 | 1.79 | 1.13 | 1.22 | 1.25 |
| PS 18:0_18:1 | 1.86 | 0.34 | 2.12 | 0.48 | 2.16 | 0.51 | 2.73 | 1.13 | 1.54 | 0.48 | 1.44 | 0.25 | 2.06 | 1.07 | 1.40 | 0.19 |
| PS 18:0_18:2 | 23.35 | 1.86 | 26.99 | 7.04 | 40.81 | 7.29 | 45.11 | 12.90 | 16.76 | 3.81 | 20.00 | 4.46 | 25.49 | 11.85 | 18.66 | 11.24 |
| PS 18:0_20:4 | 1.39 | 0.18 | 1.64 | 0.44 | 5.07 | 1.49 | 4.96 | 2.62 | 0.61 | 0.18 | 0.79 | 0.29 | 1.59 | 0.89 | 0.95 | 0.81 |
| PS 18:1_18:2 | 2.63 | 0.72 | 3.39 | 1.03 | 7.36 | 1.67 | 7.77 | 3.20 | 1.45 | 0.56 | 2.09 | 0.84 | 3.41 | 2.24 | 2.35 | 2.23 |
| PS total | 30.86 | 2.84 | 36.34 | 9.95 | 58.63 | 10.76 | 63.83 | 20.58 | 21.15 | 4.54 | 25.30 | 5.66 | 34.34 | 16.91 | 24.58 | 15.61 |
| SM 18:1;O2/14:0 | 0.15 | 0.03 | 0.16 | 0.01 | 0.12 | 0.02 | 0.13 | 0.01 | 0.14 | 0.02 | 0.15 | 0.01 | 0.13 | 0.02 | 0.16 | 0.04 |
| SM 18:1;O2/16:0 | 16.83 | 2.15 | 17.71 | 1.10 | 15.57 | 2.07 | 16.71 | 0.67 | 15.41 | 1.53 | 16.74 | 1.29 | 16.37 | 0.76 | 16.97 | 1.96 |
| SM 18:1;O2/16:1 | 0.53 | 0.09 | 0.60 | 0.05 | 0.47 | 0.07 | 0.50 | 0.03 | 0.42 | 0.06 | 0.48 | 0.03 | 0.47 | 0.04 | 0.47 | 0.04 |

| | | | | | | | | | | | | | | | | |
|---|---|---|---|---|---|---|---|---|---|---|---|---|---|---|---|---|
| SM 18:1;O2/18:0 | 2.04 | 0.40 | 1.96 | 0.18 | 2.02 | 0.36 | 2.00 | 0.18 | 1.93 | 0.30 | 1.91 | 0.12 | 2.07 | 0.38 | 1.95 | 0.15 |
| SM 18:1;O2/18:1 | 0.53 | 0.08 | 0.62 | 0.02 | 0.48 | 0.08 | 0.56 | 0.02 | 0.44 | 0.07 | 0.52 | 0.04 | 0.47 | 0.07 | 0.49 | 0.05 |
| SM 18:1;O2/18:2 | 0.00 | 0.00 | 0.01 | 0.00 | 0.00 | 0.00 | 0.00 | 0.00 | 0.00 | 0.00 | 0.00 | 0.00 | 0.00 | 0.00 | 0.00 | 0.00 |
| SM 18:1;O2/18:4 | 0.01 | 0.00 | 0.01 | 0.00 | 0.01 | 0.00 | 0.01 | 0.00 | 0.01 | 0.00 | 0.01 | 0.00 | 0.01 | 0.00 | 0.01 | 0.00 |
| SM 18:1;O2/20:0 | 0.56 | 0.12 | 0.47 | 0.08 | 0.56 | 0.13 | 0.52 | 0.05 | 0.56 | 0.16 | 0.51 | 0.06 | 0.59 | 0.18 | 0.47 | 0.06 |
| SM 18:1;O2/20:1 | 0.19 | 0.04 | 0.23 | 0.01 | 0.21 | 0.04 | 0.26 | 0.01 | 0.18 | 0.04 | 0.22 | 0.02 | 0.19 | 0.05 | 0.20 | 0.04 |
| SM 18:1;O2/20:2 | 0.01 | 0.00 | 0.01 | 0.00 | 0.01 | 0.00 | 0.01 | 0.00 | 0.01 | 0.00 | 0.01 | 0.00 | 0.01 | 0.00 | 0.01 | 0.00 |
| SM 18:1;O2/20:5 | 0.02 | 0.00 | 0.02 | 0.00 | 0.01 | 0.00 | 0.01 | 0.00 | 0.02 | 0.00 | 0.02 | 0.01 | 0.02 | 0.00 | 0.02 | 0.00 |
| SM 18:1;O2/22:0 | 2.03 | 0.37 | 1.77 | 0.23 | 1.94 | 0.38 | 1.94 | 0.23 | 2.02 | 0.47 | 1.96 | 0.26 | 2.01 | 0.49 | 1.84 | 0.29 |
| SM 18:1;O2/22:1 | 0.48 | 0.09 | 0.53 | 0.13 | 0.51 | 0.12 | 0.58 | 0.08 | 0.46 | 0.13 | 0.52 | 0.12 | 0.52 | 0.16 | 0.49 | 0.10 |
| SM 18:1;O2/22:2 | 0.15 | 0.02 | 0.19 | 0.06 | 0.24 | 0.05 | 0.28 | 0.04 | 0.13 | 0.04 | 0.16 | 0.05 | 0.18 | 0.08 | 0.15 | 0.07 |
| SM 18:1;O2/22:3 | 0.01 | 0.00 | 0.01 | 0.00 | 0.01 | 0.00 | 0.01 | 0.00 | 0.00 | 0.00 | 0.00 | 0.00 | 0.01 | 0.00 | 0.01 | 0.00 |
| SM total | 23.55 | 3.19 | 24.30 | 1.59 | 22.17 | 3.21 | 23.54 | 1.12 | 21.71 | 2.70 | 23.21 | 1.94 | 23.03 | 2.00 | 23.23 | 2.32 |
| Cer 18:1;O2/16:0 | 1.54 | 0.19 | 1.29 | 0.11 | 1.51 | 0.10 | 1.43 | 0.22 | 1.54 | 0.32 | 1.36 | 0.08 | 1.42 | 0.28 | 1.29 | 0.12 |
| Cer 18:1;O2/18:0 | 0.75 | 0.10 | 0.57 | 0.06 | 0.78 | 0.20 | 0.64 | 0.07 | 0.77 | 0.14 | 0.63 | 0.08 | 0.71 | 0.17 | 0.59 | 0.06 |
| Cer 18:1;O2/18:2 | 0.16 | 0.05 | 0.12 | 0.02 | 0.15 | 0.05 | 0.14 | 0.02 | 0.14 | 0.04 | 0.16 | 0.01 | 0.15 | 0.06 | 0.11 | 0.01 |
| Cer 18:1;O2/20:0 | 0.49 | 0.14 | 0.39 | 0.09 | 0.47 | 0.14 | 0.43 | 0.01 | 0.46 | 0.14 | 0.43 | 0.09 | 0.46 | 0.16 | 0.37 | 0.04 |
| Cer 18:1;O2/20:2 | 0.22 | 0.06 | 0.16 | 0.03 | 0.20 | 0.06 | 0.16 | 0.02 | 0.22 | 0.07 | 0.22 | 0.04 | 0.19 | 0.08 | 0.14 | 0.02 |
| Cer 18:1;O2/20:4 | 0.03 | 0.01 | 0.02 | 0.01 | 0.02 | 0.01 | 0.02 | 0.01 | 0.02 | 0.01 | 0.01 | 0.01 | 0.02 | 0.01 | 0.02 | 0.01 |
| Cer 18:1;O2/22:0 | 1.94 | 0.38 | 1.66 | 0.19 | 1.92 | 0.39 | 1.89 | 0.03 | 1.84 | 0.31 | 1.84 | 0.26 | 1.78 | 0.51 | 1.55 | 0.15 |
| Cer 18:1;O2/22:1 | 0.27 | 0.04 | 0.25 | 0.07 | 0.27 | 0.06 | 0.31 | 0.08 | 0.25 | 0.06 | 0.26 | 0.07 | 0.28 | 0.10 | 0.23 | 0.07 |
| Cer 18:1;O2/22:2 | 2.37 | 0.62 | 1.96 | 0.24 | 2.25 | 0.50 | 2.35 | 0.07 | 2.10 | 0.57 | 2.11 | 0.23 | 1.92 | 0.64 | 1.77 | 0.17 |
| Cer 18:1;O2/22:3 | 0.18 | 0.02 | 0.19 | 0.04 | 0.19 | 0.04 | 0.21 | 0.01 | 0.18 | 0.03 | 0.20 | 0.04 | 0.20 | 0.05 | 0.15 | 0.02 |
| Cer 18:1;O2/22:4 | 0.03 | 0.01 | 0.02 | 0.01 | 0.02 | 0.01 | 0.02 | 0.01 | 0.02 | 0.01 | 0.02 | 0.01 | 0.02 | 0.01 | 0.02 | 0.01 |
| Cer 18:1;O2/22:5 | 0.12 | 0.03 | 0.08 | 0.01 | 0.10 | 0.02 | 0.11 | 0.04 | 0.11 | 0.01 | 0.11 | 0.03 | 0.10 | 0.02 | 0.08 | 0.01 |

| | | | | | | | | | | | | | | | | |
|---|---|---|---|---|---|---|---|---|---|---|---|---|---|---|---|---|
| Cer 18:1;O2/22:6 | 0.02 | 0.01 | 0.02 | 0.01 | 0.06 | 0.01 | 0.06 | 0.02 | 0.03 | 0.01 | 0.02 | 0.01 | 0.05 | 0.02 | 0.04 | 0.03 |
| Cer total | 8.12 | 1.31 | 6.72 | 0.76 | 7.93 | 1.30 | 7.77 | 0.43 | 7.69 | 1.33 | 7.37 | 0.79 | 7.29 | 1.96 | 6.37 | 0.31 |
| CE 16:0 | 0.31 | 0.14 | 0.37 | 0.20 | 0.34 | 0.15 | 0.37 | 0.09 | 0.31 | 0.07 | 0.45 | 0.16 | 0.36 | 0.09 | 0.47 | 0.22 |
| CE 16:1 | 0.86 | 0.09 | 1.08 | 0.25 | 0.81 | 0.16 | 1.08 | 0.23 | 0.85 | 0.16 | 1.25 | 0.27 | 0.95 | 0.12 | 1.27 | 0.18 |
| CE 18:0 | 0.51 | 0.13 | 0.55 | 0.16 | 0.59 | 0.11 | 0.73 | 0.13 | 0.66 | 0.11 | 0.79 | 0.09 | 0.59 | 0.04 | 0.71 | 0.23 |
| CE 18:1 | 3.46 | 0.54 | 3.21 | 1.10 | 3.16 | 0.97 | 3.37 | 0.86 | 3.31 | 0.84 | 4.74 | 2.01 | 3.57 | 0.54 | 3.95 | 1.41 |
| CE 18:2 | 15.07 | 2.51 | 18.18 | 9.01 | 17.18 | 12.07 | 18.33 | 5.06 | 14.75 | 5.24 | 22.32 | 14.75 | 14.00 | 2.13 | 14.41 | 3.53 |
| CE 18:3 | 1.22 | 0.31 | 1.51 | 0.97 | 1.21 | 0.76 | 1.52 | 0.62 | 1.18 | 0.66 | 1.78 | 1.13 | 1.09 | 0.24 | 1.26 | 0.40 |
| CE 20:4 | 0.47 | 0.11 | 0.59 | 0.33 | 0.56 | 0.52 | 0.57 | 0.17 | 0.43 | 0.14 | 0.69 | 0.50 | 0.44 | 0.22 | 0.46 | 0.32 |
| CE total | 21.91 | 3.39 | 25.50 | 11.72 | 23.86 | 14.52 | 25.99 | 6.86 | 21.48 | 6.95 | 32.02 | 18.67 | 21.00 | 2.92 | 22.52 | 6.08 |
| MG 16:0 | 0.86 | 0.19 | 0.77 | 0.26 | 1.35 | 0.51 | 1.11 | 0.12 | 1.15 | 0.65 | 0.84 | 0.11 | 0.90 | 0.44 | 0.65 | 0.20 |
| MG 18:0 | 5.46 | 2.44 | 5.24 | 2.82 | 7.83 | 4.88 | 5.29 | 1.58 | 9.80 | 6.57 | 6.92 | 1.13 | 7.71 | 3.99 | 5.31 | 2.31 |
| MG total | 6.32 | 2.55 | 6.01 | 3.06 | 9.18 | 5.32 | 6.40 | 1.59 | 10.95 | 7.22 | 7.76 | 1.21 | 8.61 | 4.42 | 5.96 | 2.43 |
| DG 14:0_16:0 | 0.01 | 0.00 | 0.01 | 0.01 | 0.02 | 0.01 | 0.02 | 0.00 | 0.01 | 0.01 | 0.02 | 0.01 | 0.02 | 0.01 | 0.01 | 0.00 |
| DG 14:0_16:1 | 0.00 | 0.00 | 0.00 | 0.00 | 0.00 | 0.00 | 0.00 | 0.00 | 0.00 | 0.00 | 0.00 | 0.00 | 0.00 | 0.00 | 0.00 | 0.00 |
| DG 14:0_18:0 | 0.00 | 0.00 | 0.00 | 0.00 | 0.00 | 0.00 | 0.00 | 0.00 | 0.01 | 0.00 | 0.00 | 0.00 | 0.00 | 0.00 | 0.00 | 0.00 |
| DG 14:0_18:1 | 0.01 | 0.01 | 0.01 | 0.01 | 0.02 | 0.01 | 0.02 | 0.01 | 0.01 | 0.00 | 0.01 | 0.01 | 0.01 | 0.01 | 0.01 | 0.00 |
| DG 14:0_18:2 | 0.00 | 0.00 | 0.00 | 0.00 | 0.02 | 0.01 | 0.02 | 0.01 | 0.00 | 0.00 | 0.01 | 0.00 | 0.01 | 0.01 | 0.01 | 0.01 |
| DG 14:0_22:2 | 0.00 | 0.00 | 0.00 | 0.00 | 0.00 | 0.00 | 0.00 | 0.00 | 0.00 | 0.00 | 0.00 | 0.00 | 0.00 | 0.00 | 0.00 | 0.00 |
| DG 16:0_16:0 | 0.14 | 0.04 | 0.15 | 0.06 | 0.22 | 0.06 | 0.25 | 0.05 | 0.19 | 0.06 | 0.21 | 0.02 | 0.18 | 0.07 | 0.18 | 0.06 |
| DG 16:0_16:1 | 0.06 | 0.01 | 0.07 | 0.02 | 0.07 | 0.03 | 0.10 | 0.01 | 0.07 | 0.03 | 0.10 | 0.01 | 0.08 | 0.04 | 0.08 | 0.03 |
| DG 16:0_18:0 | 0.15 | 0.05 | 0.15 | 0.05 | 0.25 | 0.07 | 0.25 | 0.01 | 0.23 | 0.07 | 0.23 | 0.02 | 0.24 | 0.09 | 0.20 | 0.05 |
| DG 16:0_18:1 | 0.61 | 0.13 | 0.60 | 0.17 | 0.86 | 0.23 | 0.90 | 0.05 | 0.74 | 0.21 | 0.77 | 0.11 | 0.77 | 0.24 | 0.71 | 0.25 |
| DG 16:0_18:2 | 0.98 | 0.23 | 0.93 | 0.30 | 2.66 | 0.80 | 2.62 | 0.93 | 0.97 | 0.24 | 1.04 | 0.21 | 1.49 | 0.76 | 1.09 | 0.84 |
| DG 16:0_18:3 | 0.03 | 0.00 | 0.03 | 0.02 | 0.20 | 0.09 | 0.19 | 0.11 | 0.02 | 0.01 | 0.03 | 0.01 | 0.06 | 0.04 | 0.05 | 0.04 |

| | | | | | | | | | | | | | | | | |
|---|---|---|---|---|---|---|---|---|---|---|---|---|---|---|---|---|
| DG 16:0_20:1 | 0.01 | 0.01 | 0.01 | 0.00 | 0.02 | 0.01 | 0.02 | 0.01 | 0.02 | 0.01 | 0.01 | 0.00 | 0.02 | 0.01 | 0.01 | 0.00 |
| DG 16:0_20:2 | 0.02 | 0.01 | 0.02 | 0.01 | 0.08 | 0.03 | 0.08 | 0.03 | 0.03 | 0.01 | 0.03 | 0.01 | 0.04 | 0.03 | 0.03 | 0.03 |
| DG 16:0_20:3 | 0.02 | 0.00 | 0.02 | 0.01 | 0.11 | 0.05 | 0.12 | 0.07 | 0.01 | 0.01 | 0.02 | 0.01 | 0.04 | 0.02 | 0.03 | 0.02 |
| DG 16:0_20:4 | 0.08 | 0.02 | 0.09 | 0.03 | 0.51 | 0.20 | 0.48 | 0.27 | 0.06 | 0.02 | 0.08 | 0.02 | 0.15 | 0.08 | 0.10 | 0.08 |
| DG 16:0_22:1 | 0.00 | 0.00 | n.d. | n.d. | 0.00 | 0.00 | 0.00 | 0.00 | 0.00 | 0.00 | 0.00 | 0.00 | 0.00 | 0.00 | 0.00 | 0.00 |
| DG 16:0_22:4 | 0.01 | 0.00 | 0.01 | 0.00 | 0.08 | 0.03 | 0.08 | 0.05 | 0.00 | 0.00 | 0.01 | 0.00 | 0.02 | 0.01 | 0.01 | 0.01 |
| DG 16:0_22:5 | 0.00 | 0.00 | 0.00 | 0.00 | 0.06 | 0.03 | 0.07 | 0.04 | 0.00 | 0.00 | 0.00 | 0.00 | 0.01 | 0.01 | 0.01 | 0.01 |
| DG 16:1_16:1 | 0.00 | 0.00 | 0.00 | 0.00 | 0.00 | 0.00 | 0.01 | 0.00 | 0.00 | 0.00 | 0.00 | 0.00 | 0.00 | 0.00 | 0.00 | 0.00 |
| DG 16:1_18:0 | 0.03 | 0.01 | 0.03 | 0.01 | 0.04 | 0.01 | 0.05 | 0.00 | 0.04 | 0.01 | 0.05 | 0.01 | 0.04 | 0.02 | 0.05 | 0.01 |
| DG 16:1_18:1 | 0.03 | 0.01 | 0.04 | 0.01 | 0.05 | 0.03 | 0.06 | 0.01 | 0.04 | 0.01 | 0.04 | 0.01 | 0.04 | 0.02 | 0.05 | 0.02 |
| DG 16:1_18:2 | 0.02 | 0.01 | 0.03 | 0.01 | 0.07 | 0.03 | 0.10 | 0.04 | 0.02 | 0.01 | 0.03 | 0.01 | 0.04 | 0.03 | 0.04 | 0.04 |
| DG 18:0_18:0 | 0.05 | 0.01 | 0.03 | 0.01 | 0.06 | 0.02 | 0.05 | 0.01 | 0.05 | 0.02 | 0.05 | 0.01 | 0.07 | 0.02 | 0.04 | 0.01 |
| DG 18:0_18:1 | 0.75 | 0.16 | 0.56 | 0.13 | 1.07 | 0.32 | 0.99 | 0.11 | 0.95 | 0.26 | 0.93 | 0.18 | 1.07 | 0.36 | 0.81 | 0.15 |
| DG 18:0_18:2 | 1.76 | 0.28 | 1.65 | 0.48 | 4.54 | 0.98 | 4.37 | 1.38 | 1.91 | 0.58 | 2.10 | 0.29 | 2.74 | 1.39 | 1.97 | 1.40 |
| DG 18:0_18:3 | 0.03 | 0.01 | 0.03 | 0.01 | 0.21 | 0.09 | 0.20 | 0.10 | 0.03 | 0.01 | 0.03 | 0.01 | 0.07 | 0.05 | 0.04 | 0.04 |
| DG 18:0_20:1 | 0.02 | 0.01 | 0.01 | 0.01 | 0.03 | 0.01 | 0.03 | 0.01 | 0.02 | 0.01 | 0.02 | 0.01 | 0.02 | 0.01 | 0.02 | 0.01 |
| DG 18:0_20:2 | 0.12 | 0.04 | 0.10 | 0.04 | 0.35 | 0.11 | 0.35 | 0.11 | 0.13 | 0.06 | 0.13 | 0.02 | 0.20 | 0.13 | 0.14 | 0.12 |
| DG 18:0_20:3 | 0.05 | 0.02 | 0.05 | 0.01 | 0.19 | 0.06 | 0.19 | 0.07 | 0.04 | 0.01 | 0.04 | 0.01 | 0.08 | 0.04 | 0.05 | 0.03 |
| DG 18:0_20:4 | 0.20 | 0.08 | 0.21 | 0.04 | 0.89 | 0.23 | 0.87 | 0.47 | 0.16 | 0.03 | 0.20 | 0.02 | 0.32 | 0.13 | 0.29 | 0.07 |
| DG 18:0_22:4 | 0.01 | 0.00 | 0.00 | 0.00 | 0.05 | 0.02 | 0.05 | 0.04 | 0.01 | 0.01 | 0.01 | 0.00 | 0.01 | 0.01 | 0.01 | 0.01 |
| DG 18:0_22:5 | 0.00 | 0.00 | 0.00 | 0.00 | 0.03 | 0.01 | 0.03 | 0.02 | 0.00 | 0.00 | 0.00 | 0.00 | 0.01 | 0.00 | 0.00 | 0.01 |
| DG 18:1_18:1 | 0.20 | 0.04 | 0.19 | 0.06 | 0.33 | 0.07 | 0.35 | 0.07 | 0.26 | 0.10 | 0.28 | 0.03 | 0.28 | 0.11 | 0.25 | 0.09 |
| DG 18:1_18:2 | 0.47 | 0.12 | 0.46 | 0.14 | 1.68 | 0.48 | 1.60 | 0.62 | 0.47 | 0.15 | 0.57 | 0.09 | 0.84 | 0.48 | 0.60 | 0.41 |
| DG 18:1_18:3 | 0.01 | 0.00 | 0.01 | 0.01 | 0.09 | 0.05 | 0.08 | 0.05 | 0.01 | 0.00 | 0.01 | 0.00 | 0.03 | 0.02 | 0.01 | 0.01 |
| DG 18:1_20:0 | 0.00 | 0.00 | 0.00 | 0.00 | 0.00 | 0.00 | 0.00 | 0.00 | 0.00 | 0.00 | 0.00 | 0.00 | 0.00 | 0.00 | 0.00 | 0.00 |

| | | | | | | | | | | | | | | | | |
|---|---|---|---|---|---|---|---|---|---|---|---|---|---|---|---|---|
| DG 18:1_20:1 | 0.02 | 0.00 | 0.02 | 0.01 | 0.03 | 0.01 | 0.03 | 0.01 | 0.02 | 0.01 | 0.02 | 0.01 | 0.03 | 0.01 | 0.02 | 0.01 |
| DG 18:1_20:2 | 0.01 | 0.01 | 0.01 | 0.01 | 0.04 | 0.02 | 0.05 | 0.03 | 0.01 | 0.01 | 0.01 | 0.01 | 0.02 | 0.02 | 0.01 | 0.01 |
| DG 18:1_20:3 | 0.00 | 0.00 | 0.00 | 0.00 | 0.02 | 0.01 | 0.02 | 0.01 | 0.00 | 0.00 | 0.00 | 0.00 | 0.01 | 0.01 | 0.00 | 0.01 |
| DG 18:1_20:4 | 0.02 | 0.01 | 0.02 | 0.01 | 0.14 | 0.06 | 0.11 | 0.06 | 0.01 | 0.00 | 0.01 | 0.00 | 0.04 | 0.02 | 0.02 | 0.01 |
| DG 18:1_22:1 | 0.00 | 0.00 | 0.00 | 0.00 | 0.01 | 0.00 | 0.01 | 0.00 | 0.00 | 0.00 | 0.00 | 0.00 | 0.01 | 0.00 | 0.00 | 0.00 |
| DG 18:2_18:2 | 0.05 | 0.01 | 0.05 | 0.02 | 0.34 | 0.13 | 0.32 | 0.18 | 0.04 | 0.02 | 0.05 | 0.01 | 0.11 | 0.08 | 0.07 | 0.06 |
| DG 18:2_18:3 | 0.00 | 0.00 | 0.00 | 0.00 | 0.02 | 0.01 | 0.02 | 0.02 | 0.00 | 0.00 | 0.00 | 0.00 | 0.01 | 0.00 | 0.00 | 0.00 |
| DG 18:2_20:0 | 0.01 | 0.00 | 0.01 | 0.00 | 0.03 | 0.01 | 0.04 | 0.02 | 0.01 | 0.01 | 0.01 | 0.01 | 0.02 | 0.01 | 0.02 | 0.02 |
| DG 18:2_20:1 | 0.04 | 0.01 | 0.04 | 0.02 | 0.15 | 0.04 | 0.15 | 0.07 | 0.04 | 0.02 | 0.04 | 0.01 | 0.07 | 0.05 | 0.05 | 0.04 |
| DG 18:2_20:2 | 0.01 | 0.00 | 0.01 | 0.01 | 0.06 | 0.03 | 0.07 | 0.04 | 0.01 | 0.01 | 0.01 | 0.01 | 0.01 | 0.01 | 0.01 | 0.01 |
| DG 18:2_20:4 | 0.00 | 0.00 | 0.00 | 0.00 | 0.03 | 0.01 | 0.03 | 0.02 | 0.00 | 0.00 | 0.00 | 0.00 | 0.00 | 0.00 | 0.00 | 0.00 |
| DG 18:2_22:0 | 0.00 | 0.00 | 0.00 | 0.00 | 0.01 | 0.00 | 0.01 | 0.01 | 0.00 | 0.00 | 0.00 | 0.00 | 0.01 | 0.00 | 0.00 | 0.00 |
| DG 18:2_22:1 | 0.01 | 0.00 | 0.01 | 0.01 | 0.04 | 0.01 | 0.04 | 0.02 | 0.01 | 0.01 | 0.01 | 0.01 | 0.02 | 0.01 | 0.02 | 0.02 |
| DG total | 6.07 | 1.13 | 5.71 | 1.65 | 15.78 | 4.21 | 15.51 | 4.96 | 6.69 | 1.87 | 7.25 | 1.08 | 9.36 | 4.37 | 7.16 | 4.01 |

Lipidomics data were normalized based on red blood cell counts and the ion abundance of internal standards added before lipid extraction via the modified Bligh and Dyer method.

n.d.: not detected.
